# Paired multiomic profiling defines the healthy human synovium–infrapatellar fat pad stromal continuum

**DOI:** 10.64898/2026.11.13.765998

**Authors:** Hannah Swahn, Irene Lorenzo Gomez, Lisa Janssen, Chelsea Kenvisay, Merissa Olmer, Tristan Maerz, Rachel E. Miller, Anne-Marie Malfait, Dana E. Orange, Bella Mehta, Miguel Otero, Theodore J. Price, The RE-JOIN Consortium Investigators, Martin K. Lotz

## Abstract

The synovium and infrapatellar fat pad (IPFP) are anatomically contiguous tissues implicated in osteoarthritis, yet their cellular and molecular organization in the healthy human knee remains poorly defined. We combined bulk RNA sequencing with paired single-nucleus transcriptomic and chromatin accessibility profiling to establish a quantitative multiomic reference of healthy human synovium and IPFP. Analysis of 54,716 nuclei from six cadaveric donors identified 23 synovial and 20 IPFP cell populations and indicated that the two tissues share one transcriptionally continuous, largely quiescent stromal compartment. Stromal populations shared core transcriptional programs but differed primarily in extracellular matrix output, with transcripts of inhibitory regulators predominating over those of their functional opponents. TIMP3 exceeded MMP14 in 42 of 43 single-nucleus populations across both tissues and by 13-to 39-fold in bulk tissue. Chromatin accessibility primarily distinguished stromal lineages and resolved transcriptionally defined subtypes, separating synovial lining from sublining fibroblasts and adipocytes from fibroblasts. Fibroblast and adipocyte populations occupied a common transcriptional trajectory accompanied by declining expression of organelle-turnover programs and little proliferation, with no dividing adipocytes and fewer than 0.05% of fibroblasts in the cell cycle. Despite this shared organization, the IPFP exhibited greater inferred intercellular signaling, attributable in part to BMP5, which was not detected in synovium. These findings describe healthy synovium and IPFP as regionally specialized parts of a transcriptionally continuous stromal compartment and provide a quantitative reference for identifying cellular and regulatory changes in joint disease.

## Introduction

Osteoarthritis (OA) affected 595 million people in 2020, 7.6% of the world’s population, and the knee is the joint most often involved.^1^ It is now understood as a disease of the whole joint, in which cartilage is one of several affected tissues. The synovium becomes inflamed and fibrotic, the intracapsular fat becomes fibrotic and secretes inflammatory mediators, bone remodels, and these changes track with pain and with structural progression.^2^ Two of these tissues sit in direct contact inside the knee: the synovium, the membrane that lines the joint cavity and maintains the synovial fluid, and the infrapatellar fat pad (IPFP), an adipose body enclosed within the same capsule. Synovial inflammation seen on magnetic resonance imaging precedes cartilage loss and predicts it,^3,4^ and the IPFP of OA patients releases inflammatory and fibrotic mediators that act on neighboring joint tissues.^5–7^ Both tissues are innervated and contribute to knee pain,^8,9^ and are therefore potential therapeutic targets and potential sources of signals that act on other joint tissues.

The evaluation of OA-associated changes lacks a description of either tissue in health. Single-cell studies of the synovium have focused on disease. They have resolved the fibroblast subsets that separate inflammation from bone damage, the resident macrophages that form a barrier around the joint cavity, and the signaling that maintains the sublining fibroblast state,^10–15^ and in OA they have identified fibroblast states and cross-talk with cartilage.^16,17^ In nearly all of this work the comparison tissue comes from joints already undergoing replacement, or from a small number of donors included as controls for a disease comparison, whose healthy tissue was not analyzed in its own right. As a result, most reported disease-associated changes have not been measured against a true healthy reference, and important changes may have been missed. The IPFP has been studied even less, with only one human single-cell atlas covering both tissues, built from four healthy donors, with the two tissues collected as a single biospecimen and with gene expression only.^18^ A second study described the IPFP in greater depth, using single-nucleus RNA sequencing and spatial transcriptomics in 15 donors with knee OA and 6 healthy controls, but did not analyze tissue health as a biological state in its own right.^19^ Thus, the description of health as a state is lacking for the IPFP and the synovium.

Adipose tissue also imposes a technical requirement. Tissue dissociation loses mature adipocytes, which are large and fragile, so a resulting atlas describes the stromal fraction of fat and not the fat itself. Single-nucleus preparations recover mature adipocytes^20,21^ and also allow analysis of chromatin accessibility.^22,23^ Measuring both transcriptome and chromatin state in the same nucleus makes it possible to link cell state with its regulatory architecture.^23^ Neither tissue had been profiled in this way in health.

In this study we performed paired synovial-IPFP profiling leveraging tissues from young healthy donors with no history of OA. We conducted paired analyses at two levels: first, we performed whole-tissue bulk RNA sequencing (RNA-seq) of 14 synovium and 18 IPFP donors. Then, we conducted paired single-nucleus multiome measurement of gene expression and open chromatin in the same nuclei, in six donors who each supplied both tissues. We use the resulting references to ask three questions. First, what is the quantitative profile of tissue health: how does each population divide its transcriptional output among functional programs, and which member of each opposing gene pair predominates in each population? Second, how much of cell identity is encoded in accessible DNA, and at which boundaries? Third, how do the two tissues relate to one another? Specifically, do they contain the same cell types, how do they signal internally, and do their shared stromal populations lie on a continuum? These results establish a comprehensive quantitative definition of the synovium and IPFP in the healthy knee microenvironment, providing a joint view of these two tissues, which are usually studied separately, that can in turn inform the evaluation of disease-associated changes.

## Results

### Tissue-level analyses of the healthy synovium and IPFP

We first performed whole-tissue bulk RNA-seq on synovium retrieved from 14 donors (11 male, 3 female; mean age 31.9 ± 6.7 years) and IPFP from 18 donors (13 male, 5 female; mean age 31.3 ± 7.9 years) all without a history of knee trauma or OA (Fig. 1a). The healthy state of the knees was further ascertained by macroscopically scoring all cartilage areas in the knee and also by macroscopic and histological evaluation of synovium and IPFP. RNA libraries were sequenced to a mean of 14.9 and 14.5 million assigned reads. A gene was counted as detected when it reached 1 TPM in at least half of the donors, which gave 12,869 detected genes in synovium and 11,361 in the IPFP. Protein-coding genes accounted for 81.8% of the synovial and 89.6% of the IPFP library (Fig. 1b).

**Fig. 1.**
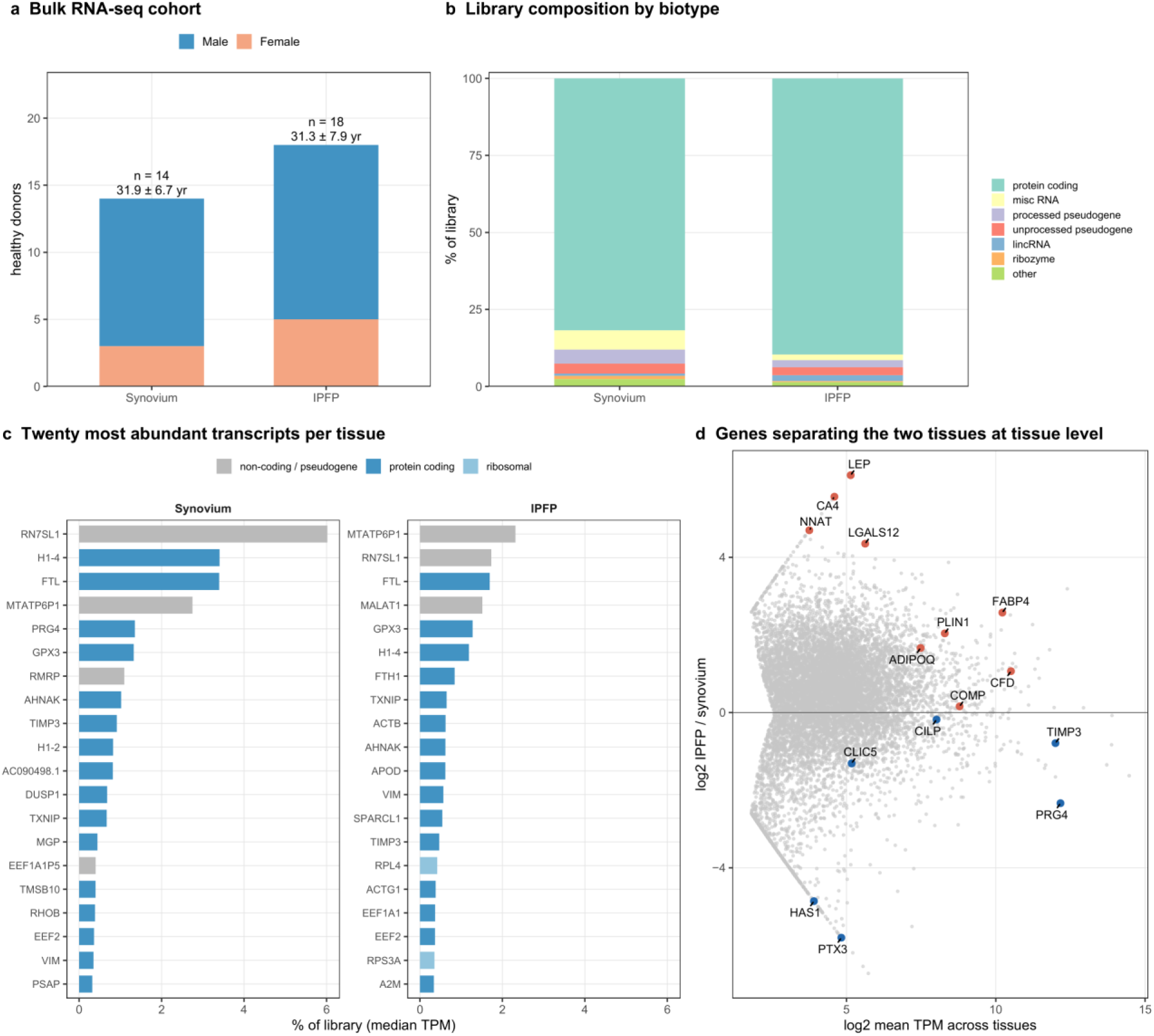
Tissue-level transcriptomes of healthy synovium and IPFP. Bulk RNA sequencing of whole tissue in a larger cohort than the single-nucleus multiome and of the same age range. **a** Donors by sex, with the number of donors and mean ± SD age above each bar. **b** Share of each library by gene biotype, as the sum of median TPM. **c** The twenty most abundant transcripts in each tissue by median TPM across donors, colored by class. Mitochondrial, ribosomal and pseudogene transcripts are retained because they are among the most abundant transcripts in bulk tissue. PRG4 is the fifth most abundant transcript in synovium. **d** Genes reaching 5 TPM in at least one tissue, plotted as tissue bias against mean abundance. Selected annotated genes are labeled; unannotated loci and clone identifiers, which dominate the extremes of the ranking, are plotted but not labeled.

The most abundant protein-coding transcripts (Fig. 1c) in synovium were H1-4, FTL, PRG4 (1.35% of the library, fifth among all transcripts), GPX3, AHNAK and TIMP3. In the IPFP they were FTL, GPX3, H1-4, FTH1, TXNIP, ACTB, AHNAK and APOD (with the long non-coding RNA MALAT1 ranking among them), with the adipocyte markers FABP4, CFD and PLIN1 within the top 200 (ranks 17, 18 and 109) and ADIPOQ at rank 253. Of 10,222 genes expressed at 5 TPM or above in at least one tissue, LEP was the most IPFP-restricted (68.5 TPM in IPFP against 0 in synovium, log2 fold change +6.1), with CA4, NNAT and LGALS12 among the next most restricted, while PRG4 was 5.0-fold higher in synovium (7,680 against 1,524 TPM) (Fig. 1d). The two tissues are therefore separable at tissue level based on their canonical lubricin and adipocyte markers.

The 200 most abundant protein-coding transcripts of the two tissues shared 138 genes and were enriched in both for epithelial–mesenchymal transition, coagulation, apoptosis, TNFα signaling via NF-κB and hypoxia; the transcripts in one tissue’s top 200 but not the other’s were enriched for matrix and circadian regulators in synovium (COL1A2, COL3A1, LUM, CILP, NR1D1, BHLHE40) and for adipogenesis in the IPFP (CD36, LPL, with CIDEC, PLIN1 and FASN among them) (Supplementary Fig. 1a, Supplementary Data 1). The most abundant transcripts of both tissues therefore share a stromal, vascular and stress-response program, with tissue-specific matrix and adipogenic additions.

### Cell populations of healthy human synovium

We performed single-nucleus multiome analysis of synovium from six healthy donors with an equivalent age range to donors analyzed using bulk RNA-seq (three male and three female; mean age 33.3 ± 11.1 years), retaining 33,932 nuclei for transcriptomic analysis (Supplementary Fig. 2a). We identified 23 populations across nine lineages (Fig. 2a,b, Supplementary Fig. 2b,c): seven fibroblast subtypes (PRG4 lining, PRG4+/CLIC5+; homeostatic, CXCL14+/APOD+; DPP4 sublining, DPP4+/SMOC2+; perivascular, LRP1B+/PDE1A+; ECM-remodeling sublining, COMP+/MFAP5+; perivascular sublining, PDZRN4+; and inflammatory lining, NFKBIA+/SLC39A14+), fibrocartilage (COL11A1+/MKX+), four endothelial populations (arterial, IGFBP3+/CLDN5+; venous, VWF+/ACKR1+; lymphatic, PROX1+/MMRN1+; and microvascular, BTNL9+/FLT1+), two mural populations (vascular smooth muscle, ACTA2+/LMOD1+; and a VSMC-like population, MYH11+/NTRK3+), three myeloid populations (inflammatory macrophages, MARCO+/MSR1+; resident macrophages, LYVE1+/F13A1+; and dendritic cells, HLA-DRB1+/DOCK8+), three lymphoid populations (plasma cells, IGKC+/TXNDC5+; T cells, IL7R+/SKAP1+; and B cells, MS4A1+/CD79A+), mast cells (KIT+/CPA3+), adipocytes (ADIPOQ+/PLIN1+) and nerve-sheath barrier cells (SLC22A3+/CNTN4+). Twenty-two of the 23 were present in all six donors and B cells in five; B cells (50 nuclei) and mast cells (60 nuclei) fall below the size downstream analyses require. Adipocytes formed a discrete synovial population (1.9% of nuclei), which dissociation-based atlases of synovium do not report^14,15^ because mature adipocytes are lost on dissociation and retained in nuclear preparations.^20,21^ Fibroblasts were the largest lineage at 44.7 ± 3.9% of nuclei, endothelium the most variable between donors (Supplementary Fig. 2b), and PRG4 lining fibroblasts the largest single population at 12.6%. Pseudobulk expression correlated with bulk RNA-seq of the same tissue (Spearman ρ = 0.51; Supplementary Fig. 3a,b). Genes detected by both platforms covered 99.4% of synovial and 99.8% of IPFP bulk RNA-seq TPM, and genes detected only in nuclei were mostly long non-coding and unannotated loci (Supplementary Fig. 1b–d, Supplementary Data 2).

**Fig. 2.**
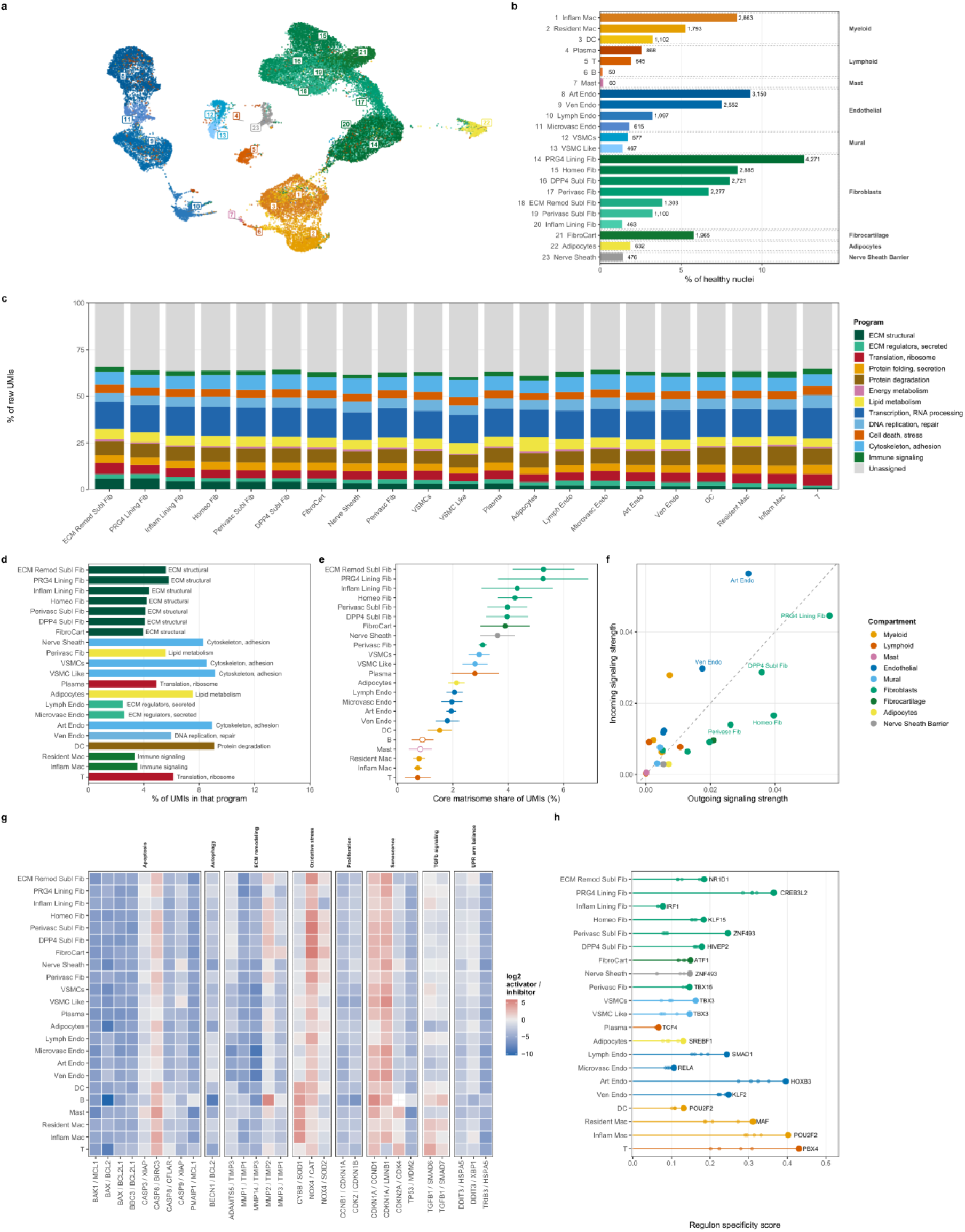
A single-nucleus reference for healthy human synovium. **a** UMAP of 33,932 nuclei from six healthy donors, 23 populations across nine lineages. Each population carries a number, keyed in (b); color is the lineage, with a light-to-dark step within it by population size. **B** Population abundance, percentage of nuclei, counts labeled, grouped into lineage blocks and ordered largest first within each. c Share of each population’s counts assigned to each of twelve programs from published gene ontology and MSigDB sets, each gene assigned to at most one program so the shares sum to 100%; 36.8% of counts are unassigned. **d** The program whose share is largest relative to the whole-tissue average. **e** Core matrisome share of counts; donor means ± SD, open points where fewer than 100 nuclei were pooled across donors. **f** Predicted communication (CellChat), outgoing against incoming strength; populations above the diagonal receive more signal than they send; the six strongest communicators are labeled. **g** Opposing gene pairs (Supplementary Data 3), as the per-donor ratio of one to the gene that blocks it; blue where the inhibitor dominates, red where the activator does. **h** The most specific regulon per population by regulon specificity score, from pySCENIC on healthy nuclei only; ranks two to five shown faintly. c, d, e, g and h share one cell-type order, descending by core matrisome share; b is ranked by abundance. c, d and h show the 21 populations meeting the abundance threshold, the other panels all 23. Label abbreviations: Fib, fibroblast; Subl, sublining; Homeo, homeostatic; Perivasc, perivascular; ECM Remod, ECM-remodeling; Inflam, inflammatory; FibroCart, fibrocartilage; Art, Ven, Lymph and Microvasc Endo, arterial, venous, lymphatic and microvascular endothelium; VSMC, vascular smooth muscle cell; Mac, macrophage; DC, dendritic cell; UPR, unfolded protein response.

### Extracellular matrix (ECM) transcripts distinguish synovial cell populations and dominate predicted cell–cell interactions

To define functions of the cell populations we divided the transcriptome into twelve programs defined by published gene ontology (GO) and MSigDB gene sets and measured each program’s share of sequencing counts (Fig. 2c). Transcription and RNA processing was the largest program in every population and near-constant at 15.17 ± 0.51%, whereas structural ECM varied 6.7-fold, from 0.87% in inflammatory macrophages to 5.80% in PRG4 lining fibroblasts (Fig. 2c,d, Supplementary Fig. 4a); the same analysis of the bulk RNA-seq TPMs ranked the programs similarly, although 8 of the 12 shares differed between platforms (Supplementary Fig. 3c). A scan of GO terms outside the twelve programs identified two additional signals: calcium handling in the mural cells (RYR2) and a metallothionein and zinc-stress response in the PRG4 lining and ECM-remodeling sublining fibroblasts (MT2A) (Supplementary Fig. 4b). Core matrisome transcripts^24^ followed the same pattern, contributing the largest percentages of UMIs in fibroblast and fibrocartilage populations (Fig. 2e, Supplementary Fig. 5a–c), and the strongest predicted signaling pathways were collagen and laminin (Supplementary Fig. 5d), with the fibroblast and endothelial lineages the strongest communicators (Fig. 2f). Synovial populations therefore differ little in general homeostatic machinery but substantially in ECM transcript output.

### Opposing gene pairs favor inhibitors of cell death, proliferation and ECM degradation

We next asked whether this output is accompanied by expression of genes that restrain cell death, proliferation and matrix degradation, measuring 29 pairs of genes whose products act in opposite directions on cell death, senescence, proliferation, the unfolded protein response, oxidative stress, autophagy, TGF-β signaling and ECM degradation (Supplementary Data 3) as the ratio in each donor of one gene to the gene that blocks it (Fig. 2g). Seven of the eight programs favored restraint and the eighth, senescence, was split: BCL2 exceeded BAX in all 23 populations by a median 34-fold, MMP14 was below TIMP3 in 22 of 23 by a median 33-fold (38.7-fold in bulk synovium), both cell-cycle pairs favored the inhibitor in 20 of 23, and the adaptive arm of the unfolded protein response exceeded the terminal arm in 37 of 69 comparisons, with the terminal arm favored in none. The activator was favored by CDKN1A over LMNB1 and CCND1, in 15 and 12 populations, and by NOX4 over catalase, in 14 of 23, although the other two oxidative stress pairs favored the inhibitor. Across both tissues, 24 of the 29 pairs favored the inhibitor (binomial p = 5 × 10⁻⁴ in each), the direction of 27 to 29 pairs was unchanged when any one donor was left out, and bulk RNA-seq gave the same direction for 24 of 29 pairs in synovium and 23 in the IPFP, although several activator genes, BAX among them, were near the detection floor in bulk. Nuclear counts favored long genes, so fold differences measured in nuclei were not quantitative, and NOX4 over catalase was not reproduced in bulk. Healthy synovium therefore expresses transcripts that favor restraint of cell death, division and matrix degradation.

### Synovial regulons resolve lineages, and the PRG4 lining is the most distinct fibroblast subtype

pySCENIC analysis of the healthy synovial nuclei identified 228 regulons across the 23 populations (Fig. 2h, Supplementary Fig. 6a), recovering known lineage factors: KLF2 in venous endothelium and MAF in resident macrophages ranked first, and LEF1 in T cells and PPARG in adipocytes ranked among the five most specific (Supplementary Fig. 6a). Fibroblast subtypes were the exception: regulon activity correlated at a median of 0.93 across the seven,, with AP-1 family members as having the highest regulon activity in sublining subsets. PRG4 lining fibroblasts were distinguished by regulon specificity, carrying the highest within-compartment regulon scores of any subtype (FOXC2 0.48, CREB3L2 0.48, SOX5 0.42; CUX1 ninth at 0.36) while still correlating with the other subtypes from 0.77 to 0.96 (Supplementary Fig. 6b,c). Transcriptional identity in synovium is thus specific at the lineage level. Among fibroblasts, the PRG4 lining subtype had the most specific regulons.

### snATAC-seq of the same nuclei resolves cell lineage strongly and subtype weakly

Single-nucleus ATAC-seq (snATAC-seq) analyses of the same nuclei revealed that chromatin in these tissues strongly marks lineage but only weakly resolves cellular subtypes (summarized in Supplementary Fig. 7). In the accessibility data of the same nuclei, 23,525 nuclei carried the transcriptomic labels by nuclear barcode and 21 populations met the thresholds used downstream (Methods) (Fig. 3a, Supplementary Fig. 8a–c, Supplementary Fig. 9a–d). Grouping the nuclei by chromatin alone reproduced the transcriptomic cell types only in part (adjusted Rand index 0.435, where 1 is an exact match), and when each nucleus was compared with its 30 most similar nuclei, 56% of them shared its cell type when similarity was measured by chromatin accessibility, against 82% when it was measured by RNA (Supplementary Fig. 8d,e). At matched cluster number, 11 of the 23 populations received a chromatin cluster of their own (Supplementary Fig. 8f,g). Of 69,646 peaks, 54,826 were enriched in at least one population and only 23.7% were exclusive to one (Fig. 3b). The fraction of peaks lying in a gene promoter (within 2 kb of a transcription start site) fell from 45.7% at exclusive peaks to 22.3% at widely shared ones (Fig. 3c, Supplementary Fig. 10a), and 14,844 peak–gene links joined 12,026 peaks to 3,226 genes (Supplementary Fig. 10b–d). In synovium, chromatin accessibility therefore distinguishes lineages well and subtypes poorly. Populations share a common pool of distal regulatory elements and differ by only a small set of population-specific elements (Supplementary Fig. 7a,b).

**Fig. 3.**
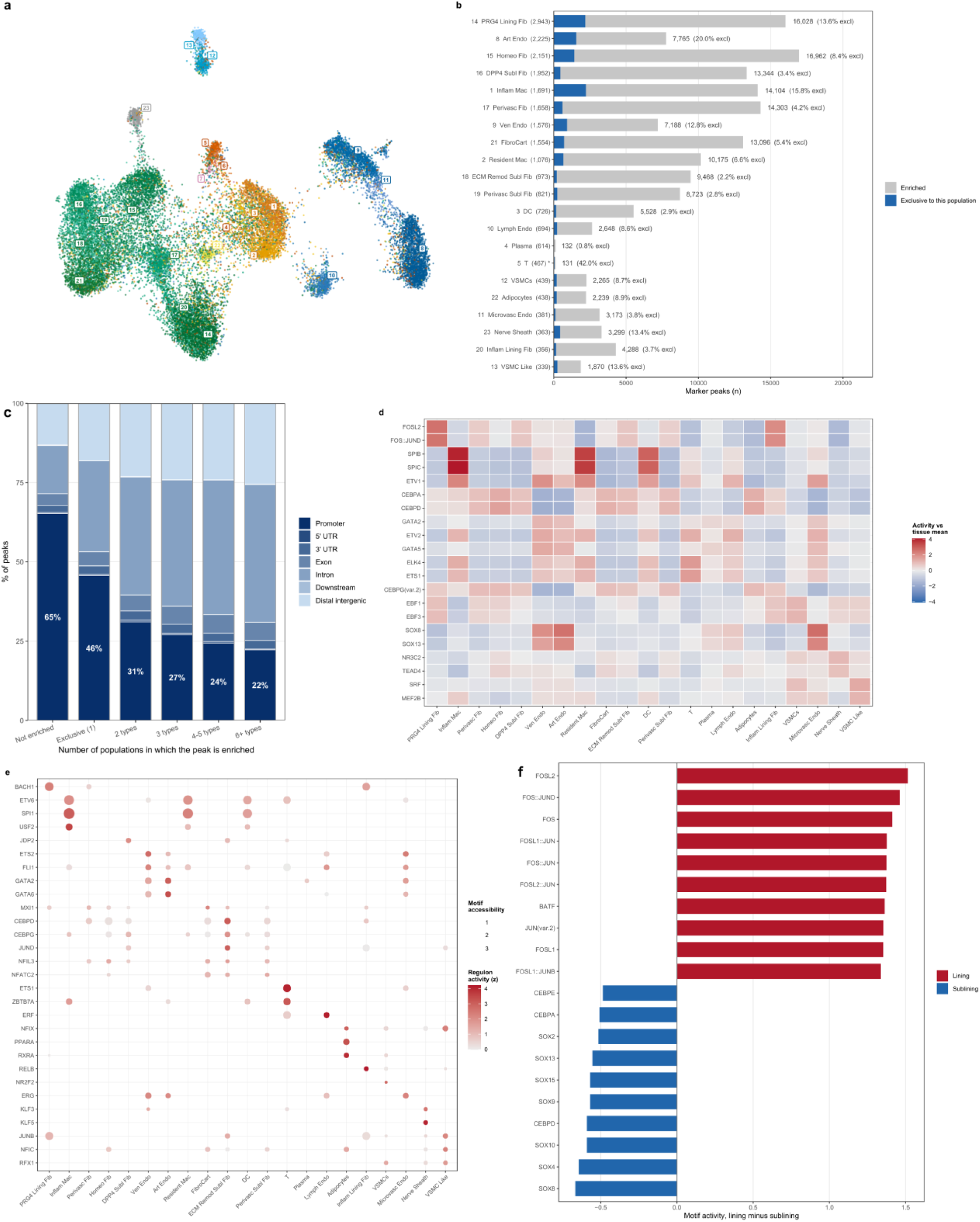
Chromatin accessibility in healthy human synovium. **a** Chromatin embedding of the 23,525 nuclei carrying a transcriptomic label, six donors, 23 populations, numbered and colored as in Fig. 2; the row labels of (b) carry the same numbers. **b** Marker peaks per population from donor-blocked pseudobulk (21 one-versus-rest contrasts): 54,826 of 69,646 peaks enriched in at least one population, 23.7% exclusive to one, drawn as the inner bar. Populations are ordered by nuclei, since marker counts track power; the asterisk marks T cells, whose test universe was reduced by independent filtering. **c** Genomic context of the same peaks by the number of populations in which each peak is enriched; promoter share labeled, falling from 46% at exclusive peaks to 22% at peaks shared by six or more. **d** chromVAR activity of the two most accessible motifs per population, against the tissue mean, donor-blocked. **e** Factors whose motif accessibility and regulon activity agree, from the RNA and ATAC measurements of the same nuclei; 366 factor-by-population calls carry both (median r = 0.26 matched, 0.02 shuffled). **f** The fibroblast compartment refitted alone: 421 of 633 motifs separate lining from sublining, ten strongest each way, AP-1 above and SOX and CEBP below. Fuller versions are Supplementary Figs. 8–12. Labels are abbreviated as in Fig. 2.

### Synovial transcription factor programs are shared within lineages and agree with the regulome in aggregate

We next asked which transcription factor motifs are accessible in each population, scoring 633 JASPAR CORE motifs in the same nuclei (Methods). The most accessible motifs differed between lineages and were largely shared by subtypes within a lineage (Fig. 3d, Supplementary Fig. 11a). CEBP-family motifs (CEBPD, then CEBPA) ranked first in five of the seven fibroblast subtypes and in fibrocartilage and adipocytes, SPI-family motifs (SPIC, SPIB) in all three myeloid populations, and SOX8 in three of the four endothelial subtypes, with motif profiles correlated within a lineage (r = 0.809–0.973) and less so across all population pairs (mean r = 0.369) (Supplementary Fig. 11b). Among the fibroblasts, the exception was the synovial lining, where both lining fibroblast subtypes ranked AP-1 motifs first (FOSL2 2.54 and FOS::JUND 2.51 in the PRG4 lining, 2.01 and 1.99 in the inflammatory lining). We then compared motif accessibility with regulon activity for the 144 factors that had both a motif and one of the 228 regulons. Three factors, all myeloid, agreed closely enough to pass multiple-testing correction individually: SPI1, TFEC and ETV6 (r = 0.92–0.96, padj = 0.048; Supplementary Fig. 11c). NFIA in the fibroblast and adipocyte group (r = 0.85) and GATA2 in the endothelium (r = 0.84) ranked next. Agreement between the two modalities was therefore evident across the factors as a whole (Supplementary Fig. 11c) but could be confirmed for only a few individual factors (Fig. 3e, Supplementary Fig. 11g).

We then asked whether these regulons are supported by open chromatin in the population where each is most specific (Supplementary Fig. 11d–g). The factor’s own motif was more accessible there for 66 of 144 factors, against 48 expected when motif profiles were shuffled; the peaks linked to the regulon’s target genes were more often accessible there than those of size-matched random target sets for 47 of 102 testable factors; and those elements carried the factor’s own motif more often than other factors’ motifs for 60 of 87. Twenty-one factors were supported by all three tests: eight in inflammatory macrophages, led by SPI1 and TFEC; nine across the endothelium, led by ERG and GATA2; and four in fibroblasts, led by FOSL2 in the PRG4 lining cells.

### A single regulatory boundary separates lining from sublining fibroblasts

Because chromatin did not distinguish the fibroblast subsets well, we refitted the peak and motif models within the fibroblast compartment alone. Lining and sublining fibroblasts differed at 421 of 633 motifs and 8,299 of 56,380 peaks (Fig. 3f, Supplementary Fig. 12a); all 14 top lining motifs were AP-1 family members, and the layer-defining peaks were depleted of promoters. Subtypes within a layer were not separable in the same way. Compared pairwise in the same model, the three sublining subtypes differed at no motif and the two lining subtypes at 17 of 633 (Supplementary Fig. 7c). When each subtype was compared only with other fibroblasts, PRG4 lining fibroblasts kept 42% of their enriched peaks, but the other six subtypes kept only 0.4–18% (Supplementary Fig. 12b). Most of what had distinguished those subtypes in the whole-tissue analysis was therefore shared with other fibroblasts. The two subtypes whose names do not indicate a layer, homeostatic and perivascular fibroblasts, both fell on the sublining side of a lining–sublining axis (Supplementary Fig. 12c). The lining–sublining division is therefore the only regulatory boundary resolved within the synovial fibroblast compartment, and one of the two boundaries most strongly resolved by chromatin in either tissue (Supplementary Fig. 7c).

### Cell populations of the healthy human IPFP

We applied the same transcriptomic analysis to IPFP from the same six donors, retaining 20,784 nuclei (Supplementary Fig. 13a). We identified 20 populations across seven lineages (Fig. 4a,b, Supplementary Fig. 13b,c): three adipocyte subtypes (secretory, CFD+/C1QTNF7+; lipogenic, GPAM+/DGAT2+; and a low-complexity population), six fibroblast subtypes (adhesion, ITGA11+/SULF1+; matrix-high, COMP+/CILP+; SMOC1+/GHR+; CXCL14+/IGFBP7+; lipid-laden, CIDEA+/CD36+; and a lining-like population, CLIC5+/HAS1+), four endothelial, two mural, two myeloid and two lymphoid populations, and mast cells. The low-complexity adipocyte population had a median of 1,235 detected genes per nucleus, against 2,866 and 3,234 in the other two, and its enriched genes were macrophage and endothelial transcripts (CD163, F13A1, PECAM1) and long nuclear-retained transcripts, with no adipocyte program. All 20 were detected in all six donors; six had at least 25 nuclei in fewer than four donors (the lining-like fibroblast, lymphatic endothelium, vascular smooth muscle, monocyte/dendritic cells, B/plasma cells and mast cells) and were excluded from analyses requiring donor-level replication. Fibroblasts made up 40.6% of the atlas, adipocytes 22.1% and endothelium 20.1% (Supplementary Fig. 13b). Pseudobulk expression correlated with bulk IPFP RNA-seq (Spearman ρ = 0.66; Supplementary Fig. 14a,b).

**Fig. 4.**
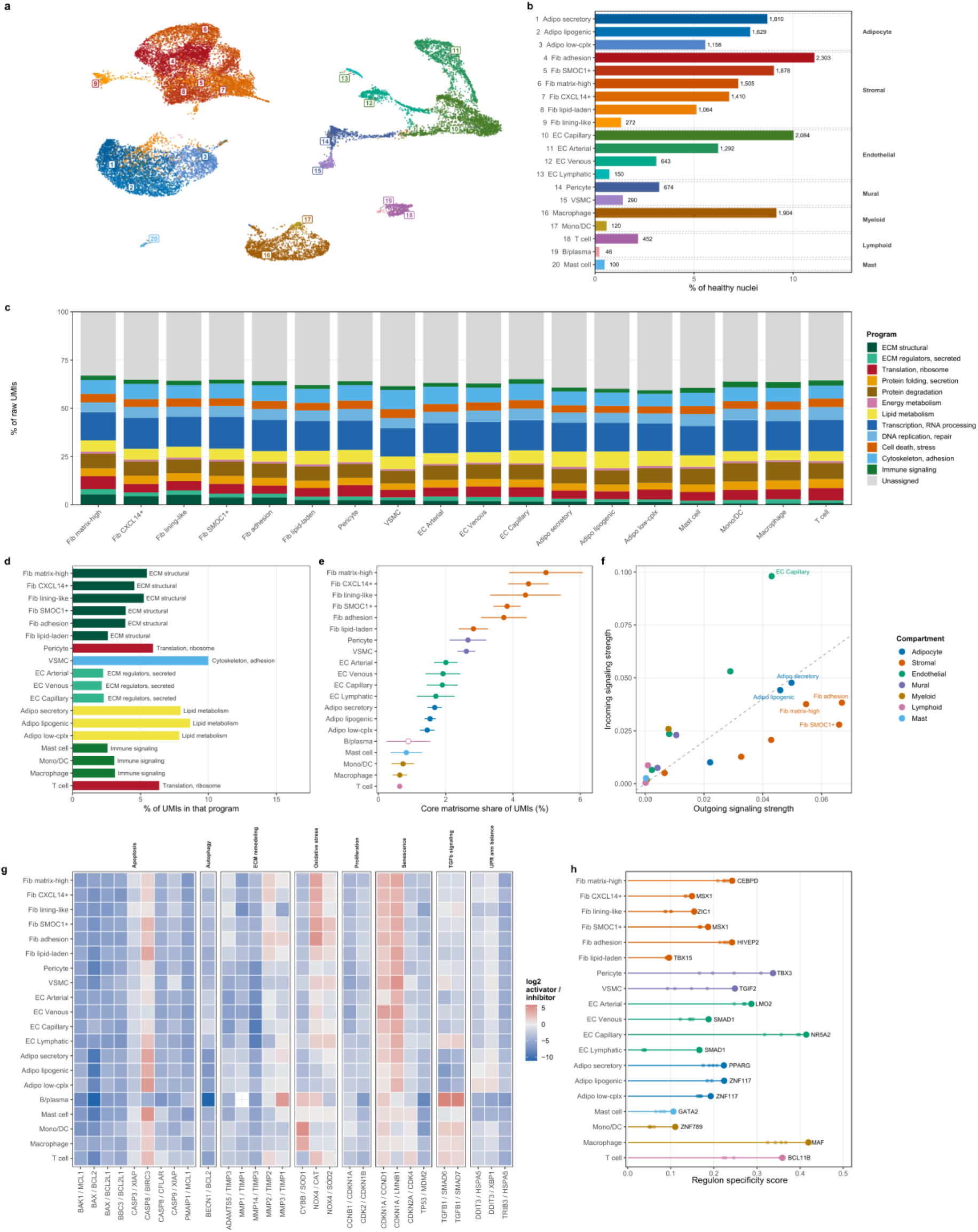
A single-nucleus reference for the healthy human IPFP. **a** UMAP of 20,784 nuclei from the same six donors, 20 populations across seven lineages. Each population carries a number, keyed in (b); color is the lineage, with a light-to-dark step within it by population size. **B** Population abundance, counts labeled, grouped into lineage blocks and ordered largest first within each. c Program shares as Fig. 2c; 36.8% unassigned. **d** The program with the largest share relative to the tissue average, per population (as Fig. 2d). **e** Core matrisome share of counts; donor means ± SD, open points where fewer than 100 nuclei were pooled. **f** Predicted communication, outgoing against incoming. **g** The opposing gene pairs of Fig. 2g. **h** The most specific regulon per population, IPFP-only pySCENIC run. Panel order as Fig. 2; c and d show the 18 populations meeting the abundance threshold, h the 19 meeting the regulon threshold, the rest all 20. Label abbreviations: Fib, fibroblast; Adipo, adipocyte; low-cplx, low-complexity; EC, endothelial cell; VSMC, vascular smooth muscle cell; Mono/DC, monocyte/dendritic cell; UPR, unfolded protein response.

### ECM transcripts distinguish IPFP populations, and the endothelium expresses secreted ECM regulators

Applied to the IPFP (Fig. 4c), the twelve-program partition used for synovium again made transcription and RNA processing the largest and most constant program (15.39 ± 0.65%, 1.17-fold range across the eighteen populations meeting the abundance threshold), with the same rank order in bulk RNA-seq, although 10 of the 12 shares differed between platforms (Supplementary Fig. 14c). Structural ECM varied 6.9-fold, from 0.79% in macrophages to 5.46% in the matrix-high fibroblast, and secreted ECM regulators 2.3-fold. Structural ECM was the most enriched program in all six fibroblast subtypes, lipid metabolism in all three adipocyte subtypes (8.66% of counts in the lipogenic subtype) and secreted ECM regulators in all three endothelial subtypes (Fig. 4d, Supplementary Fig. 15a). A GO scan outside the twelve added a metallothionein stress response in the matrix-high and lining-like fibroblasts (MT2A), calcium handling in vascular smooth muscle (RYR2) and cyclic nucleotide synthesis in pericytes (GUCY1A2) (Supplementary Fig. 15b). Core matrisome transcripts occupied 2.8–5.0% of counts in the fibroblast subtypes against 1.45–1.67% in the adipocyte subtypes (Fig. 4e, Supplementary Fig. 16a–c), and the strongest predicted interactions were again ECM–receptor pairs, sent mainly by the stroma (Fig. 4f, Supplementary Fig. 16d). In the IPFP, therefore, the stroma expresses most of the structural matrix transcripts, the endothelium expresses secreted ECM regulators, and the adipocytes express little ECM and a lipid program.

### Opposing gene pairs favor inhibitors of cell death and ECM degradation in the IPFP

We applied the same 29 opposing gene pairs tested in synovium (Fig. 2g) to the IPFP (Fig. 4g). Of 579 detected population-by-pair combinations, 339 favored the inhibitor by at least two-fold in every donor and 50 favored the activator. Six of the eight programs favored restraint; oxidative stress favored the activator in two of its three pairs, and senescence was split. Cell death and autophagy were the most one-sided programs, BCL2 exceeding BAX in 19 of the 20 populations by a median 138-fold and BECN1 in 18, and TIMP3 exceeded MMP14 in all 20 by a median 36.5-fold (12.9-fold in bulk IPFP). Senescence was the one program in which the activator side was common, at 22 of 80 combinations, and it was spread across the tissue, with 13 of the 20 populations having at least one activator-dominant senescence pair, in every case CDKN1A against LMNB1 or CCND1. As in synovium, transcripts in the healthy IPFP favored restraint of cell death and matrix degradation, although a senescence-associated signature was detectable.

### IPFP regulons separate the lineages, but the fibroblast subtypes largely share one regulatory program

pySCENIC applied to all 20 IPFP cellular populations returned 225 regulons (Fig. 4h, Supplementary Fig. 17a). The three adipocyte subtypes were most specifically marked by adipogenic factors (PPARG, TEAD1 and TCF7L2 in the secretory subtype; ZNF117, PPARG and CEBPA in the lipogenic one), and the other populations by lineage-defining factors, among them MAF, SPI1 and IRF8 in macrophages, BCL11B and TCF7 in T cells, GATA2 in mast cells, and SOX17 among the five most specific in capillary endothelium. The mesenchymal subtypes did not have separate regulatory programs: the two most similar population pairs in the tissue were the two adipocyte subtypes and the SMOC1+/GHR+ and CXCL14+/IGFBP7+ fibroblasts (both r = 0.993), all fifteen fibroblast pairs correlated at 0.87 or above (Supplementary Fig. 17b,c), and within the adipogenic axis silhouette widths ranged from −0.17 to 0.13. Regulon specificity was uneven across the fibroblast subtypes: 39 regulons were most specific to the adhesion subtype and 22 to the matrix-high one, against 3, 1 and 0 for the SMOC1+/GHR+, CXCL14+/IGFBP7+ and lipid-laden subtypes. The most specific regulon nonetheless differed between most subtypes (HIVEP2 in the adhesion, CEBPD in the matrix-high, ZIC1 in the lining-like and TBX15 in the lipid-laden subtype, and MSX1 in both the SMOC1+/GHR+ and CXCL14+/IGFBP7+ subtypes; Fig. 4h), at specificity scores of 0.10 to 0.24. The IPFP regulome therefore resolves the lineages and the adipocyte–fibroblast boundary, but the fibroblast subtypes largely share one program.

### snATAC-seq of the IPFP resolves lineage, and peak exclusivity depends on cluster resolution

In the IPFP-derived chromatin, 18,886 nuclei carried the transcriptomic labels and 14 populations met the testing thresholds (Fig. 5a, Supplementary Fig. 18a–c, Supplementary Fig. 19a–d). Chromatin-only clustering again reproduced the cell types only in part (adjusted Rand index 0.41), and when each nucleus was compared with its 30 most similar nuclei, 56% of them shared its cell type in chromatin, against 83% in RNA (Supplementary Fig. 18d,e). At matched cluster number, 12 of the 20 populations, including four of the six fibroblast subtypes, received a chromatin cluster of their own. Four clusters each held two populations: lipogenic with low-complexity adipocytes, SMOC1+/GHR+ with lipid-laden fibroblasts, venous with lymphatic endothelium, and T with B/plasma cells (Supplementary Fig. 18f,g). Of 99,697 peaks, 84,719 were enriched in at least one population and 21.0% were exclusive to one, against 52.6% when the test was applied at lower resolution, to 14 broader populations. Exclusive peaks were the more promoter-proximal set (39.1% against 21–23% for peaks enriched in three or more populations, in every quartile of peak coverage; Fig. 5b,c, Supplementary Fig. 20a), and 17,141 peak–gene links joined 14,231 peaks to 3,397 genes (Supplementary Fig. 20b–d). IPFP populations are therefore distinguished by a small set of population-specific, promoter-proximal elements, together with graded use of a larger shared set of distal elements.

**Fig. 5.**
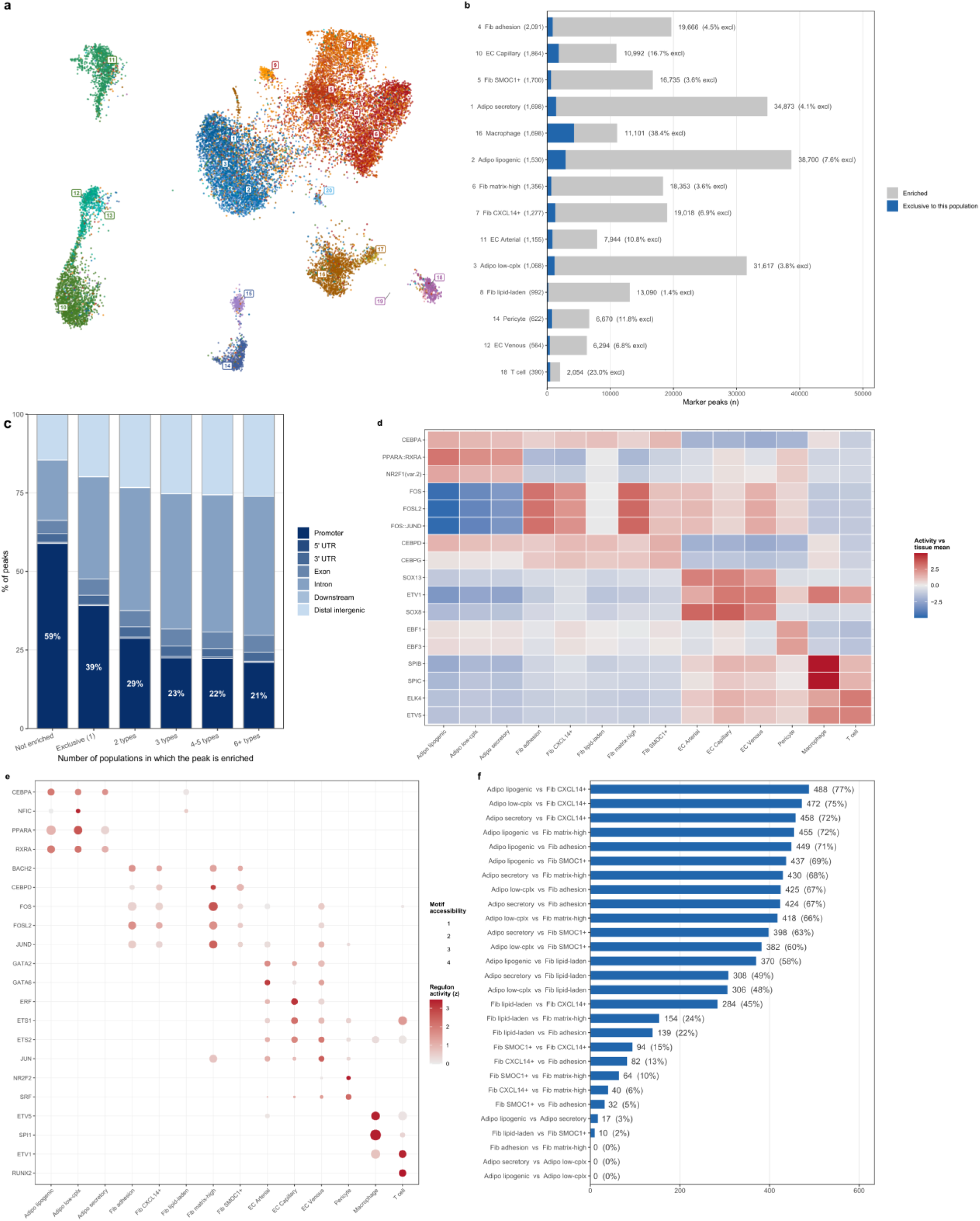
Chromatin accessibility in the healthy human IPFP. **a** Chromatin embedding of 18,886 labeled nuclei, 20 populations, numbered and colored as in Fig. 4; the row labels of (b) carry the same numbers. **b** Marker peaks per population (14 one-versus-rest contrasts): 84,719 of up to 99,697 peaks enriched, 21.0% exclusive to one population. The comparable figure at this reference’s earlier 14-population resolution was 52.6%, so exclusivity depends on cluster resolution as much as on the peaks themselves. **c** Genomic context by the number of populations in which each peak is enriched; promoter share falls from 39% at exclusive peaks to 21% at peaks shared by six or more, the same direction as synovium. **d** chromVAR activity of the top two motifs per population. **e** Factors whose motif and regulon agree: 338 factor-by-population calls across 116 factors and 14 populations (median r = 0.39 matched, 0.21 shuffled). **f** The adipogenic axis: bars give the number of motifs differing at adjusted p < 0.05 for each pair of populations (percentage of the 633 tested); separation follows the adipocyte– fibroblast boundary: all 15 pairs spanning it separate at 306 to 488 of 633 motifs, all 13 pairs within either compartment at 0 to 284, and the two axis endpoints at 455. The parent-label comparison is Supplementary Fig. 23. Fuller versions are Supplementary Figs. 18–22. Labels are abbreviated as in Fig. 4.

### AP-1, CEBP, SOX and SPI motifs mark the IPFP lineages, and many IPFP regulons are supported by accessible chromatin

As in synovium, the most accessible motifs differed mainly between lineages (Fig. 5d, Supplementary Fig. 21a). In all three adipocyte subtypes the motif with the largest accessibility difference from the other populations was that of the nuclear receptor heterodimer PPARA::RXRA, followed by NR2F1 and CEBPA. The fibroblast subtypes split into two signatures: the adhesion, CXCL14+/IGFBP7+ and matrix-high subtypes were led by AP-1 motifs (FOS::JUND, FOSL2, FOS, FOSL1::JUN), and the SMOC1+/GHR+ and lipid-laden subtypes by CEBP-family motifs (CEBPD, CEBPA, CEBPG, CEBPE). SOX8 led the arterial and capillary endothelium and ranked second, after ETV1, in venous endothelium, SPIB, SPIC, EHF and SPI1 the macrophages, ETS factors the T cells, and EBF3 and EBF1 the pericytes. Motif profiles were correlated within lineages (0.99 across fibroblast subtypes, 0.985 across endothelial subtypes, 0.876 across adipocyte subtypes) against a mean of 0.70 across all population pairs (Supplementary Fig. 21b). For the 140 factors carrying both a motif and a regulon, the median cross-modal correlation was 0.387 against 0.212 for randomly paired factors (Fig. 5e, Supplementary Fig. 21c). Across both tissues the family of the top-ranked motif largely followed lineage. The clearest split was within the fibroblasts, where AP-1 led the two synovial lining subtypes and three IPFP subtypes and CEBP the others; endothelial populations were led by SOX or ETS motifs, and the synovial mural and lymphoid populations also split between families (Supplementary Fig. 7d).

The three tests of whether a regulon is supported by accessible chromatin in its own population gave the same result in the IPFP (Supplementary Fig. 21d–g): the motif was more accessible in the regulon’s own population for 75 of 140 factors against 41.1% expected (p = 0.001), the peaks linked to its target genes were more accessible there than for random gene sets for 79 of 105, and those elements carried the factor’s own motif above other factors’ for 69 of 98 (paired Wilcoxon p = 1.3 × 10⁻⁵). Thirty-eight factors passed all three, recovering an expected regulator of every testable lineage, among them CEBPA in the lipogenic adipocyte, GATA2 in capillary endothelium, SPI1 in macrophages and NR2F2 in pericytes, with the fibroblast AP-1 factors split between the adhesion (ATF2, FOSL2) and matrix-high (FOS, FOSB, JUND) subtypes.

### Chromatin separates the adipocyte from the stroma, and the stromal subtypes from each other

We refitted the peak and motif models within the adipogenic axis of the three adipocyte and five testable fibroblast subtypes (the lining-like subtype has at least 25 nuclei in only three donors). Of 633 motifs, every adipocyte-versus-fibroblast contrast separated 306 to 488, consistent with a distinct adipocyte regulatory state, while the two biologically named adipocyte subtypes differed from each other by only 17 (Fig. 5f, Supplementary Fig. 22a). Among the fibroblasts the number of differing motifs was graded, from 284 motifs between the lipid-laden and CXCL14+/IGFBP7+ subtypes down to none at all between the adhesion and matrix-high subtypes. When the fibroblast subtypes were projected onto an axis defined by the lipogenic adipocyte and the matrix-high fibroblast (the populations with the highest lipid-metabolism and core-matrisome shares, respectively), they fell in order: matrix-high at the fibroblast pole (−0.996), then adhesion (−0.994), CXCL14+/IGFBP7+ (−0.954) and SMOC1+/GHR+ (−0.924), with the lipid-laden subtype alone well away from the pole at −0.587 (Supplementary Fig. 22c). When the contrast was restricted to the adipogenic axis, the subtypes retained different fractions of their whole-tissue enriched peaks: the adhesion, CXCL14+/IGFBP7+ and matrix-high subtypes retained 0.51–0.57 and the adipocytes 0.62–0.75, while the SMOC1+/GHR+ subtype retained 0.28 and the lipid-laden one almost none (Supplementary Fig. 22b). The fibroblast subtypes therefore occupy graded positions along the axis, and only the lipid-laden subtype lies appreciably toward the adipocyte end (Supplementary Fig. 22c). This subtype, however, received 31% of its nuclei as held-out predicted doublets (Methods), so part of its shift toward the adipocyte may reflect adipocyte–fibroblast doublets.

Before subclustering, the IPFP reference had only two stromal labels. Compared directly within the same nuclei, these two labels differed at only 2 of 633 motifs, whereas the subtypes obtained by re-clustering those nuclei differed by up to 266 motifs in a model fitted within the stromal arm (Supplementary Fig. 23). The subtypes therefore capture chromatin differences that the original two-way split averaged out. In the adipocyte arm, refitted within the arm, the two biologically named subtypes differed at 59 motifs (17 in the adipogenic-axis model of Fig. 5f, fitted across all testable axis states), whereas the low-complexity population differed from them at 0 and 1 motifs. This is consistent with a group of nuclei defined by low sequencing depth, with no regulatory state of its own (Supplementary Fig. 24). Because these nuclei are equally close to both named subtypes, we kept them under a separate, flagged label and excluded them from analyses they would distort (Methods); in the joint reference they merge with the other adipocytes.

### Synovium and IPFP share their major cell populations and signaling architecture

Analyzed together across the same six donors and all twelve libraries, the two tissues retained 54,716 nuclei (33,932 synovium, 20,784 IPFP) and resolved 16 populations (Fig. 6a,b, Supplementary Figs. 25–27). The joint object was clustered at a coarser resolution than either tissue alone, so related subtypes merged: the 14 fibroblast subtypes resolved as three joint fibroblast populations (sublining, homeostatic and lining), the three IPFP adipocyte subtypes as one, the myeloid populations as one and synovial B cells with IPFP B/plasma cells as one (synovial plasma cells formed no joint cluster; Supplementary Fig. 26), while endothelial and mural populations mapped one to one (Supplementary Data 4). Every population contained nuclei from both tissues and none exceeded 87% of a single tissue, but proportions differed: synovium was dominated by sublining fibroblasts (18.1% of its nuclei), with macrophages and lining fibroblasts at 15.6% each, and the IPFP by adipocytes (23.1%) and homeostatic fibroblasts (18.3%) (Fig. 6c). Nuclei from the two tissues occupied the same regions of the embedding, with the adipocyte and synovial lining clusters the most tissue-skewed (Supplementary Fig. 28a,b). The joint nerve-sheath barrier population was predominantly synovial (554 of 672 nuclei); its 118 IPFP nuclei, from all six donors, had been placed mostly (99) among the CXCL14+/IGFBP7+ fibroblasts by the IPFP reference and expressed the same perineurial markers (SLC22A3, SLC2A1, CLDN1, NGFR) at lower detection.

**Fig. 6.**
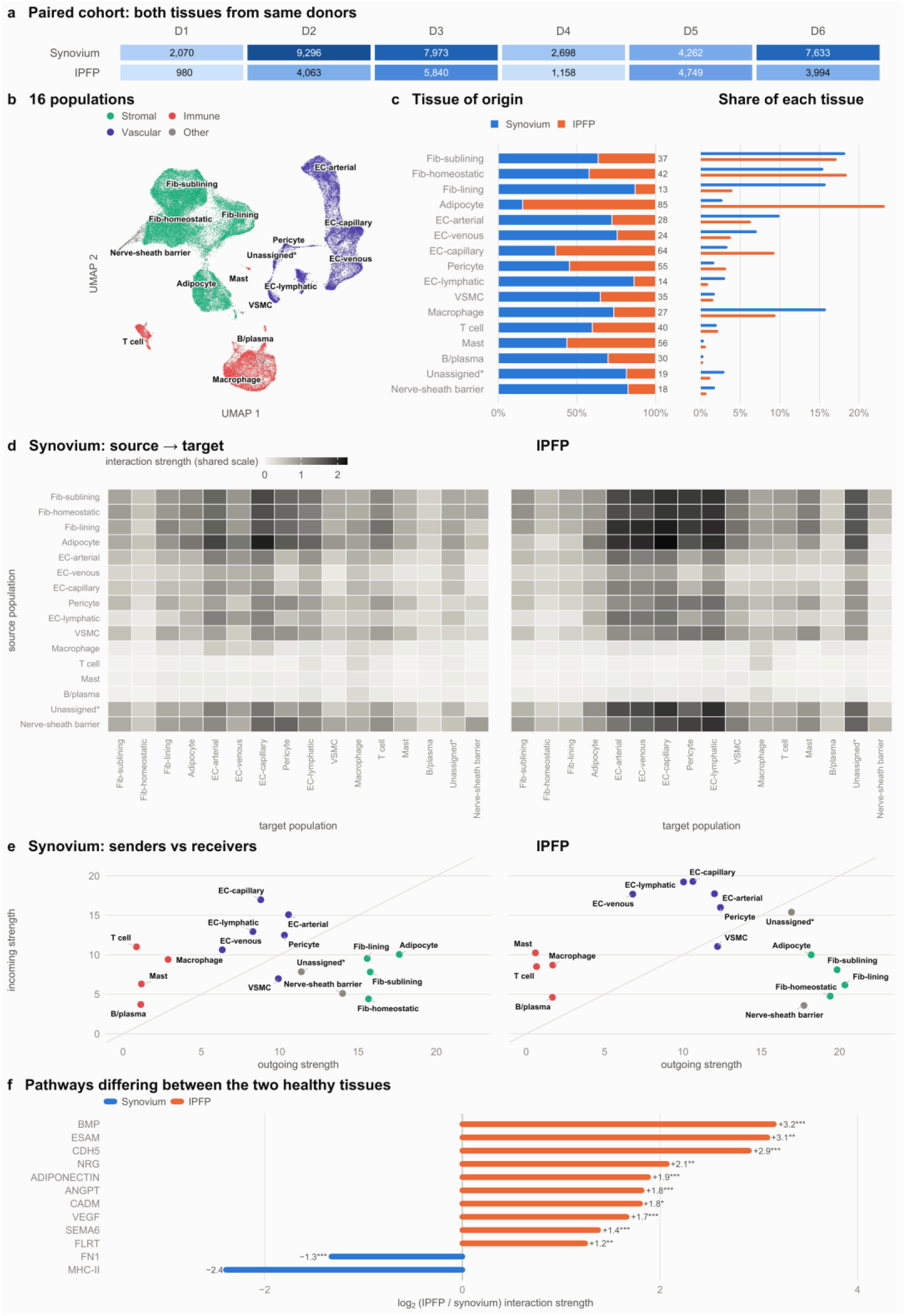
Cell–cell communication within healthy synovium and IPFP. **a** Nuclei per donor (D1–D6, as in Supplementary Figs. 2 and 13) and tissue after quality control; all six donors gave both tissues (from the same knee in five; Methods), 12 libraries and 54,716 nuclei (33,932 synovium, 20,784 IPFP). **b** UMAP of all 54,716 nuclei, colored by lineage family and labeled with the 16 populations annotated on this object (Methods); “Unassigned*” is a low-signal population that could not be annotated. **c** Composition in two normalizations. Left: tissue of origin per population, each bar 100%, IPFP percentage labeled; no population is tissue-exclusive, and they range from 13% IPFP (synovial lining fibroblasts) to 85% (adipocytes). Right: each tissue’s composition, each tissue 100%. **d** Communication inferred separately within each tissue over the same 16 populations; shading is summed interaction strength from sender (rows) to receiver (columns), one shared color scale (0–2.27). **e** Senders against receivers, from the row and column sums of d; shared axis limits. The top three IPFP senders (lining, sublining and homeostatic fibroblasts) are within 5% of each other and change order when doublets are removed, so no top IPFP sender is reported (Methods). **f** Pathways present in both tissues with strength ≥ 0.03, ranked by log2 fold change (IPFP over synovium), twelve largest shown; asterisks give CellChat’s permutation test over cell labels (one, two and three asterisks: p < 0.05, 0.01 and 0.001). Inference used no population-size term and was strictly within tissue.

Cell–cell communication inferred within each tissue gave both tissues the same architecture: fibroblast and adipocyte populations had the strongest outgoing signal, endothelial populations received most of it, and lymphoid populations contributed little in either direction (Fig. 6d,e). Adipocyte to capillary endothelium was the strongest sender–receiver pair in both synovium and IPFP (interaction strength 2.11 and 2.27), and adipocytes were the strongest synovial sender despite making up only 2.6% of synovial nuclei in the joint reference.

The IPFP had more inferred signaling than synovium on every global measure despite 39% fewer nuclei: 5,985 against 5,176 significant ligand–receptor pairs, total interaction strength 181.1 against 150.3, and 65 against 59 detected pathways. Among pathways detected in both tissues at strength 0.03 or more, BMP showed the largest shift toward the IPFP (log2 fold change +3.15), followed by ESAM, CDH5, NRG and ADIPONECTIN, and the endothelium-directed pathways ANGPT (+1.8) and VEGF (+1.7) were also IPFP biased. Among the twelve largest shifts, only MHC-II (−2.4) and FN1 (−1.3) were synovium biased (Fig. 6f). The higher BMP signaling in the IPFP was attributable to ligand expression: BMP5 was not detected in synovium and supplied 48% of the IPFP’s BMP signal, BMP6 was 4.6-fold stronger in the IPFP, and the receptor pairs were largely the same in both tissues (five of six shared). In the IPFP, capillary and arterial endothelium and the homeostatic and sublining fibroblasts were the main BMP senders, and in both tissues the adipocyte was the main receiver (Supplementary Fig. 29).

### Synovial and IPFP stromal cells lie on one continuous trajectory, with distinct regulators at its root, middle and end

A trajectory fitted over the 30,455 mesenchymal nuclei of the joint reference (sublining, homeostatic and lining fibroblasts and adipocytes), rooted at the population scoring highest for a published DPP4+ progenitor signature,^18^ returned a single lineage with no branch point, from sublining to homeostatic to lining fibroblast to adipocyte (median pseudotime 31.2, 38.7, 53.3 and 79.4). Monocle3 gave an identical population order (Spearman ρ = 1.00) (Fig. 7a–d, Supplementary Fig. 30a–c). Synovium refitted alone returned the same order. In the IPFP, the trajectory extended from the matrix-high fibroblast through the adhesion, SMOC1+/GHR+ and CXCL14+/IGFBP7+ subtypes to the adipocytes, with only the lining-like fibroblast placed beyond them. Rooted by the same rule, the IPFP trajectory began at the matrix-high fibroblast, which carries four of the panel’s five markers (DPP4, COMP, MFAP5 and PI16) (Fig. 7g, Supplementary Fig. 31). Along the axis 684 genes rose and 237 fell in all three trajectories, the adipocyte gene AQP7 rising and matrix and fibroblast genes (COMP, CILP, CTHRC1) falling in all three (Fig. 7h), and lipid-handling genes (PLIN5, GPAM, PCK1) rising in the IPFP’s trajectory.

**Fig. 7.**
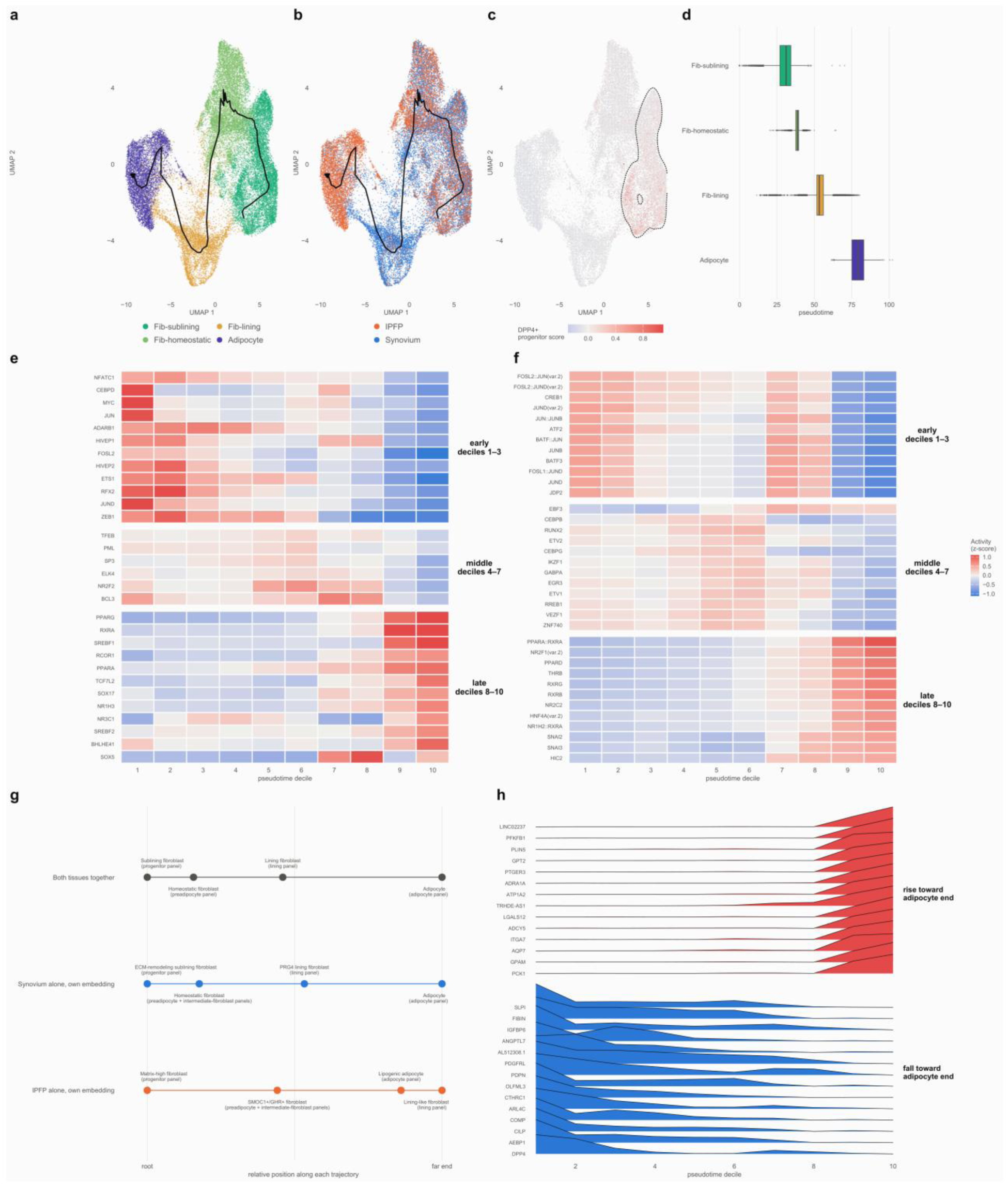
One continuous stromal trajectory across healthy synovium and IPFP, and the regulators that change along it. **a** The mesenchymal manifold of the joint reference (30,455 nuclei, six donors, four populations) with the fitted trajectory. **b** The same manifold by dissected tissue of origin. **c** The published DPP4+ progenitor panel scored per nucleus; the dashed contour marks the population scoring highest, the root of the fit. **d** Pseudotime by population. **e** Regulon activity (AUCell) of the leading transcription factors of each stage, donor-averaged and standardized within tissue; rows are factors meeting the donor-support rules (Methods), grouped by the third of the axis in which they peak, and columns are pseudotime deciles. **f** The same for motif accessibility (chromVAR, 633 JASPAR motifs); features reaching 90% of their peak in another third of the axis, including 21 of the 34 AP-1 motifs, are excluded from the mid-axis rows. e and f share one color scale (z-score within tissue), symmetric about zero and clipped at ±1.10 (the 99th percentile of |z|; 6 of 660 tiles lie beyond it and are shown at the scale end). **g** The trajectory refitted within each tissue’s own reference; the x axis is relative position along each fit, from its root to its far end. Each node is the population nominated by a published marker panel (panel in brackets). In each tissue fitted alone the preadipocyte and intermediate-fibroblast panels nominate the same population (homeostatic fibroblast in synovium, SMOC1+/GHR+ fibroblast in the IPFP); in the joint reference the intermediate-fibroblast panel’s best match is not a mesenchymal population, so it has no node there. **h** Genes with the largest change along the trajectory, rising or falling toward the adipocyte end (genes moving the same way in at least five of six donors), donor-averaged per decile and standardized per gene, one profile per gene: the rise is a step at deciles 8–9, and the fall is concentrated between deciles 1 and 2 and continues slowly thereafter.

Distinct regulatory programs were active at the root, middle and end of the trajectory (Fig. 7e,f, Supplementary Fig. 32a,b). AP-1 target-gene activity was highest at the root: from the first two deciles to the last two, the JUND, JUN and FOSL2 regulons fell by 1.7, 1.2 and 1.3 standard deviations with all six donors agreeing. The MYC and CEBPD regulons, which mark the adipogenic route in mouse single-cell trajectory analysis,^25^ also fell along the trajectory, as did ZEB1 and ETS1. AP-1 motif accessibility had two maxima of similar height, at the sublining and at the lining fibroblast, for all 34 AP-1 motifs (second maximum 84–100% of the first). In the middle deciles the CEBP motifs (CEBPB, CEBPG) and the NR2F2 regulon peaked. At the terminal end the two modalities agreed: PPARG, RXRA, SREBF1, PPARA, TCF7L2 and NR1H3 rose by 0.7 to 1.9 standard deviations in six of six donors, and the nuclear-receptor motifs opened, led by PPARA::RXRA (Fig. 7e,f, Supplementary Fig. 32b).

Compared population by population along the trajectory, the four populations differ in regulatory state (Fig. 8a,b). The sublining root has the highest AP-1 target-gene activity and a high matrix output, which in the IPFP is the highest of any population (Figs. 2e and 4e). AP-1 motifs are as accessible in the lining as in the sublining, although the lining expresses fewer AP-1 target genes, and the two boundaries most strongly resolved by chromatin, the synovial lining and the adipocyte, both lie in the second half of the axis.

**Fig. 8.**
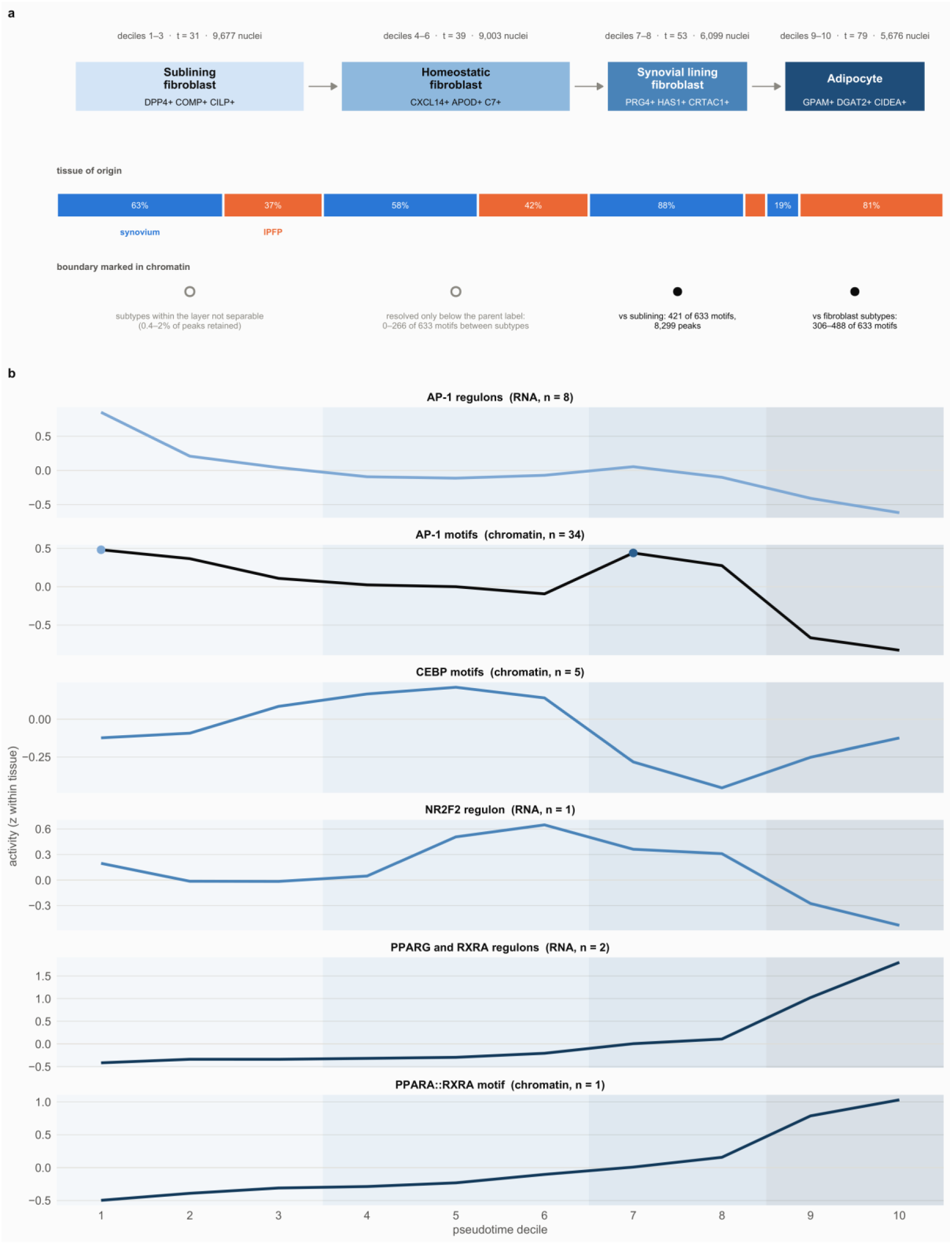
The stromal trajectory summarized population by population. **a** The four populations in order along the trajectory, positioned by pseudotime decile so that they align with panel b. Decile span, median pseudotime and number of nuclei are given above each; arrows indicate direction from the chosen root, which these data do not determine. Middle row, tissue of origin of all nuclei in the deciles each population spans (not of the population’s own nuclei; for those see Fig. 6c). Bottom row, whether the boundary between each population and its neighbor is marked in chromatin, determined independently of the trajectory: for the lining and adipocyte, from the compartment refits (Figs. 3f and 5f); for the sublining, from the fraction of enriched peaks retained within the fibroblast compartment (Supplementary Fig. 12b); for the homeostatic fibroblast, from the IPFP subdivision test (2 of 633 motifs between the original labels, up to 266 between subtypes; Supplementary Fig. 23). Filled markers, boundaries marked in chromatin (lining, adipocyte); open markers, not separable at the population label (sublining; homeostatic fibroblast, resolved only between subtypes). **b** Six regulatory tracks on the same axis, donor-averaged and standardized within tissue, with the population bands repeated behind each; family tracks are the mean over every matching feature of the 633 motifs or 141 regulons, and n is that count. Tracks 1 and 2 show the main discrepancy: AP-1 target-gene activity falls from the root, whereas AP-1 motifs remain accessible at both the root and the lining fibroblast (two maxima marked). CEBPA is excluded from the CEBP track, being the family member with the strongest terminal peak.

### Healthy stromal cells rarely cycle, no dividing adipocytes were detected, and organelle-turnover programs decline along the trajectory

To assess cell turnover in adult tissue, we identified dividing nuclei (Methods). Immune cells were dividing in both tissues (IPFP 51 of 2,460 nuclei, 2.07%; synovium 39 of 5,984, 0.65%; four and five of six donors): T cells at similar rates in the two tissues (12 of 426, 2.8%, and 14 of 620, 2.3%), and macrophages more often in the IPFP than in synovium (34 of 1,793, 1.9%, against 16 of 4,560, 0.35%). Very few stromal cells were dividing: 7 of 16,307 synovial fibroblast nuclei (0.043%, three of six donors), 1 of 8,067 IPFP fibroblast nuclei (0.012%), and none of 5,165 adipocyte nuclei in any adipocyte subtype of either tissue (Fig. 9a,b). The dividing fibroblasts were spread across subtypes: three in the PRG4 lining layer, two in fibrocartilage, one each in the homeostatic and DPP4+ sublining populations (1 of 2,721 DPP4+ nuclei), and one in the IPFP’s lipid-laden subtype. On the trajectory they fell between deciles 2 and 8, and none fell in the adipocyte deciles 9 and 10 (Fig. 9c). At the observed division rate, synovial fibroblasts would take about three years (range one to eleven) to replace their own number, and fibroblast division exceeded the rate required for 10% annual adipocyte renewal^26^ by at least 25-fold in synovium and met it at the point estimate in the IPFP (Fig. 9d).

**Fig. 9.**
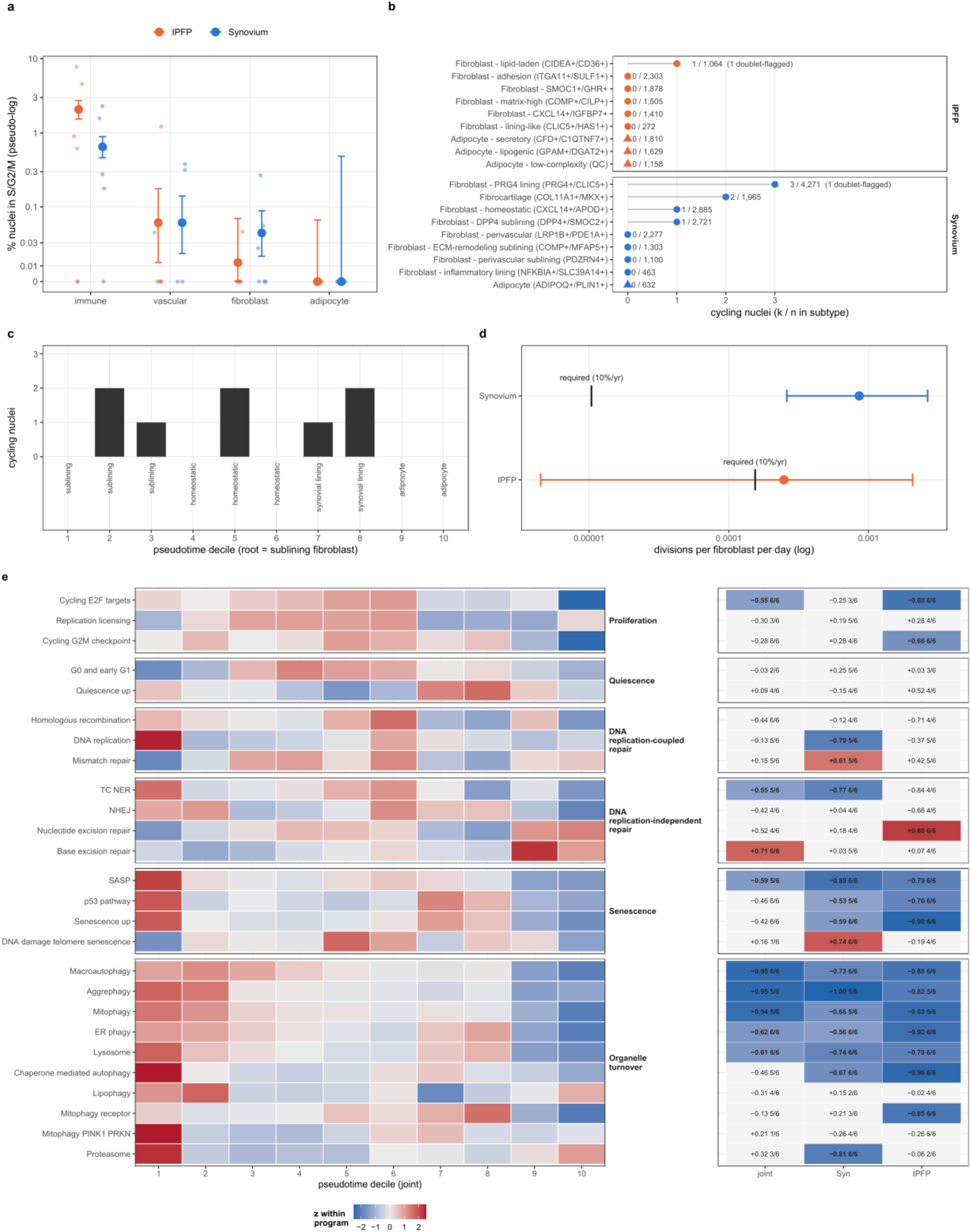
Stromal cells of the healthy joint rarely cycle, no dividing adipocytes were detected, and organelle-turnover programs decline along the stromal trajectory. **a** Stromal cells are almost never cycling. Percentage of nuclei in S or G2/M per compartment and tissue, restricted to nuclei whose joint label and label in the tissue’s own reference agree. Large points, pooled rate; bars, exact Poisson 95% interval (zero counts at 0 with their one-sided upper bound); small points, individual donors. Pseudo-log scale. **b** No cycling nuclei in any adipocyte subtype, and fibroblast division is rare and dispersed. Label-concordant cycling nuclei in each native mesenchymal subtype of the IPFP (top) and synovium (bottom); labels, cycling nuclei over nuclei in the subtype; circles, fibroblast subtypes; triangles, adipocyte subtypes; doublet-flagged nuclei are noted. **c** Division lies in the fibroblast stretch of the trajectory, with no cycling nuclei in the adipocyte deciles. Cycling nuclei per pseudotime decile of the joint trajectory (root, sublining fibroblast; 30,455 mesenchymal nuclei), label-verified mesenchymal nuclei only; the modal population of each decile is given below the axis. **d Observed fibroblast division could supply the estimated rate of adipocyte renewal. Progenitor divisions per fibroblast per day required by 10% annual adipocyte renewal (tick labeled required)**^26^ **against the rate supplied by the observed fibroblast division (point; interval combines the Poisson 95% interval with an assumed 8 to 16 h S/G2/M window). e** Organelle-turnover programs and the senescence and SASP sets peak at the fibroblast root. Twenty-six turnover programs along the joint trajectory: depth-adjusted module scores, donor-averaged per decile and z-scored within program (left), with Spearman ρ against decile and the number of concordant over tested donors in the joint trajectory and in each tissue fitted alone (right; shaded where |ρ| ≥ 0.5 with at least five of six donors). n = 54,716 nuclei (a) and the 30,455 mesenchymal nuclei of the joint trajectory (c, e); six donors, two tissues.

Organelle-turnover programs fell from the fibroblast root to the adipocyte in the joint trajectory and in each tissue fitted alone: macroautophagy (Spearman ρ −0.95, −0.73 and −0.88), lysosome (−0.61, −0.74, −0.78) and ER-phagy (−0.62, −0.56, −0.92), each in six of six donors, and aggrephagy (−0.95, −1.00, −0.82) in five of six (Fig. 9e). Senescence and SASP gene sets were also highest at the root (Fridman senescence set^27^ −0.42, −0.59, −0.98; six of six donors), and E2F-target activity fell along the trajectory (joint ρ −0.55, IPFP −0.88). Stromal cells of the healthy adult joint therefore rarely cycle: no dividing adipocytes were detected, the rare dividing fibroblasts are dispersed across subtypes and along the trajectory, and the expression of organelle-turnover programs declines from the root to the adipocyte.

## Discussion

Synovium and IPFP are routinely sampled in OA research, but neither had been comprehensively and quantitatively described in health. This study provides that baseline for both tissues from the same donors, at tissue level and, in the same nuclei, by RNA and chromatin. The data indicate that healthy cells in these tissues express programs for producing and protecting the extracellular matrix and lubricant of the joint and for lipid storage, are predicted to signal to the vasculature, and rarely divide. Within the stroma, chromatin accessibility most strongly resolves two boundaries, between lining and sublining fibroblasts and between fibroblasts and adipocytes, whereas the remaining subtypes differ quantitatively (Fig. 10).

**Fig. 10.**
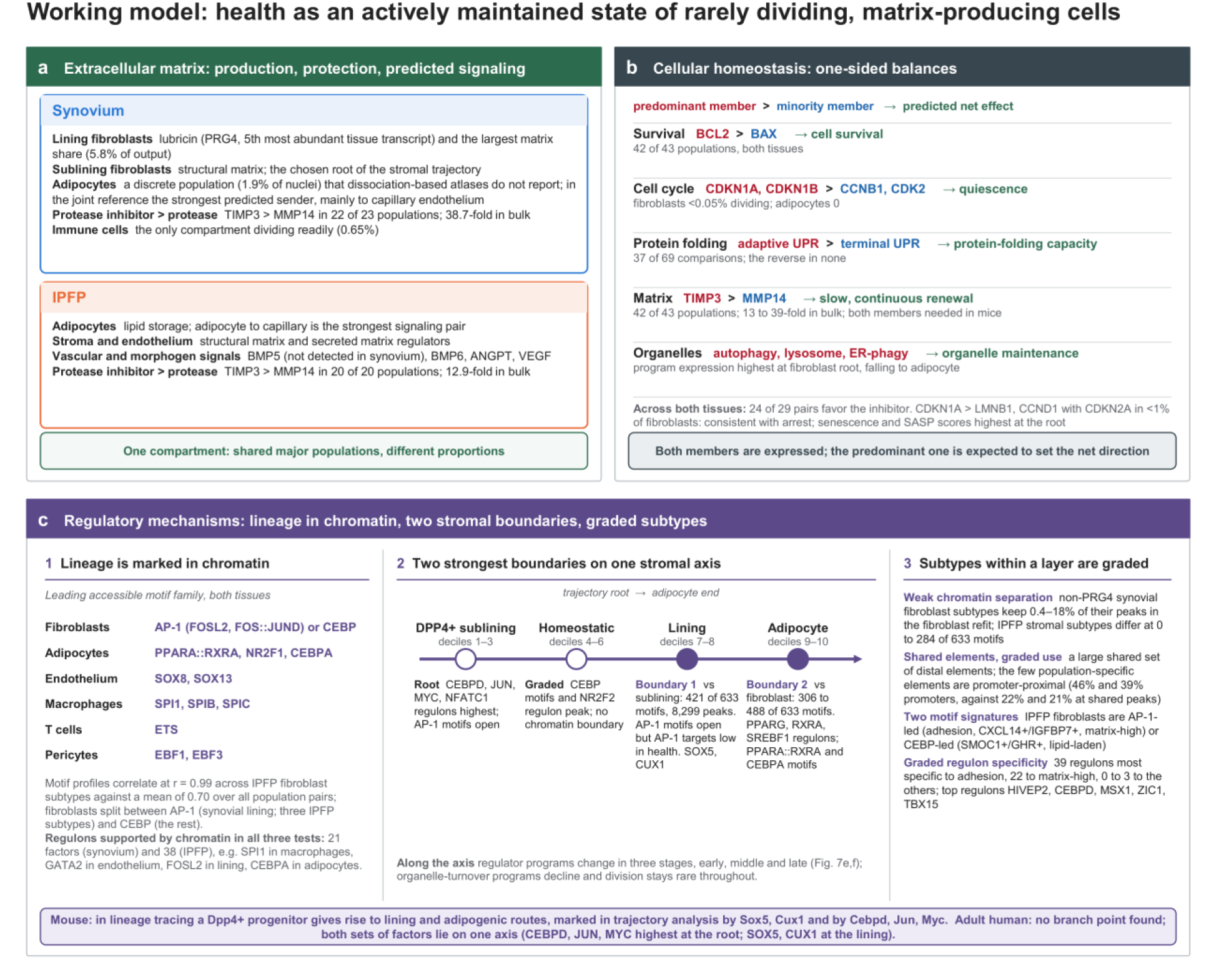
What healthy synovium and IPFP cells do, and how they are regulated: a summary. A schematic synthesis of Figs. 1–9 and Supplementary Fig. 7; every value is taken from the Results or these figures. **a** Extracellular matrix. In synovium the lining fibroblasts produce lubricin (PRG4, the fifth most abundant tissue transcript) and carry the largest structural matrix share (5.8% of transcript output), the sublining fibroblasts produce structural matrix, the adipocytes form a discrete population (1.9% of nuclei), not reported by dissociation-based atlases, that in the joint reference is the strongest predicted sender, mainly to capillary endothelium, and immune cells are the only compartment dividing readily. In the IPFP the adipocytes store lipid and form the strongest single signaling pair, adipocyte to capillary endothelium, stroma and endothelium express structural matrix and secreted matrix regulator transcripts, and the vascular and morphogen signals BMP5 (not detected in synovium), BMP6, ANGPT and VEGF are IPFP biased. In both tissues TIMP3 exceeds MMP14 in nearly every population (38.7-fold in bulk synovium, 12.9-fold in bulk IPFP), and at joint-reference resolution the two tissues form one stromal compartment of the same major populations in different proportions. **b** Cellular homeostasis. For each program the predominant (red) and minority (blue) member of the opposing gene pairs and the net effect predicted from the imbalance (green); the organelle row is a trend along the trajectory and has no opposing member; 24 of the 29 pairs favor the inhibitor in each tissue (Supplementary Data 3). Both members are expressed, so the pairs do not neutralize each other; because they are strongly one-sided, the predominant member is expected to set the net direction of each process. Fold differences are given from bulk RNA-seq, because nuclear folds are inflated for long genes. Organelle-turnover programs are highest at the fibroblast root of the trajectory and fall toward the adipocyte. **c** Regulatory mechanisms, at three levels. (1) Lineage: the leading accessible motif family largely follows lineage, and fibroblasts split between AP-1 (the synovial lining and three IPFP subtypes) and CEBP, and the regulons supported by chromatin in all three cross-modal tests recover an expected regulator of each testable lineage. (2) The stromal axis, rooted at the population scoring highest for the DPP4+ progenitor signature (the DPP4+ sublining), to the adipocyte, with the two boundaries most strongly resolved by chromatin (filled nodes): the synovial lining, where AP-1 motifs are accessible but AP-1 target genes are expressed at a low level, and the adipocyte, marked by PPARG, CEBPA and the PPARA::RXRA motif; regulator programs turn over in early, middle and late stages along it. (3) Subtypes within a layer share one regulatory program and are distinguished quantitatively: they share most accessible elements, differ by few population-specific, mostly promoter-proximal ones, split in the IPFP into AP-1-led and CEBP-led motif signatures, and differ in how many regulons are most specific to them. Bottom band: in mouse lineage tracing a Dpp4+ progenitor gives rise to two routes; single-cell trajectory analysis marks the lining route by Sox5 and Cux1 and the adipogenic route by Cebpd, Jun and Myc.^25^ In the adult tissue profiled here no branch point was found: the human orthologs of both sets of factors are present, on one continuous axis (CEBPD, JUN and MYC highest at the root, SOX5 and CUX1 specific to the lining fibroblast). Ratios are of mRNA and indicate direction only; they do not measure protein stoichiometry.

ECM transcript output is a main feature that distinguishes stromal populations. The populations of both tissues were far more alike in their core cellular machinery than in what they produced: transcription and RNA processing accounted for close to 15% of counts in every population, whereas the structural ECM share varied about seven-fold between populations, to the same extent in both tissues (6.7-fold in synovium, 6.9-fold in the IPFP). ECM output also distinguished populations and dominated their predicted interactions. At tissue level PRG4 was the fifth most abundant transcript of synovium, indicating that lubricin is a major product of the lining. Lubricin protects the cartilage surface and restrains synovial overgrowth.^28^ In the IPFP these functions were divided between lineages: the stroma expressed most structural matrix transcripts, the endothelium expressed secreted regulators of it, and the adipocytes expressed a lipid-handling program with little matrix output. Because populations differ mainly in the amount of ECM transcripts they produce, a disease-associated change in a population’s matrix output may reflect altered output by an existing population as readily as the appearance of a new cell type. Expression data alone cannot distinguish these possibilities.

Healthy cells expressed both the activating and the inhibitory member of each opposing gene pair. The two were not balanced, however. Most pairs were strongly one-sided, and the predominant member is expected to set the net direction of each process. Across both tissues 24 of the 29 pairs favored the inhibitor, a direction that was reproduced in bulk tissue. For example, TIMP3 exceeded MMP14 in 42 of 43 populations and by 13-to 39-fold in bulk, BCL2 exceeded BAX in 42 of 43 populations, cyclin-dependent kinase inhibitors exceeded their targets in 20 of 23 synovial populations, and the adaptive arm of the unfolded protein response exceeded the terminal arm in about half of the comparisons and was never exceeded by it. The predicted net result is matrix preservation, cell survival, quiescence and protein-folding capacity. This profile is consistent with the needs of quiescent cells that secrete large amounts of collagen and proteoglycan, and the adaptive unfolded protein response is the homeostatic form of that response.^29^ TIMP3 binds sulfated glycosaminoglycans in the ECM,^30^ so its excess may reflect TIMP3 deposited in the matrix produced by these cells. The less abundant member is nonetheless required: mice lacking MMP14 develop arthritis from inadequate collagen turnover,^31^ and mice lacking TIMP3 lose joint collagen and aggrecan with age.^32^ These data suggest that healthy tissue expresses both arms, with protease-inhibitor transcripts in excess of protease transcripts, consistent with quiescence being an actively maintained state.^33^ Testing this will require biochemical measurement of protein levels and protease activity. CDKN1A also exceeded LMNB1 and CCND1, but CDKN2A was detected in fewer than 1% of fibroblast nuclei, so we interpret this as the cell-cycle arrest of quiescent fibroblasts rather than an accumulation of senescent cells, although senescence and SASP gene-set scores were highest at the fibroblast root and a minor senescent-like component cannot be excluded. These mRNA ratios indicate the direction of a process and do not measure protein stoichiometry. Their size in nuclei is also inflated for long genes such as BCL2, which is why we base the argument on direction.

Healthy stromal cells also rarely divide. No adipocyte nuclei and fewer than 0.05% of fibroblast nuclei were dividing, and at this rate synovial fibroblasts would replace their own number in about three years (range one to eleven). In contrast, 0.65% of synovial and 2.07% of IPFP immune nuclei were dividing, including T cells at 2–3% in both tissues and macrophages at 0.35% in synovium and 1.9% in the IPFP. This is consistent with locally renewing resident macrophages^12^ and shows that division was detectable. Along the stromal trajectory, the expression of organelle-turnover programs changed, with autophagy, lysosome and ER-phagy programs highest at the fibroblast root and falling toward the adipocyte. In this snapshot, tissue maintenance therefore appears to rely more on existing cells than on proliferation. Lineage-tracing studies describe sublining progenitors that proliferate after injury and give rise to lining fibroblasts,^34,35^ and a Dpp4+ sublining progenitor that also gives rise to IPFP adipocytes.^25,36^ In the healthy adult joint, dividing DPP4+ nuclei were rare (1 of 2,721), although the overall fibroblast division rate would be sufficient to supply the estimated rate of adipocyte renewal (Fig. 9d).

The regulatory data are consistent with populations that differ quantitatively, separated by few categorical boundaries. Chromatin accessibility clearly marked lineage, and subtype much less, in both tissues. Within the stroma, chromatin most strongly resolved two boundaries. The first separated synovial lining from sublining fibroblasts and was marked by AP-1 motifs. The second separated adipocytes from fibroblasts and was marked by PPARG, CEBPA and PPARA::RXRA, the canonical adipogenic machinery.^37^ The other fibroblast subtypes had highly correlated regulon programs (r ≥ 0.87 in the IPFP) and differed in graded fashion, although IPFP stromal subtypes still differed at up to 284 motifs, the largest of these differences involving the doublet-enriched lipid-laden subtype. They made graded use of a large shared set of distal elements and had few population-specific elements, most of them promoter-proximal. In the IPFP, they also split into AP-1-led and CEBP-led motif signatures, and regulon specificity was graded and concentrated in two subtypes (Fig. 10c). Because these differences are continuous, increasing cluster resolution is unlikely to define a fixed number of fibroblast subtypes. This may also explain the range of progenitor definitions proposed for adipose stroma.^21,38,39^ The lining boundary has one further feature. Lining fibroblasts keep AP-1 motifs accessible while expressing fewer AP-1 target genes than the sublining (Supplementary Fig. 7e). This pattern suggests that AP-1 elements in the lining are open but little used in health, which could allow a rapid response to stimuli.

At the resolution of the joint reference, the two tissues formed one stromal compartment in different proportions. In both, predicted signaling ran from matrix-producing and adipocyte populations to the endothelium, and the IPFP showed more inferred signaling, with BMP showing the largest pathway shift, about half of it carried by BMP5, which was not detected in synovium.

The stromal populations of both tissues lay on one continuous transcriptional trajectory ending at the adipocyte. Lineage tracing in mouse places the divergence of synovium and IPFP at a Dpp4+ progenitor, and single-cell trajectory analysis places Sox5 and Cux1 on the lining route and Cebpd, Jun and Myc on the adipogenic route.^25^ Both sets of factors are present in adult tissue on one unbranched axis, the lining-route factors at the lining fibroblast and the adipogenic-route factors at the sublining root. A bifurcation that occurs once during development need not persist as a branch point in adult tissue, which may explain why we did not recover the branch point proposed by Tang et al.^18^ (Supplementary Fig. 37).

This study has limitations. Six donors cannot describe variation with age, sex or body mass index, and the donors were young, so these references describe healthy adult tissue and do not extend to aged tissue without disease. Nuclear RNA is not whole-cell RNA and underrepresents cytoplasmic transcripts.^40,41^ Pseudotime describes similarity between cells and does not show movement of a cell from one state to another, and the cycling fractions are a single time point. Regulons are inferred from co-expression, motif accessibility identifies a factor family and cannot resolve individual factors, and cell–cell communication is predicted from expressed ligand–receptor pairs. More broadly, several of these analyses generate hypotheses about the healthy joint, among them the predominance of inhibitors in opposing gene pairs, signaling directed toward the endothelium and a continuous stromal trajectory; these will need to be confirmed with protein-level, spatial and functional methods.

In conclusion, this study provides the first single-nucleus multiomic reference of healthy human synovium and IPFP, with a tissue-level transcriptome of both tissues in a larger cohort. The data suggest that rarely dividing stromal cells are transcriptionally active in health: they express matrix and lubricant genes, organelle-turnover programs and transcripts that restrain protease activity, cell death and division, and are predicted to signal toward the vasculature. These cells lie on one continuous transcriptional axis with two boundaries most strongly resolved by chromatin. These references provide the baseline against which osteoarthritis, rheumatoid arthritis, injury and aging can be interpreted, and can reveal disease-associated changes that would be missed without a healthy reference.

## Methods

### Ethics approval

Human tissue collection was approved by the Scripps Human Subjects Committee (20-7636). Informed consent/authorization was obtained for research use of donor tissue.

### Donors and tissue collection

Knee tissues were obtained from organ donors without a history of knee osteoarthritis, rheumatoid arthritis or knee injury, through the tissue procurement program described under Ethics approval. Synovium was taken from the medial gutter and the IPFP from the anterior compartment, from the same knee except where stated below. Outerbridge macroscopic knee grade^42^ was recorded for every donor and only knees graded 0 or 1 were classified as healthy. Two donor sets are used in this study. The bulk RNA sequencing cohort comprised 14 synovium donors (11 male, 3 female; mean age 31.9 ± 6.7 years, range 18–40; mean BMI 27.8) and 18 IPFP donors (13 male, 5 female; mean age 31.3 ± 7.9 years, range 18–48; mean BMI 28.3), with 13 donors contributing both tissues. The single-nucleus multiome cohort comprised six donors (three male, three female; mean age 33.3 ± 11.1 years), each contributing both synovium and IPFP, giving twelve libraries. For five donors both tissues came from the same knee; for one (D3) the single-nucleus synovium came from the contralateral knee, also macroscopic grade 1, while the IPFP and both bulk libraries came from the same knee as each other. Three of these donors are also in the bulk synovium cohort and five in the bulk IPFP cohort. Recorded sex was confirmed from sex-chromosome expression in each multiome donor: in the donor pseudobulk, XIST and the summed Y-linked genes (RPS4Y1, DDX3Y, UTY, USP9Y, EIF1AY, KDM5D, ZFY, NLGN4Y and TMSB4Y) differed by more than 100-fold in the direction of the recorded sex, independently in synovium and IPFP, for all six donors (Supplementary Fig. 2a). Data and experimental protocols associated with this study will be available through the SPARC Portal (RRID:SCR_017041) upon publication at the following DOIs: https://doi.org/10.26275/uhbq-r2s7, https://doi.org/10.26275/ifye-wvaq, https://doi.org/10.26275/t9lp-ovyr, and https://doi.org/10.26275/2sia-jzkd.43-46 This study used a subset of donors from these datasets; the SPARC subject identifier, age, sex and assays of each donor are listed in Supplementary Data 5.

### Nucleus isolation and multiome library preparation

Nuclei were isolated from frozen IPFP and synovium tissues, stored in liquid nitrogen, using the Singulator S200 Automated Tissue Dissociation System (Precision Cell Systems; RRID:SCR_029160). The procedure was performed according to a previously published protocol.^47^ Prior to loading, 100–150 mg of frozen tissue was finely shaved and minced into small pieces using a scalpel on a pre-chilled petri dish. The tissue fragments were then transferred to a pre-cooled Singulator cartridge containing 75 µL of RNase inhibitor (40 U/µL). The cartridge was placed into the Singulator, and the low-volume nuclei isolation program was run. Following homogenization, the resulting suspension was passed through 20 µm and 40 µm strainers to remove tissue and cellular debris. The flow-through was collected into a 5 mL LoBind tube and centrifuged at 150 × g for 10 minutes at 4 °C. The supernatant was carefully removed, and the nuclei pellet was resuspended in 600 µL of PBS containing 0.02 mg/mL recombinant albumin and 1 U/µL RNase inhibitor. To further eliminate cellular debris, a second centrifugation was performed at 150 × g for 10 minutes at 4 °C. The supernatant was discarded, and the nuclei pellet was resuspended in 100 µL of diluted nuclei buffer. For nuclei counting and quality assessment, 18 µL of the nuclei suspension was mixed with 2 µL of Acridine Orange. Nuclei were counted using the Countess II FL Automated Cell Counter (Thermo Fisher Scientific, RRID:SCR_025370; AOPI staining). Additionally, nuclei morphology and membrane integrity were visually inspected under a microscope at 40× or 60× magnification (AOPI staining), following the nuclear integrity check guidelines provided by 10x Genomics. Based on the estimated nuclei concentration, the nuclei pellet was resuspended in chilled diluted nuclei buffer to a target concentration of 4,000–5,000 nuclei/µL for a final recovery of 20,000 nuclei in 5 µL. Libraries were prepared with the Chromium Next GEM Single Cell Multiome ATAC + Gene Expression kit (10x Genomics, PN-1000283), with 33,000 nuclei loaded per reaction, and sequenced paired-end on an AVITI system (Element Biosciences; RRID:SCR_029159).

### Bulk RNA sequencing

Total RNA was extracted from whole tissue as described^48^ and sequenced as part of the same donor program. Reads were adapter-and quality-trimmed with Trim Galore (Cutadapt^49^), aligned to the GRCh38 human reference with HISAT2^50^ and assigned to GENCODE v44 (RRID:SCR_014966) genes with featureCounts,^51^ and gene-level counts were converted to transcripts per million (TPM). Library-level quality control was applied before any analysis, and only the QC-passed, condition-normal libraries enter the results reported here. A gene was called detected when it reached at least 1 TPM in at least half of the donors of a tissue. We did not call a gene detected from any single donor, because the number of genes detected per library reflected non-coding background in libraries with low exonic assignment (genes detected tracked the non-coding share at r = 0.98 in synovium and 0.93 in the IPFP), whereas protein-coding detection varied far less (5,173–14,221 genes in synovium and 4,209–13,579 in the IPFP). The most abundant transcripts were ranked by the median TPM across donors, which limits the influence of individual libraries with atypical composition; mitochondrial and ribosomal transcripts were flagged but retained, because they are among the most abundant transcripts in bulk tissue. For the tissue contrast, genes reaching 5 TPM in at least one tissue were compared as log2((IPFP + 1)/(synovium + 1)) of the per-tissue medians. Because TPM is length-normalized, ratios between two genes within a sample are molar and are therefore directly comparable with the UMI-based ratios computed in the nuclei.

### Bulk and single-nucleus gene detection and abundant-gene function

Bulk and single-nucleus genes were matched on Ensembl (RRID:SCR_002344) gene identifier through the Cell Ranger reference, and only the 36,366 genes annotated in both were compared. Single-nucleus expression was taken from the tissue pseudobulk per donor, as counts per million; because UMI counts are molar, this is the nuclear analog of TPM. A gene was called detected on a platform at 1 TPM (bulk) or 1 CPM (nuclei) in at least half of that platform’s donors, with 5 and 10 as sensitivity floors. Single-nucleus counts cover the gene body and bulk counts the exons, so intron-rich genes are more easily detected in nuclei. For function, the 200 most abundant protein-coding transcripts per tissue by median TPM, excluding mitochondrial-encoded and ribosomal protein genes, and the part of each not in the other tissue’s top 200, were tested for over-representation in MSigDB (RRID:SCR_016863) Hallmark and GO (Gene Ontology, RRID:SCR_002811) Biological Process sets of 10 to 500 genes (msigdbr) by one-sided hypergeometric test against the protein-coding genes detected in that tissue, with Benjamini–Hochberg correction; the top 500 were tested as a sensitivity analysis.

### Read processing, quality control and filtering of the multiome data

Sequencing reads were processed with Cell Ranger ARC against the GRCh38 reference, which returns a joint gene expression and ATAC fragment output per library. Nuclei were retained on the gene expression assay with 1,000–25,000 UMIs and less than 5% mitochondrial reads, and on the ATAC assay with a nucleosome signal below 2, a transcription start site enrichment above 1 and at least 500 (synovium; at most 100,000) or 250 (IPFP) fragments in peaks. Peaks were restricted to the standard chromosomes. Doublets were called with scDblFinder^52^ per library on the twelve nuclei libraries of the joint reference, once with population labels as clusters and once from random cell pairs, and were not removed from any reference. The calls were used in two places: in the IPFP subclustering, where doublets were held out of clustering and assigned afterwards by nearest-neighbor vote so that a doublet could not generate a subcluster of its own, and in a sensitivity analysis of the joint-reference cell–cell communication results.

### Normalization, integration, clustering and annotation

For each tissue, single-nucleus transcriptomes were normalized with SCTransform^53,54^ (glmGamPoi^55^) regressing percent mitochondrial reads, integrated across all libraries of the larger donor program processed with the same pipeline (synovium, 16 libraries; IPFP, 17 libraries; including aged and osteoarthritic donors) by canonical correlation analysis^56^ on the SCT assay with 3,000 integration features and 50 anchor dimensions, clustered on RNA alone at the resolution maximizing a combined silhouette and modularity score (synovium 0.6, 23 clusters; IPFP 0.2, 15 clusters) and annotated from marker genes. The six healthy donors of each tissue were then re-integrated alone (SCTransform per donor without covariate regression, 3,000 integration features, 30 anchor dimensions and 30 principal components), with the population labels carried over by nuclear barcode; all analyses reported here use these healthy-only objects. For the multiome objects, gene expression was normalized with SCTransform regressing percent mitochondrial reads and donors were integrated by canonical correlation analysis on the SCT assay with 3,000 integration features and 50 anchor dimensions. ATAC data were normalized by term frequency–inverse document frequency, reduced by singular value decomposition, and LSI components 2–40 were used, the first being dropped because it correlates with sequencing depth. A weighted nearest-neighbor graph combining both modalities^57^ was used for the joint embedding, and multiome nuclei were labeled by exact barcode match to the same nuclei in the tissue’s RNA reference (100% of retained nuclei in both tissues), so no label was inferred; the marker genes naming each population are given in the population names themselves. Nuclei were matched between modalities by nuclear barcode, so the chromatin analyses use the transcriptomic labels of the same nuclei.

### Statistical framework

Nuclei from the same donor share genotype, chromatin state and sample handling, and are therefore not independent observations. Every inferential test in this study treats the donor as the biological replicate, and no test is performed across nuclei. Expression, program shares, matrisome shares and opposing-gene ratios were computed as pseudobulk, one value per donor and population, and are reported as the mean across donors with the standard deviation, together with the number of donors supporting the direction of each claim. Differential accessibility of peaks and motif activity were tested by donor-paired pseudobulk in models blocking on donor, fitted with DESeq2^58^ for peak counts and limma^59^ for motif deviations, both as ∼ donor + population, with population contrasts specified as one population against the mean of the others and significance at a Benjamini–Hochberg adjusted p below 0.05. chromVAR deviation scores are approximately normal and already corrected for GC content and mean accessibility, so a linear (Gaussian) model was used. Populations entered a test only if they contributed at least 100 nuclei in total and at least 25 nuclei in at least four of the six donors; populations failing that rule are named in the Results and excluded from the analyses concerned. Where a claim rests on a trend without a formal test, the number of donors agreeing in direction is stated, and leave-one-donor-out refits are reported for the trajectory.

### Transcriptional program partition

The transcriptome was partitioned into twelve programs defined from published Gene Ontology^60^ and MSigDB^61^ gene sets, each gene assigned to exactly one program by a fixed priority order so that program shares sum without double counting; the assignment, the conflicts it resolves and a sensitivity analysis over the priority order are reported in the supplement. A program’s value in a population is the share of that population’s pooled sequencing counts falling in its genes; per-donor shares give the standard deviations shown. The same partition was applied unchanged to the IPFP and to the bulk TPM matrices. To test whether the twelve programs missed a signal, all remaining GO terms were scanned for enrichment in any population outside the partition.

### Opposing gene pairs

Twenty-nine pairs of genes were assembled across eight programs, each pair comprising one gene product that drives a process and one that opposes it, selected from established biology and each supported by a primary or review reference (Supplementary Data 3); 19 are direct antagonists acting on the same molecular step and 10 are indirect indicators of the balance between two arms of a pathway. For each pair, each population and each donor, the ratio of the two genes’ pseudobulk expression was computed, and a pair was called as favoring one side in a population only when it did so in every donor by at least two-fold. Ratios are reported as the median across populations. Pairs in which either gene fell below the detection floor in a population were recorded as undetected rather than imputed. Robustness was assessed by a pair-level binomial sign test (because population-by-pair calls are not independent), against 5,000 random pairs of expressed genes, by leave-one-donor-out resampling, with both genes required at 1 CPM or more, and in the bulk TPM of the same tissues. Because single-nucleus counts cover the gene body, the nucleus-to-bulk difference in each ratio was compared with the length ratio of the two genes. Ratios are of transcripts; they indicate the direction of a balance and do not measure protein stoichiometry or activity.

### Regulon inference

Gene regulatory networks were inferred with pySCENIC,^62,63^ which builds co-expression modules around each transcription factor, prunes each module to the targets carrying that factor’s binding motif within their regulatory region, and scores the resulting regulon in every nucleus by AUCell. Motif pruning used the hg19 RefSeq cisTarget (RRID:SCR_024808) databases (mc9nr; 500 bp upstream and 10 kb centered on the transcription start site, seven species) with the matching motif annotation (motifs-v9-nr); cisTarget matches databases to expression data on gene symbol. Regulons were assigned to populations by regulon specificity score, and population similarity was computed as the correlation of regulon activity profiles. Because a regulon’s specificity depends on the set of populations it is scored against, the fibroblast compartment was additionally scored within itself, and both the within-compartment and the global specificity scores are reported.

### Chromatin accessibility: peaks, differential accessibility and peak-to-gene links

Peaks called by Cell Ranger ARC in each library were merged into one peak set per tissue (20 bp–10 kb, standard chromosomes; 124,364 peaks in synovium and 155,673 in the IPFP), on which every library was re-quantified. Peak enrichment was tested per population by donor-paired pseudobulk as described above, restricted to peaks with at least 10 counts in at least four donor–population pseudobulk samples. A peak was called exclusive when it was enriched in exactly one population; because that quantity depends on cluster resolution, it is reported at two resolutions in the IPFP. Promoter overlap (within 2 kb of an annotated transcription start site) was computed with ChIPseeker^64^ and compared within quartiles of peak coverage, so that the promoter share of exclusive peaks is not an artifact of their depth. Peaks were linked to genes within 500 kb by correlation between accessibility and expression across nuclei, computed at tissue level.

### Motif accessibility

The 633 human JASPAR (RRID:SCR_003030) CORE motifs^65^ were matched to peaks and per-nucleus deviation scores computed with chromVAR,^66^ which corrects for GC content and mean accessibility. Scores were averaged to one profile per donor and population and tested in the donor-blocked linear model described above. A motif is described as ranked first in a population when it carried the largest one-versus-rest activity difference among motifs reaching significance. Because motif accessibility reflects a binding preference shared within a transcription factor family, these assignments identify a family and are reported at that level.

### Cross-modal validation of regulons against chromatin

For every transcription factor carrying both a regulon and a motif, three tests were applied in the population where its regulon was most specific: whether the factor’s own motif was more accessible in that population, assessed against a null in which motif profiles were shuffled between factors; whether the peaks linked to that regulon’s target genes were more accessible in that population than the peaks linked to size-matched random gene sets; and whether those peaks carried the factor’s own motif more often than other factors’ motifs, assessed by a paired Wilcoxon test. Factors passing all three tests are reported, and factors passing only one are not treated as validated.

### Compartment refits

Where a compartment was not separated by a whole-tissue contrast, the peak and motif models were refitted within that compartment alone, so that the compartment becomes the denominator. This was done for the synovial fibroblasts (lining against sublining, and subtypes within a layer) and for the IPFP adipogenic axis (the three adipocyte and five testable fibroblast subtypes). The fraction of a population’s whole-tissue enriched peaks retained under the refit is reported, because a population that loses nearly all of its enriched peaks is distinguished mainly from other lineages and little from related populations within its compartment.

### Cell–cell communication

Communication was inferred with CellChat^67,68^ from expressed ligand–receptor pairs, separately within each tissue. In each tissue’s own reference (Fig. 2f, Fig. 4f) the full human database was used with CellChat’s population-size weighting; in the joint reference the inference was restricted to protein ligand–receptor pairs (secreted, extracellular-matrix and cell–cell contact) and used no population-size term, so that interaction strength there does not simply reflect population size. Significance was assessed by CellChat’s permutation test over cell labels. Because the two tissues were dissected separately, no cross-tissue signaling was inferred and none is reported. Pathway differences were broken down by summing interaction strength per ligand, per ligand–receptor pair and per sending and receiving population within each tissue’s communication table, without repeating the inference. Donor robustness and a doublet-removal sensitivity analysis were run for the joint-reference results, and the one quantity that does not survive doublet removal is identified as such in the figure legend and is not reported.

### Joint reference and trajectory analysis

The twelve nuclei libraries were integrated together by canonical correlation analysis on the SCTransform-normalized data, with 3,000 integration features and 50 anchor dimensions, and clustered at resolution 0.3, chosen from a sweep of 0.1–2.0 by a combined silhouette and modularity score; this gave a joint reference of 54,716 nuclei and 16 populations, annotated from marker genes with the per-tissue labels of the same nuclei as a cross-check (at resolution 0.8 the same object gives 23 clusters). A trajectory was fitted over the 30,455 mesenchymal nuclei of the joint reference with Slingshot,^69^ rooted at the population scoring highest for a published DPP4+ progenitor signature, scored per nucleus as a module score against expression-matched control genes. The fit was repeated with Monocle3,^70,71^ which can place a branch anywhere along a principal graph, including between population centers, with each tissue refitted alone in its own embedding, and with every leave-one-donor-out subset. Genes and regulators changing along the axis were summarized by pseudotime decile, donor-averaged and standardized within tissue, and a feature was reported as peaking mid-axis only if at least four of six donors placed its maximum within one decile of the pooled maximum and no other third of the axis reached 90% of that maximum.

### Cell-cycle state and turnover programs

For each nucleus the number of detected S-phase and G2/M genes (Seurat cc.genes.updated.2019, 97 present)^72^ was compared with the number detected from 180 random gene sets matched on mean expression (25 bins) in the same nucleus, giving an empirical p and a z-score against that nucleus’s own null. Twenty further matched sets were held out as decoys and put through the identical test to measure the false-discovery proportion at each z threshold; the call was the least stringent threshold with a proportion of at most 10% (z ≥ 5 with p ≤ 0.01; measured 2.1%, 136 nuclei called against 2.8 expected by chance), additionally requiring Seurat phase S or G2/M. The genes separating called nuclei were led by MKI67, E2F8 and HJURP, and called nuclei carried E2F-target and homologous-recombination scores well above those of non-cycling nuclei. A cycling nucleus was attributed to a compartment only when its joint label and its label in the tissue’s own reference agreed, because 17 of the 18 cycling nuclei in the joint adipocyte cluster were immune cells by their native label. Rates are reported per donor with exact Poisson 95% intervals; zero counts carry the exact one-sided bound. Replacement times assumed an 8 to 16 h S/G2/M window, and the supply check compared the progenitor divisions per day required by 10% annual adipocyte renewal^26^ with the observed fibroblast rate, taking nucleus counts to be proportional to cell counts. Twenty-six MSigDB programs covering proliferation, quiescence, senescence, organelle turnover and DNA repair (Supplementary Data 6) were scored with AddModuleScore, residualized on log UMI and log detected genes across all nuclei, and summarized along each pseudotime trajectory with the donor-averaged decile statistic used for the trajectory genes.

### Comparison with published atlases

Published marker panels^18,19,21^ were scored against our populations as module scores against expression-matched control gene sets, at population level, and reciprocal best matches were required before two populations from different studies were said to nominate each other. Where differences in study design could account for a discrepancy with a published atlas, this is noted in the text.

### Software

Single-nucleus libraries were processed with Cell Ranger ARC 2.0.2 (10x Genomics; RRID:SCR_023897) against GRCh38, and bulk libraries with Trim Galore 0.6.10 (RRID:SCR_011847; Cutadapt 5.1, RRID:SCR_011841), HISAT2 2.2.1 (RRID:SCR_015530) and featureCounts (RRID:SCR_012919; Subread 2.1.1, RRID:SCR_009803). Analyses were performed in R 4.3.1 (RRID:SCR_001905) with Seurat 5.1.0 (RRID:SCR_016341; SeuratObject 5.0.2)^73^ and Signac 1.13.0 (RRID:SCR_021158)^74^ for single-nucleus processing, sctransform 0.4.1 (RRID:SCR_022146) with glmGamPoi 1.12.2 for normalization, scDblFinder 1.16.0 (RRID:SCR_022700) for doublet calling, harmony 1.2.0 (RRID:SCR_022206)^75^ for donor correction in the IPFP subclustering, limma 3.56.2 (RRID:SCR_010943) for donor-blocked linear models, chromVAR 1.24.0 (RRID:SCR_026570) with JASPAR2020 0.99.10 and motifmatchr 1.24.0 (RRID:SCR_026739) for motif accessibility, ChIPseeker 1.38.0 (RRID:SCR_021322) for peak annotation, CellChat 2.1.2 (RRID:SCR_021946) for communication inference, slingshot 2.10.0 (RRID:SCR_017012) and monocle3 1.3.7 (RRID:SCR_018685) for trajectories, AUCell 1.22.0 (RRID:SCR_021327) for regulon scoring, msigdbr 7.5.1 (RRID:SCR_022870) for gene sets, and DESeq2 1.40.2 (RRID:SCR_015687)^58^ and data.table 1.17.0 (RRID:SCR_026117) for peak differential accessibility and the bulk analyses, with pySCENIC 0.11.2 (RRID:SCR_025802) in Python 3.6 (RRID:SCR_008394). Random seeds in R were fixed.

### Subclustering of the IPFP adipocyte and stromal compartments

The IPFP reference was first annotated at a resolution giving fourteen populations, of which three were mesenchymal: adipocytes and two stromal labels. Because that resolution left the stroma coarser than the published IPFP atlases resolve it, and left the adipocytes undivided, we re-clustered the mesenchymal nuclei internally and asked whether the resulting subdivisions were supported.

The adipocyte and the two stromal labels were re-clustered in two arms. The two stromal labels were pooled before splitting, so the six resulting subtypes are not a refinement of one label each; three derive from each side of the original labels. Nuclei called as doublets were held out of the clustering and assigned afterwards by nearest-neighbor vote, so a doublet could not create a subcluster of its own.

Resolution was chosen by reproducibility across feature sets. Each arm was clustered twice over a range of resolutions: once in the embedding inherited from the whole-tissue integration, and once on variable genes selected within the arm alone, with Harmony correction on donor. The chosen resolution is the one at which the two independent feature sets recover the most similar partition of the same nuclei, measured by adjusted Rand index. Silhouette width and modularity were not used because neither identifies a number here: silhouette is highest at the coarsest adipocyte split (two clusters) and is flat near 0.10 (0.07–0.12) across every stromal resolution from six clusters to ten, and modularity rises monotonically without turning over (silhouette and cross-embedding agreement in Supplementary Fig. 33). With predicted doublets removed, as they were for clustering, the adipocyte arm reaches its maximum agreement at three subclusters (adjusted Rand index 0.53, against 0.36 at two), and the stromal arm does not reach the 0.50 threshold at any resolution that places at least two donor-supported subclusters, so it is reported as a descriptive subdivision (Supplementary Fig. 33d; subclusters in Supplementary Figs. 34 and 35).

Each subcluster was then required to be donor-supported (at least 25 nuclei in at least four of the six donors) and was tested for two artifacts not detected by clustering metrics. Sequencing depth was tested by equalizing every nucleus in an arm to a common UMI target and re-clustering from scratch, the depth effect being the adjusted Rand index between the full-depth and downsampled partitions of the same pipeline; and marker ubiquity was scored against the per-gene, per-cell-type detection rate computed over the whole fourteen-population reference, which is independent of this subclustering (Supplementary Fig. 24). Doublet enrichment was taken from the proportion of held-out predicted doublets each subcluster received.

Three subclusters failed one of these and were named so that they could not be mistaken for ordinary cell states: one adipocyte subcluster was defined by low library complexity and had no transcriptional program of its own, one stromal subcluster was both depth-suspect and doublet-enriched, and one was present at the required size in only three of the six donors. They were retained so that every nucleus in the reference carried a label, and were excluded from the analyses where they would distort a result; the trajectory was refitted as a sensitivity analysis with the flagged subtypes removed.

Population names were assigned by five rules: names describe only measured properties and do not imply potency, direction or lineage; every gene named in a label has the highest mean expression of any subtype in its arm; lncRNAs and clone identifiers are not used; functional descriptors are used only where supported by a measured program; and quality-limited populations are identified as such. The transcriptional programs and matrisome shares behind the descriptors are shown in Supplementary Fig. 36, and the chromatin test of the subdivision in Supplementary Fig. 23.

### Statistics and reproducibility

The donor was the unit of replication for every inferential test (see *Statistical framework*). The multiome cohort comprised six donors, each contributing both tissues, and the bulk cohort 14 synovium and 18 IPFP donors; no statistical method was used to predetermine sample size. Differential accessibility was tested with DESeq2 and motif activity with limma, both blocking on donor, with Benjamini–Hochberg correction; enrichment by one-sided hypergeometric test; cycling rates with exact Poisson 95% intervals; and cross-modal agreement against shuffled or size-matched random nulls and by paired Wilcoxon test. Trajectories were repeated with a second method, in each tissue alone and with every donor left out in turn. Random seeds in R were fixed.

### Use of AI tools

The analyses were built on analysis pipelines developed by the authors. An AI coding agent (Claude, Anthropic; accessed through Claude Code, April–November 2026) was used, under the authors’ direction, to refactor and extend these pipelines and to write the downstream analysis and figure scripts, to propose cell-type labels for clusters of the all-donor objects from their marker genes, which the authors reviewed and finalized, and to prepare a first draft of the manuscript text, which the authors then revised and edited over several rounds. The authors designed the analyses, selected methods and parameters, and interpreted the results. All code was run on the authors’ computers; all figures were generated by these scripts from the analysis output, and no generative image models were used. Numerical values were inserted programmatically from result tables. The authors take full responsibility for the content. All analysis code is available as described under Code availability.

### Data availability

Data analyzed in this study are derived from the healthy donors included in datasets that will be available through the SPARC Portal (RRID:SCR_017041) upon publication at the following DOIs: https://doi.org/10.26275/uhbq-r2s7, https://doi.org/10.26275/ifye-wvaq, https://doi.org/10.26275/t9lp-ovyr, and https://doi.org/10.26275/2sia-jzkd.43-46 Raw sequencing data (bulk RNA-seq and single-nucleus RNA-seq and ATAC-seq) will be available through dbGaP (RRID:SCR_002709) under accession phs004628. Tissue-processing protocols are available at protocols.io (https://doi.org/10.17504/protocols.io.ewov193molr2/v1; https://doi.org/10.17504/protocols.io.5qpvokqwxl4o/v1).^47,48^ Processed results supporting the main findings are provided as Supplementary Data 1–4 and 6, and the SPARC subject identifier of each donor in Supplementary Data 5.

## Acknowledgements

The authors thank Michelle Payad, Zaira Flores, Jessica Nitta, and the team at Lifesharing, San Diego for providing access to the precious gift of tissue donation from the local San Diego & Imperial communities that made the research possible. We thank the Scripps Research Genomics Core (RRID:SCR_029158) members Tony Mondala, Stephen Head and Sean Wang for their support with omics data generation. Our conclusions and discussion on this topic are influenced by our role on the RE-JOIN Consortium and by the active engagement of this team.

This work was supported by the National Institute of Arthritis and Musculoskeletal and Skin Diseases (NIAMS) of the National Institutes of Health under Award Number UC2 AR082186, as part of the Restoring Joint Health and Function to Reduce Pain (RE-JOIN; RRID:SCR_023514) consortium of the NIH Helping to End Addiction Long-term (HEAL) Initiative (RRID:SCR_022808) and by the National Institute on Aging under Award Number R01AG049617 (To ML). This paper’s content is solely the responsibility of the authors and does not necessarily represent the official views of the National Institutes of Health.

## Author contributions

M.K.L. and H.S. conceptualized this study. H.S. and M.K.L. drafted the manuscript, with contributions from the other authors. H.S. performed data analysis. All authors approved the final version of the manuscript and figures.

## The RE-JOIN Consortium

Armen Akopian, Kyle Allen, Alejandro Almarza, Benjamin Arenkiel, Yangjin Bae, Bruna Balbino de Paula, Anita Bandrowski, Mario Danilo Boada, Jacqueline Boccanfuso, Jyl Boline, Dawen Cai, Carpio, Dellina Lane, Robert Caudle, Racel Cela, Yong Chen, Rui Chen, Brian Constantinescu, Cortez, Ibdanelo, Yenisel Cruz-Almeida, M. Franklin Dolwick, Chris Donnelly, Zelong Dou, Joshua Emrick, Malin Ernberg, Danielle Freburg-Hoffmeister, Spencer Fullam, Janak Gaire, Akash Gandhi, Benjamin Goolsby, Stacey Greene, Nele Haelterman, Michael Iadarola, Shingo Ishihara, Azeez Ishola, Sudhish Jayachandran, Zixue Jin, Frank Ko, Priya Kulkarni, Zhao Lai, Brendan Lee, Yona Levites, Carolina Leynes, Jun Li, Martin Lotz, Lindsey Macpherson, Tristan Maerz, Camilla Majano, Anne-Marie Malfait, Maryann Martone, Bella Mehta, Richard Miller, Rachel Miller, Michael Newton, Alia Obeidat, Merissa Olmer, Dana Orange, Miguel Otero, Kevin Otto, Folly Patterson, Marlena Pela, Sienna Perry, Theodore Price, Hernan Prieto, Russell Ray, Dongjun Ren, Margarete Ribeiro Dasilva, Alexus Roberts, Elizabeth Ronan, Oscar Ruiz, Shad Smith, Mairobys Socorro, Kaitlin Southern, Joshua Stover, Michael Strinden, Hannah Swahn, Evelyne Tantry, Sue Tappan, Luis Tovias Sanchez, Airam Vivanco-Estela, Joost Wagenaar, Lai Wang, Kim Worley, Joshua Wythe, Jiansen Yan, and Julia Younis.

## References

1. GBD 2021 Osteoarthritis Collaborators. Global, regional, and national burden of osteoarthritis, 1990-2020 and projections to 2050: a systematic analysis for the Global Burden of Disease Study 2021. Lancet Rheumatol. 5, e508–e522 (2023).

2. Hunter, D. J. & Bierma-Zeinstra, S. Osteoarthritis. Lancet 393, 1745–1759 (2019).

3. Roemer, F. W. et al. Presence of MRI-detected joint effusion and synovitis increases the risk of cartilage loss in knees without osteoarthritis at 30-month follow-up: the MOST study. Ann. Rheum. Dis. 70, 1804–1809 (2011).

4. Atukorala, I. et al. Synovitis in knee osteoarthritis: a precursor of disease? Ann. Rheum. Dis. 75, 390–395 (2016).

5. Klein-Wieringa, I. R. et al. The infrapatellar fat pad of patients with osteoarthritis has an inflammatory phenotype. Ann. Rheum. Dis. 70, 851–857 (2011).

6. Bastiaansen-Jenniskens, Y. M. et al. Stimulation of fibrotic processes by the infrapatellar fat pad in cultured synoviocytes from patients with osteoarthritis: a possible role for prostaglandin F2α. Arthritis Rheum. 65, 2070–2080 (2013).

7. Ioan-Facsinay, A. & Kloppenburg, M. An emerging player in knee osteoarthritis: the infrapatellar fat pad. Arthritis Res. Ther. 15, 225 (2013).

8. Bohnsack, M. et al. Distribution of substance-P nerves inside the infrapatellar fat pad and the adjacent synovial tissue: a neurohistological approach to anterior knee pain syndrome. Arch. Orthop. Trauma Surg. 125, 592–597 (2005).

9. Belluzzi, E. et al. Contribution of Infrapatellar Fat Pad and Synovial Membrane to Knee Osteoarthritis Pain. Biomed. Res. Int. 2019, 6390182 (2019).

10. Mizoguchi, F. et al. Functionally distinct disease-associated fibroblast subsets in rheumatoid arthritis. Nat. Commun. 9, 789 (2018).

11. Croft, A. P. et al. Distinct fibroblast subsets drive inflammation and damage in arthritis. Nature 570, 246–251 (2019).

12. Culemann, S. et al. Locally renewing resident synovial macrophages provide a protective barrier for the joint. Nature 572, 670–675 (2019).

13. Wei, K. et al. Notch signalling drives synovial fibroblast identity and arthritis pathology. Nature 582, 259–264 (2020).

14. Zhang, F. et al. Defining inflammatory cell states in rheumatoid arthritis joint synovial tissues by integrating single-cell transcriptomics and mass cytometry. Nat. Immunol. 20, 928–942 (2019).

15. Zhang, F. et al. Deconstruction of rheumatoid arthritis synovium defines inflammatory subtypes. Nature 623, 616–624 (2023).

16. Chou, C. H. et al. Synovial cell cross-talk with cartilage plays a major role in the pathogenesis of osteoarthritis. Sci. Rep. 10, 10868 (2020).

17. Knights, A. J. et al. Synovial fibroblasts assume distinct functional identities and secrete R-spondin 2 in osteoarthritis. Ann. Rheum. Dis. 82, 272–282 (2023).

18. Tang, S. et al. Single-cell atlas of human infrapatellar fat pad and synovium implicates APOE signaling in osteoarthritis pathology. Sci. Transl. Med. 16, eadf4590 (2024).

19. Peters, H. et al. Cell and transcriptomic diversity of infrapatellar fat pad during knee osteoarthritis. Ann. Rheum. Dis. 84, 351–367 (2025).

20. Sárvári, A. K. et al. Plasticity of Epididymal Adipose Tissue in Response to Diet-Induced Obesity at Single-Nucleus Resolution. Cell Metab. 33, 437–453.e5 (2021).

21. Emont, M. P. et al. A single-cell atlas of human and mouse white adipose tissue. Nature 603, 926–933 (2022).

22. Satpathy, A. T. et al. Massively parallel single-cell chromatin landscapes of human immune cell development and intratumoral T cell exhaustion. Nat. Biotechnol. 37, 925–936 (2019).

23. Ma, S. et al. Chromatin Potential Identified by Shared Single-Cell Profiling of RNA and Chromatin. Cell 183, 1103–1116.e20 (2020).

24. Naba, A. et al. The matrisome: in silico definition and in vivo characterization by proteomics of normal and tumor extracellular matrices. Mol. Cell. Proteomics 11, M111.014647 (2012).

25 . Li, J. et al. Synovium and infrapatellar fat pad share common mesenchymal progenitors and undergo coordinated changes in osteoarthritis. J. Bone Miner. Res. 39, 161–176 (2024).

26 . Spalding, K. L. et al. Dynamics of fat cell turnover in humans. Nature 453, 783–787 (2008).

27 . Fridman, A. L. & Tainsky, M. A. Critical pathways in cellular senescence and immortalization revealed by gene expression profiling. Oncogene 27, 5975–5987 (2008).

28. Rhee, D. K. et al. The secreted glycoprotein lubricin protects cartilage surfaces and inhibits synovial cell overgrowth. J. Clin. Invest. 115, 622–631 (2005).

29. Walter, P. & Ron, D. The unfolded protein response: from stress pathway to homeostatic regulation. Science 334, 1081–1086 (2011).

30. Yu, W. H., Yu, S., Meng, Q., Brew, K. & Woessner, J. F. Jr. TIMP-3 binds to sulfated glycosaminoglycans of the extracellular matrix. J. Biol. Chem. 275, 31226–31232 (2000).

31. Holmbeck, K. et al. MT1-MMP-deficient mice develop dwarfism, osteopenia, arthritis, and connective tissue disease due to inadequate collagen turnover. Cell 99, 81–92 (1999).

32. Sahebjam, S., Khokha, R. & Mort, J. S. Increased collagen and aggrecan degradation with age in the joints of Timp3(-/-) mice. Arthritis Rheum. 56, 905–909 (2007).

33. Coller, H. A., Sang, L. & Roberts, J. M. A new description of cellular quiescence. PLoS Biol. 4, e83 (2006).

34. Roelofs, A. J. et al. Joint morphogenetic cells in the adult mammalian synovium. Nat. Commun. 8, 15040 (2017).

35. Collins, F. L. et al. Taxonomy of fibroblasts and progenitors in the synovial joint at single-cell resolution. Ann. Rheum. Dis. 82, 428–437 (2023).

36. Roelofs, A. J., McClure, J. J., Hay, E. A. & De Bari, C. Stem and progenitor cells in the synovial joint as targets for regenerative therapy. Nat. Rev. Rheumatol. 21, 211–220 (2025).

37. Rosen, E. D. & MacDougald, O. A. Adipocyte differentiation from the inside out. Nat. Rev. Mol. Cell Biol. 7, 885–896 (2006).

38. Merrick, D. et al. Identification of a mesenchymal progenitor cell hierarchy in adipose tissue. Science 364, eaav2501 (2019).

39. Schwalie, P. C. et al. A stromal cell population that inhibits adipogenesis in mammalian fat depots. Nature 559, 103–108 (2018).

40. Bakken, T. E. et al. Single-nucleus and single-cell transcriptomes compared in matched cortical cell types. PLoS ONE 13, e0209648 (2018).

41. Wu, H., Kirita, Y., Donnelly, E. L. & Humphreys, B. D. Advantages of Single-Nucleus over Single-Cell RNA Sequencing of Adult Kidney: Rare Cell Types and Novel Cell States Revealed in Fibrosis. J. Am. Soc. Nephrol. 30, 23–32 (2019).

42. Outerbridge, R. E. The etiology of chondromalacia patellae. J. Bone Joint Surg. Br. 43-B, 752–757 (1961).

43. Swahn, H., Olmer, M. & Lotz, M. Bulk RNA-seq profiling of the human knee medial synovium (Version 1). SPARC Portal 10.26275/uhbq-r2s7 (2026).

44. Swahn, H., Olmer, M. & Lotz, M. Single-nucleus RNA-seq and ATAC-seq profiling of the human knee medial synovium (Version 1). SPARC Portal 10.26275/ifye-wvaq (2026).

45. Swahn, H., Olmer, M. & Lotz, M. Bulk RNA-seq profiling of the human knee infrapatellar fat pad (Version 1). SPARC Portal 10.26275/t9lp-ovyr (2026).

46. Swahn, H., Olmer, M. & Lotz, M. Single-nucleus RNA-seq and ATAC-seq profiling of the human knee infrapatellar fat pad (Version 1). SPARC Portal 10.26275/2sia-jzkd (2026).

47. Lorenzo Gomez, I., Wang, E., Olmer, M. & Lotz, M. Nuclei Isolation from Human Synovium and Infrapatellar Fat Pad for 10x Multiome (Version 1). *protocols.io* 10.17504/protocols.io.ewov193molr2/v1 (2024).

48. Lorenzo Gomez, I., Olmer, M. & Lotz, M. RNA Isolation of Human Synovium and Fat Pad Tissues (Version 1). *protocols.io* 10.17504/protocols.io.5qpvokqwxl4o/v1 (2024).

49. Martin, M. Cutadapt removes adapter sequences from high-throughput sequencing reads. EMBnet.journal 17, 10–12 (2011).

50. Kim, D., Paggi, J. M., Park, C., Bennett, C. & Salzberg, S. L. Graph-based genome alignment and genotyping with HISAT2 and HISAT-genotype. Nat. Biotechnol. 37, 907–915 (2019).

51. Liao, Y., Smyth, G. K. & Shi, W. featureCounts: an efficient general purpose program for assigning sequence reads to genomic features. Bioinformatics 30, 923–930 (2014).

52. Germain, P. L., Lun, A., Garcia Meixide, C., Macnair, W. & Robinson, M. D. Doublet identification in single-cell sequencing data using scDblFinder. F1000Res. 10, 979 (2021).

53. Hafemeister, C. & Satija, R. Normalization and variance stabilization of single-cell RNA-seq data using regularized negative binomial regression. Genome Biol. 20, 296 (2019).

54. Choudhary, S. & Satija, R. Comparison and evaluation of statistical error models for scRNA-seq. Genome Biol. 23, 27 (2022).

55. Ahlmann-Eltze, C. & Huber, W. glmGamPoi: fitting Gamma-Poisson generalized linear models on single cell count data. Bioinformatics 36, 5701–5702 (2021).

56. Stuart, T. et al. Comprehensive integration of single-cell data. Cell 177, 1888–1902.e21 (2019).

57. Hao, Y. et al. Integrated analysis of multimodal single-cell data. Cell 184, 3573–3587.e29 (2021).

58. Love, M. I., Huber, W. & Anders, S. Moderated estimation of fold change and dispersion for RNA-seq data with DESeq2. Genome Biol. 15, 550 (2014).

59. Ritchie, M. E. et al. limma powers differential expression analyses for RNA-sequencing and microarray studies. Nucleic Acids Res. 43, e47 (2015).

60. Ashburner, M. et al. Gene ontology: tool for the unification of biology. Nat. Genet. 25, 25–29 (2000).

61. Liberzon, A. et al. The Molecular Signatures Database (MSigDB) hallmark gene set collection. Cell Syst. 1, 417–425 (2015).

62. Aibar, S. et al. SCENIC: single-cell regulatory network inference and clustering. Nat. Methods 14, 1083–1086 (2017).

63. Van de Sande, B. et al. A scalable SCENIC workflow for single-cell gene regulatory network analysis. Nat. Protoc. 15, 2247–2276 (2020).

64. Yu, G., Wang, L. G. & He, Q. Y. ChIPseeker: an R/Bioconductor package for ChIP peak annotation, comparison and visualization. Bioinformatics 31, 2382–2383 (2015).

65. Fornes, O., et al. JASPAR 2020: update of the open-access database of transcription factor binding profiles. Nucleic Acids Res. 48, D87–D92 (2020).

66. Schep, A. N., Wu, B., Buenrostro, J. D. & Greenleaf, W. J. chromVAR: inferring transcription-factor-associated accessibility from single-cell epigenomic data. Nat. Methods 14, 975–978 (2017).

67. Jin, S. et al. Inference and analysis of cell-cell communication using CellChat. Nat. Commun. 12, 1088 (2021).

68. Jin, S., Plikus, M. V. & Nie, Q. CellChat for systematic analysis of cell–cell communication from single-cell transcriptomics. Nat. Protoc. 20, 180–219 (2025).

69. Street, K. et al. Slingshot: cell lineage and pseudotime inference for single-cell transcriptomics. BMC Genomics 19, 477 (2018).

70. Trapnell, C. et al. The dynamics and regulators of cell fate decisions are revealed by pseudotemporal ordering of single cells. Nat. Biotechnol. 32, 381–386 (2014).

71. Cao, J. et al. The single-cell transcriptional landscape of mammalian organogenesis. Nature 566, 496–502 (2019).

72. Tirosh, I. et al. Dissecting the multicellular ecosystem of metastatic melanoma by single-cell RNA-seq. Science 352, 189–196 (2016).

73. Hao, Y. et al. Dictionary learning for integrative, multimodal and scalable single-cell analysis. Nat. Biotechnol. 42, 293–304 (2024).

74. Stuart, T., Srivastava, A., Madad, S., Lareau, C. A. & Satija, R. Single-cell chromatin state analysis with Signac. Nat. Methods 18, 1333–1341 (2021).

75. Korsunsky, I. et al. Fast, sensitive and accurate integration of single-cell data with Harmony. Nat. Methods 16, 1289–1296 (2019).

