## Supplementary Information for "Paired multiomic profiling defines the healthy human synovium–infrapatellar fat pad stromal continuum"

**Supplementary Figures**

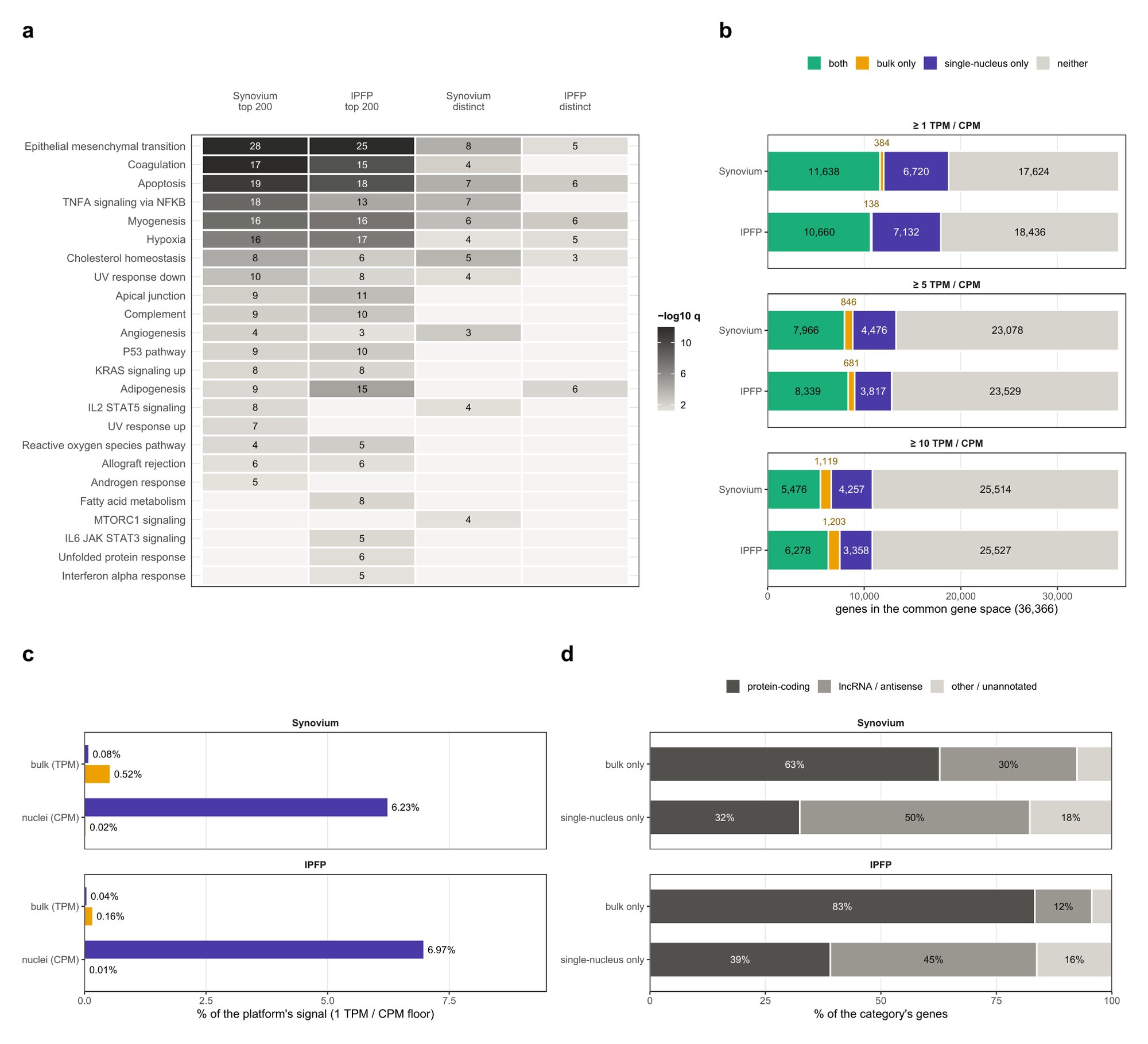

**Supplementary Fig. 1 | Bulk RNA-seq against the single-nucleus data, and the function of the most abundant transcripts.** **a** Hallmark gene sets over-represented among the 200 most abundant protein-coding transcripts of each tissue, excluding mitochondrial-encoded and ribosomal protein genes, and among the part of each list that is not in the other tissue's top 200 (distinct). Numbers, genes of the list in the set; shading, −log10 q from a one-sided hypergeometric test against the protein-coding genes detected in that tissue, Benjamini–Hochberg corrected; blank, q ≥ 0.05 or no gene of the list in the set. GO Biological Process results are in Supplementary Data 1. **b** The 36,366 genes annotated in both references, by whether bulk and single nuclei detect them, at 1, 5 and 10 TPM (bulk) or CPM (single-nucleus tissue pseudobulk); a gene counts as detected when it reaches the floor in at least half of that platform's donors. Counts of the thin bulk-only segments are printed above the bar. **c** Share of each platform's total signal carried by the bulk-only and single-nucleus-only genes, at the 1 TPM or CPM floor; colors as in b. **d** Gene class of the bulk-only and single-nucleus-only genes. n = 14 synovium and 18 IPFP bulk donors; six single-nucleus donors per tissue.

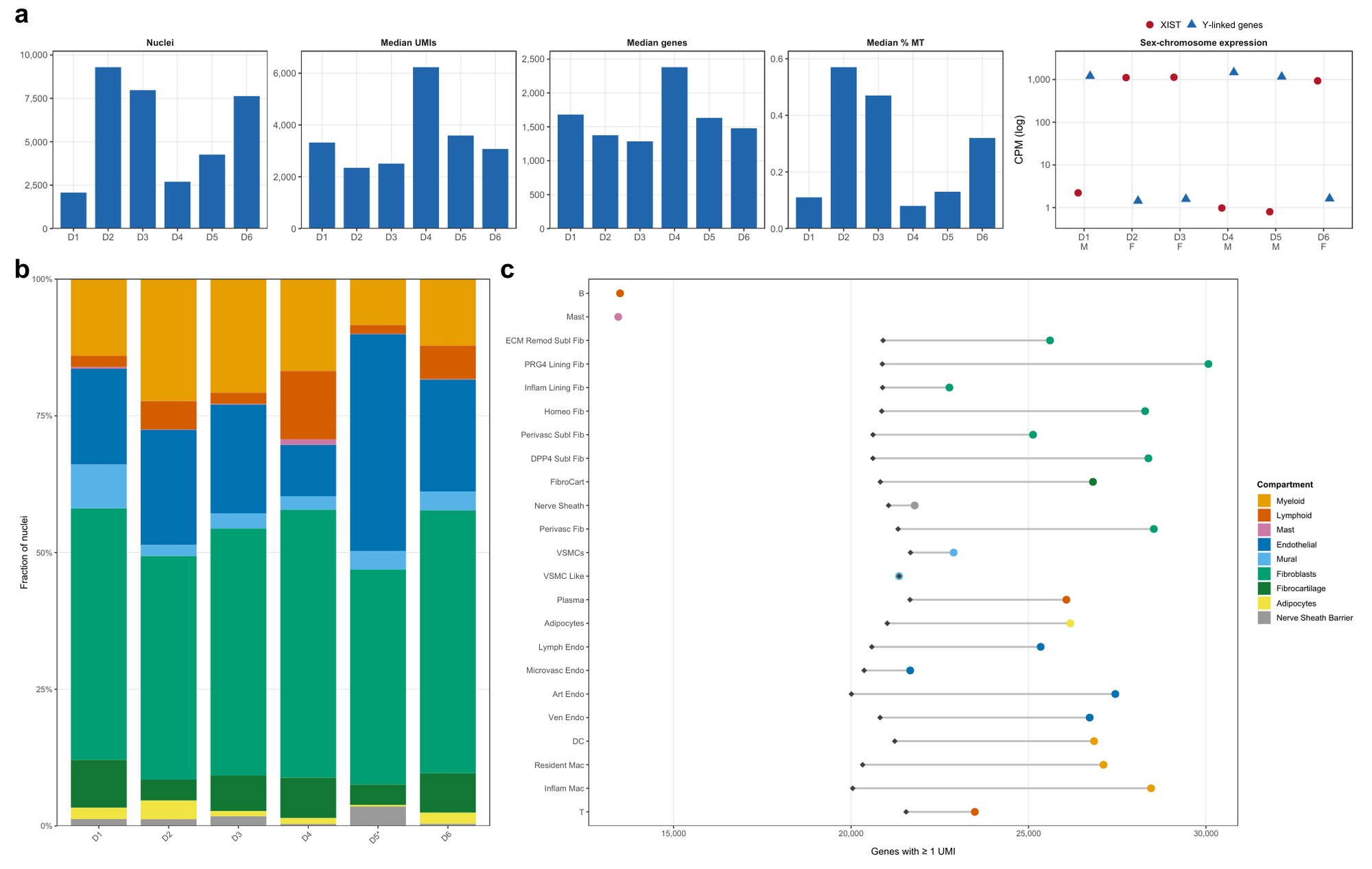

**Supplementary Fig. 2 | Cohort, sequencing quality, composition and gene detection (synovium).** **a** Per-donor nuclei, median UMIs, median genes and median mitochondrial percentage; donors are labeled D1–D6 in cohort order. Right, sex-chromosome expression in each donor's pseudobulk (counts per million, log scale): XIST and the summed Y-linked genes RPS4Y1, DDX3Y, UTY, USP9Y, EIF1AY, KDM5D, ZFY, NLGN4Y and TMSB4Y, with the recorded sex under each donor. Expression matched the recorded sex in every donor, in both synovium and IPFP (IPFP, Supplementary Fig. 13a). **b** Compartment composition per donor. Fibroblasts are the most stable compartment at 39–49% and endothelium the least at 9–40%; the starred donor carries 40% endothelial nuclei against a median near 20%. **c** Genes detected per population as sequenced (circles) and after equalizing depth by analytic subsampling to the shallowest abundance-filtered population (diamonds). B cells and mast cells never reach that depth and carry no matched point; once depth is equalized the 21 populations detect within about 8% of each other. Population labels in Supplementary Figs. 2–12 are abbreviated (for example, ECM Remod Subl Fib, ECM-remodeling sublining fibroblast; Homeo Fib, homeostatic fibroblast).

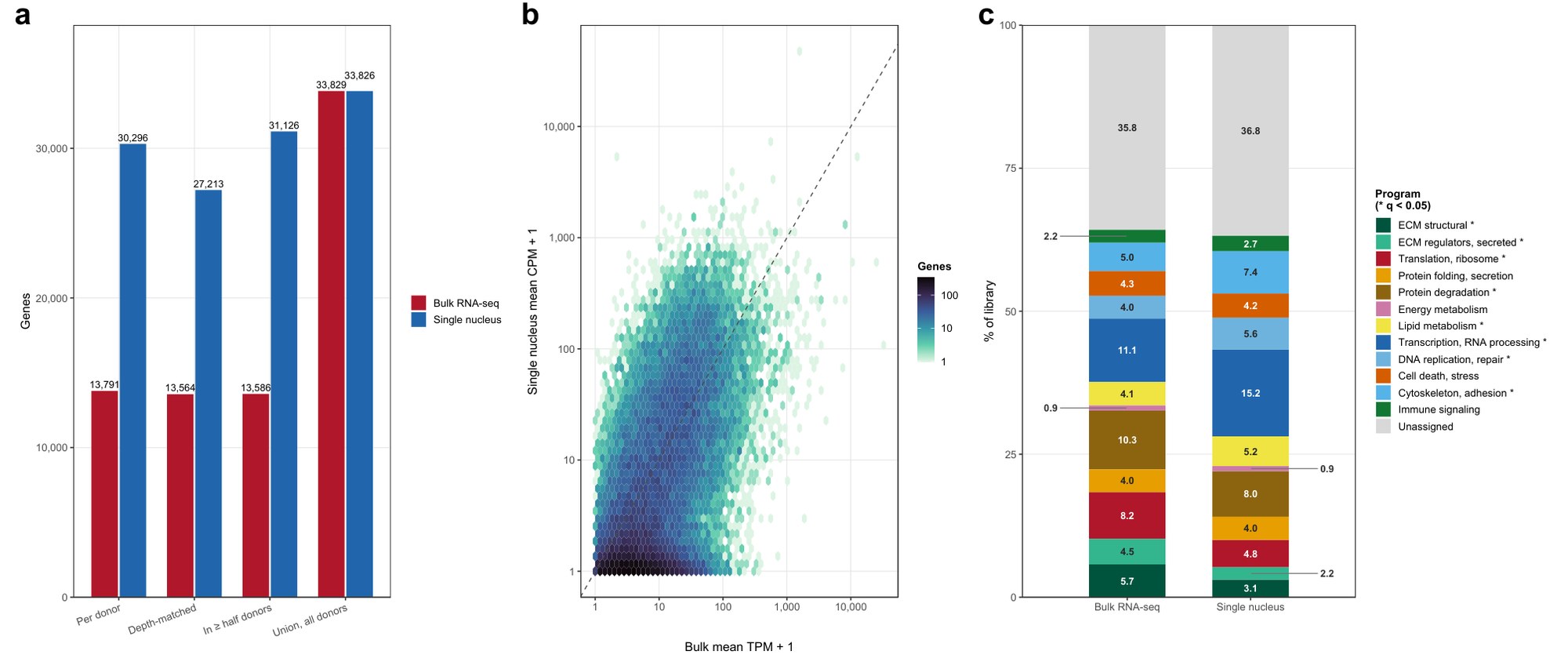

**Supplementary Fig. 3 | The healthy synovial reference against bulk RNA-seq of the same tissue.** **a** Genes detected on each platform in the common gene space. **b** Gene-level concordance, Spearman ρ = 0.51 over 32,027 shared genes. **c** The twelve programs of Fig. 2c for the whole tissue on each platform; 8 of 12, marked with asterisks, differ at q < 0.05 (Welch t-test on donor-level shares, Benjamini–Hochberg adjusted). Five of the six single-nucleus donors have a bulk synovium library (four from the same knee), and donor-matched pairs agree no better than mismatched ones (mean ρ = 0.499 against 0.495).

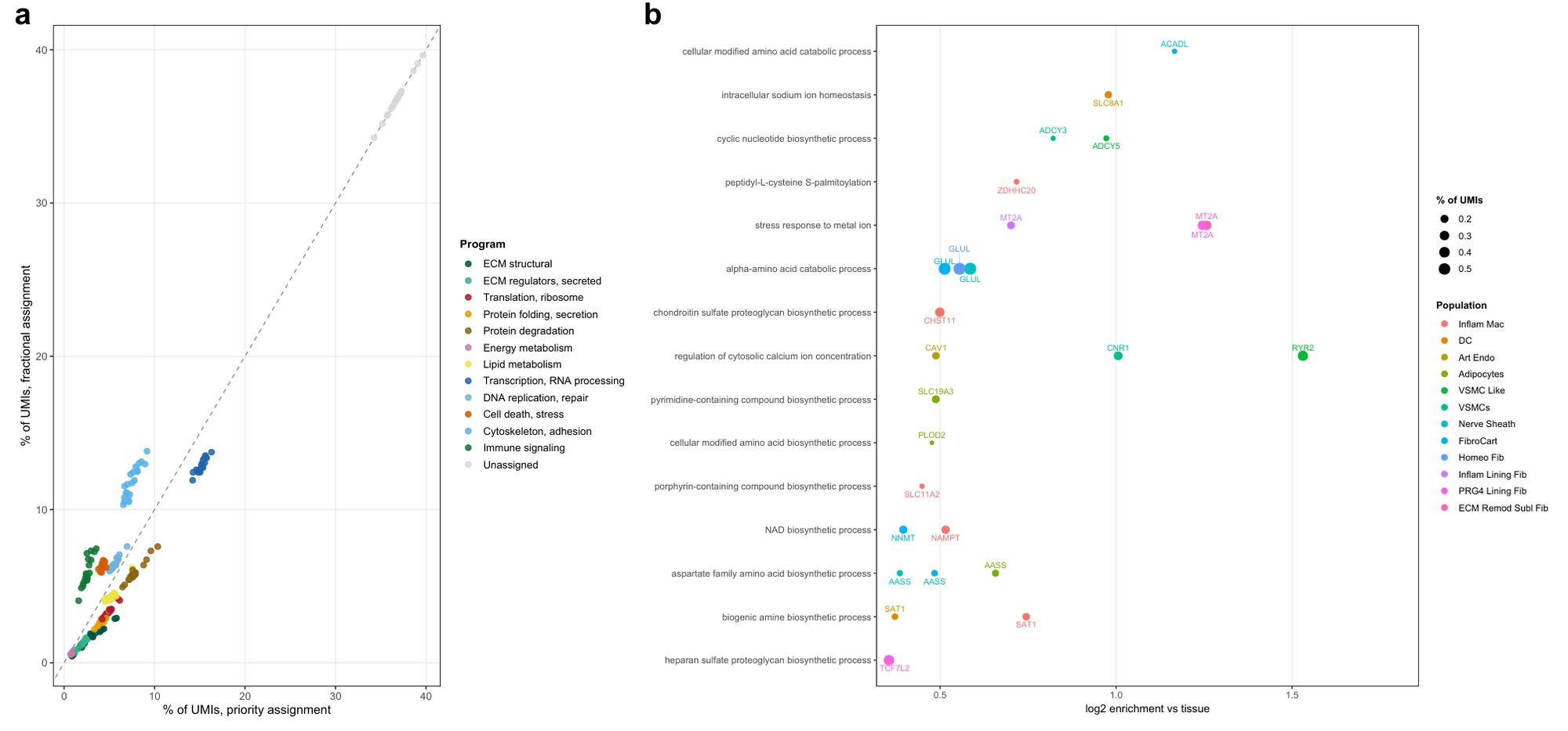

**Supplementary Fig. 4 | Robustness and coverage of the twelve-program transcriptional partition (synovium).** **a** Every program share recomputed under fractional assignment, where a multi-program gene's counts split equally, against the priority assignment used in Fig. 2c; r = 0.976. **b** An unbiased scan of gene ontology biological-process terms within a size window, restricted to terms the twelve programs do not cover, top gene labeled. The strongest hit is regulation of cytosolic calcium in VSMC-like cells (RYR2), followed by a metallothionein and zinc-stress response in the PRG4 lining and ECM-remodeling sublining fibroblasts (MT2A).

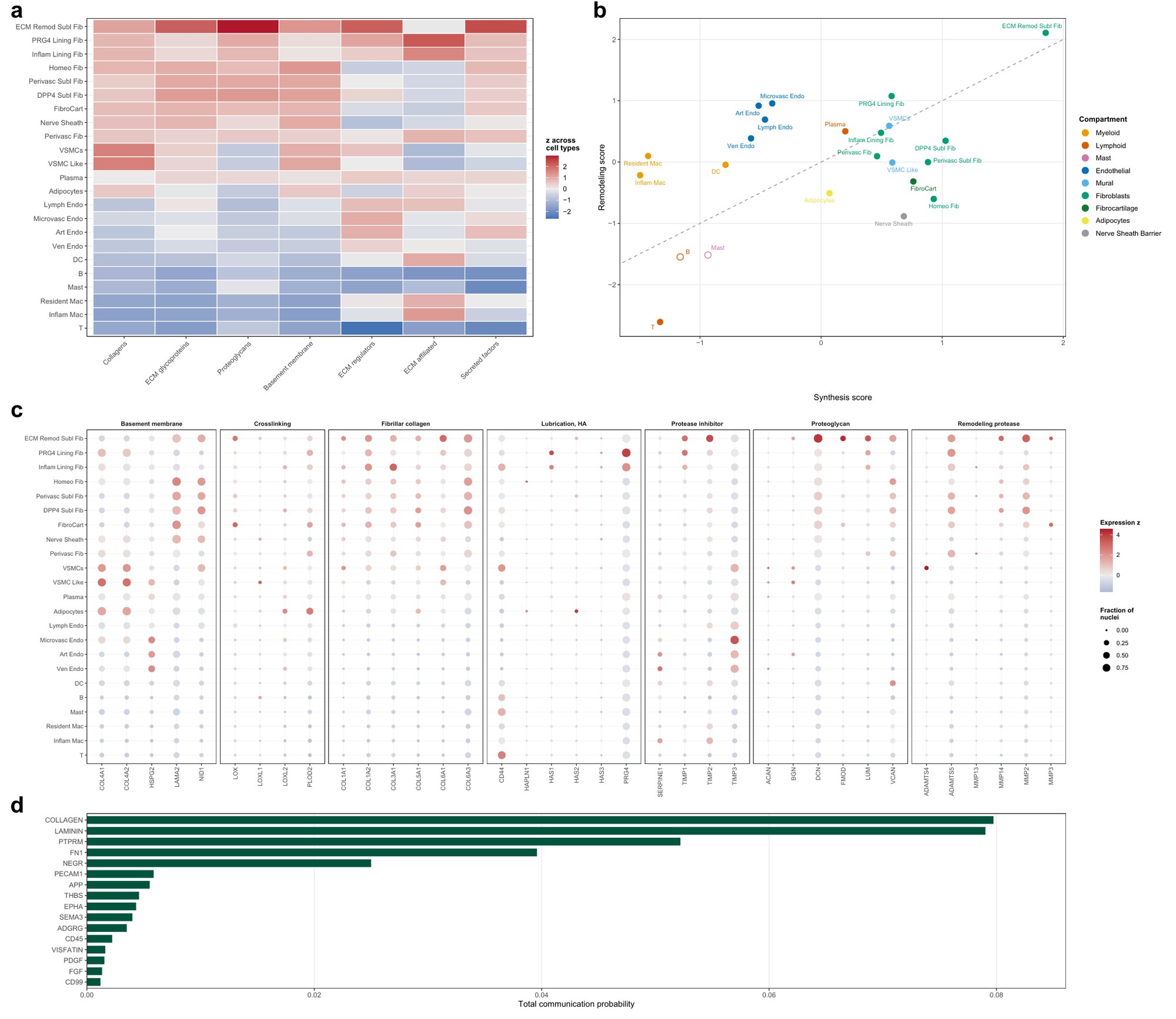

**Supplementary Fig. 5 | Matrisome composition, the synthesis–remodeling balance and predicted matrix signaling (synovium).** **a** Naba matrisome divisions per population, share of counts z-scored across populations. **b** Structural synthesis against remodeling output; above the line is remodeling-dominant, open points where fewer than three donors clear the nucleus floor. **c** Canonical matrisome markers; dot size is the fraction of nuclei with a non-zero count. **d** The strongest inferred signaling pathways (CellChat, all pathways), led by collagen and laminin.

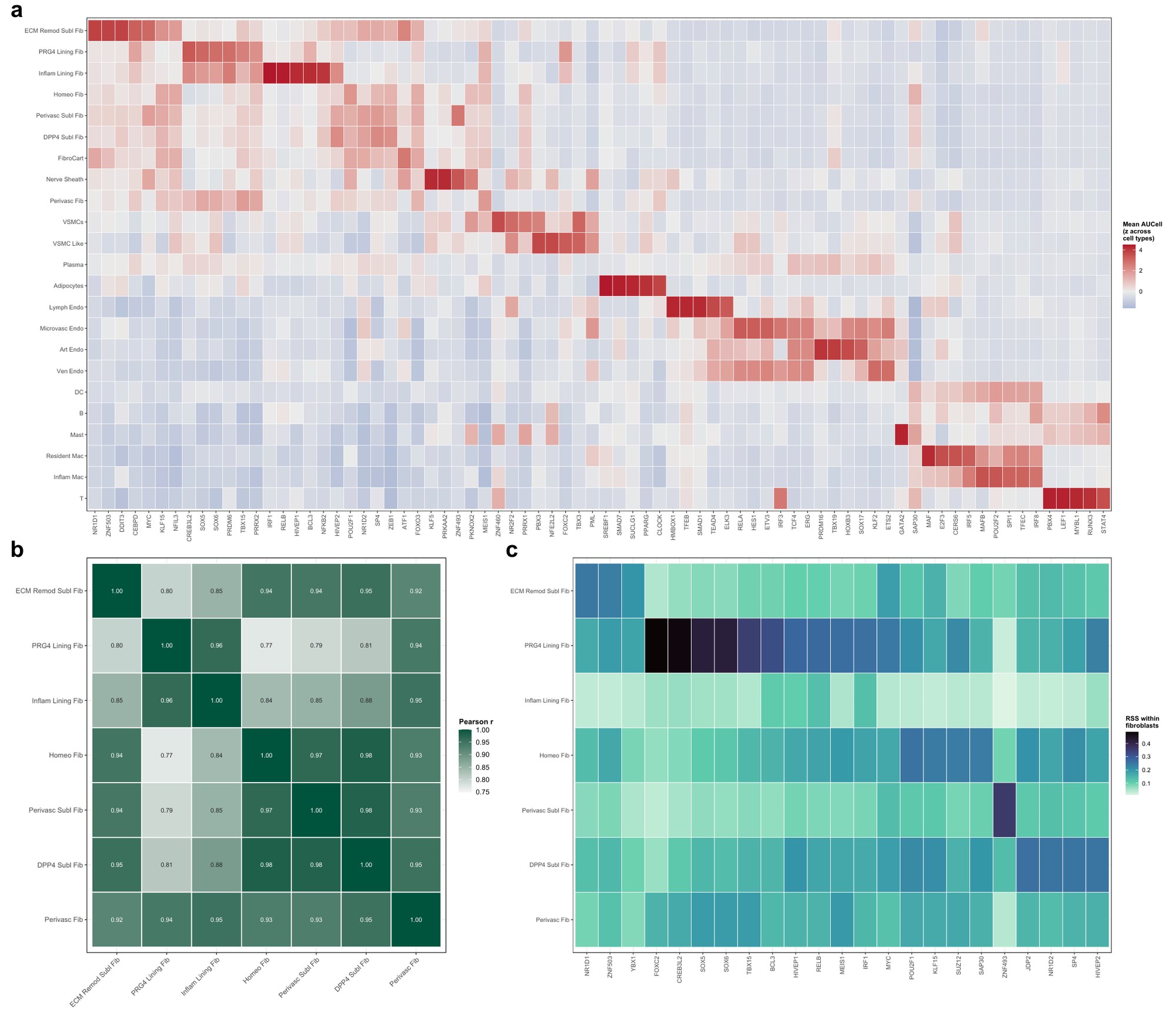

**Supplementary Fig. 6 | Regulon activity of the synovial populations.** pySCENIC was run on healthy nuclei only, without our cell-type labels. **a** The five most specific regulons per population, ordered by the population they peak in; canonical lineage factors are recovered unprompted, including PPARG in adipocytes, SOX17 in arterial and KLF2 in venous endothelium, MAF in resident macrophages and LEF1 in T cells. **b** Fibroblast subtype similarity on regulon activity; median off-diagonal r = 0.93. PRG4 lining fibroblasts are only modestly set apart on this readout: they correlate with the other subtypes from 0.77 to 0.96, the four lowest values in the matrix, and above the median against perivascular (0.94) and inflammatory lining (0.96) fibroblasts. **c** Specificity recomputed within the fibroblast compartment only, where they do stand apart, carrying the highest within-compartment specificity of any subtype (FOXC2 0.48, CREB3L2 0.48, SOX5 0.42).

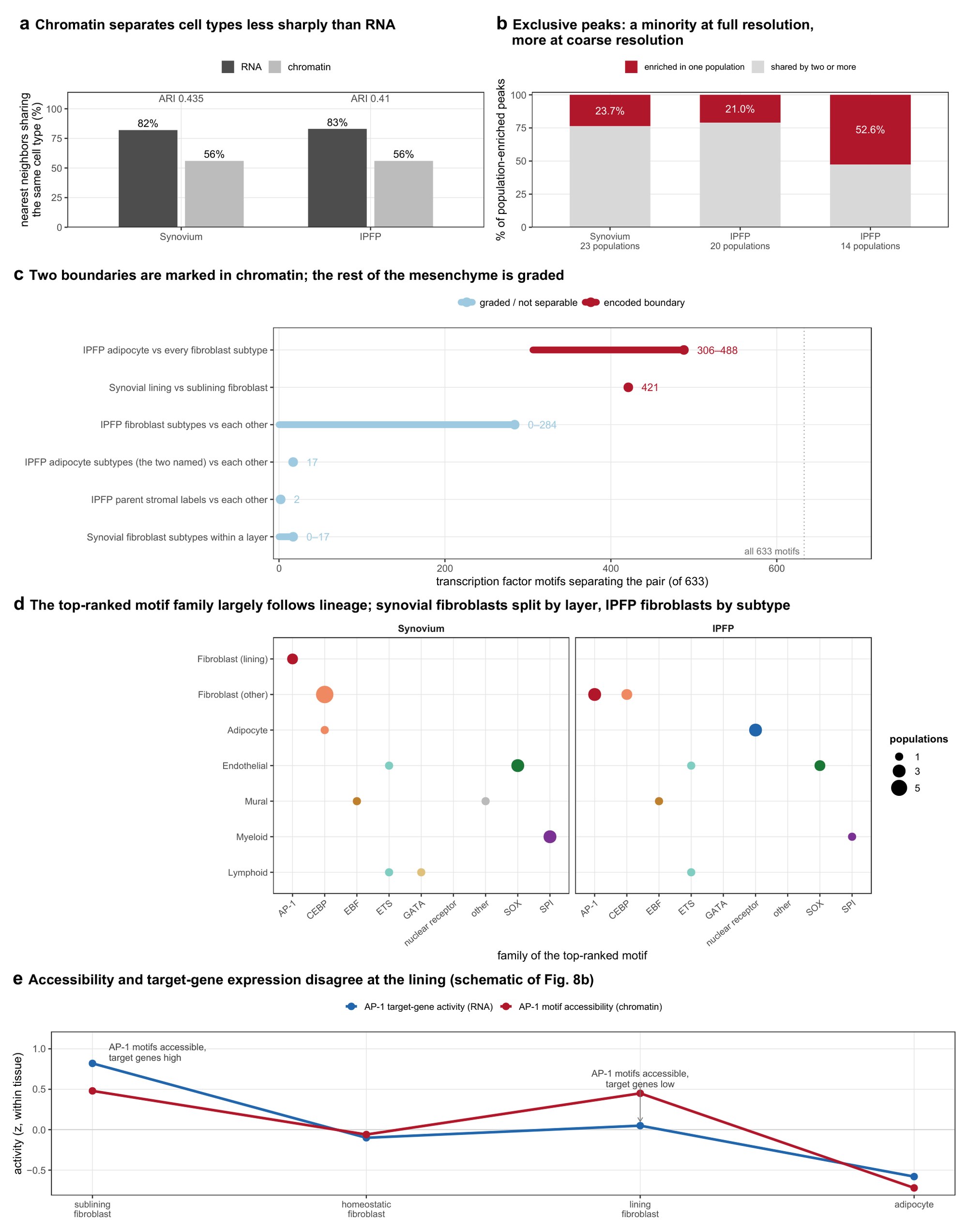

**Supplementary Fig. 7 | What the chromatin data show, across both tissues.** A summary of the four chromatin sections of the Results. Only panel e is a schematic. **a** Agreement between a nucleus's nearest neighbors and its own cell type, computed separately in RNA and in chromatin for each tissue, with the adjusted Rand index of a chromatin-only clustering against the transcriptomic labels given above each pair. Chromatin assigns cell type less sharply than RNA in both tissues. **b** Population-enriched peaks split into those enriched in exactly one population and those shared by two or more. The third bar repeats the IPFP test over the 14 coarser populations rather than the 20 subtypes, the comparison showing that peak exclusivity depends on cluster resolution. **c** Motifs separating each pair of populations, out of 633, from the compartment refits, so that the denominator is the compartment and not the whole tissue. Ranges are drawn where the contrast covers several pairs. Two contrasts separate most strongly; the remaining four, all within the mesenchyme, are graded or not separable at all, although single IPFP fibroblast pairs reach 284 motifs. Compared pairwise in the fibroblast-only model, the three synovial sublining subtypes differ at no motif and the two lining subtypes at 17. Drawn from the refits of Figs. 3f and 5f, these within-layer contrasts and Supplementary Fig. 23. **d** The family of the top-ranked motif in every population that met the testing threshold and belongs to a drawn lineage (the nerve-sheath barrier population, top motif NR3C2, has no lineage row), by lineage and tissue; point size is the number of populations of that lineage whose top-ranked motif belongs to that family. The family largely follows lineage. The fibroblasts split: in synovium the two lining subtypes rank AP-1 first and the other five subtypes and fibrocartilage CEBP, while in the IPFP three subtypes (adhesion, CXCL14+/IGFBP7+ and matrix-high) rank AP-1 first and two (SMOC1+/GHR+ and lipid-laden) CEBP. Endothelial populations split between SOX and ETS families, and in synovium the mural (EBF, SRF) and lymphoid (ETS in T cells, GATA in plasma cells) populations also split. Assignments name a family and not an individual protein, because motif accessibility is shared within a family. **e** Schematic of the two AP-1 tracks of Fig. 8b, on a common z-score axis: target-gene activity is highest at the root and falls, whereas motif accessibility returns to near its root height at the lining fibroblast. At the lining the AP-1 motifs are accessible but the AP-1 target genes are expressed at a low level; at the sublining both are high.

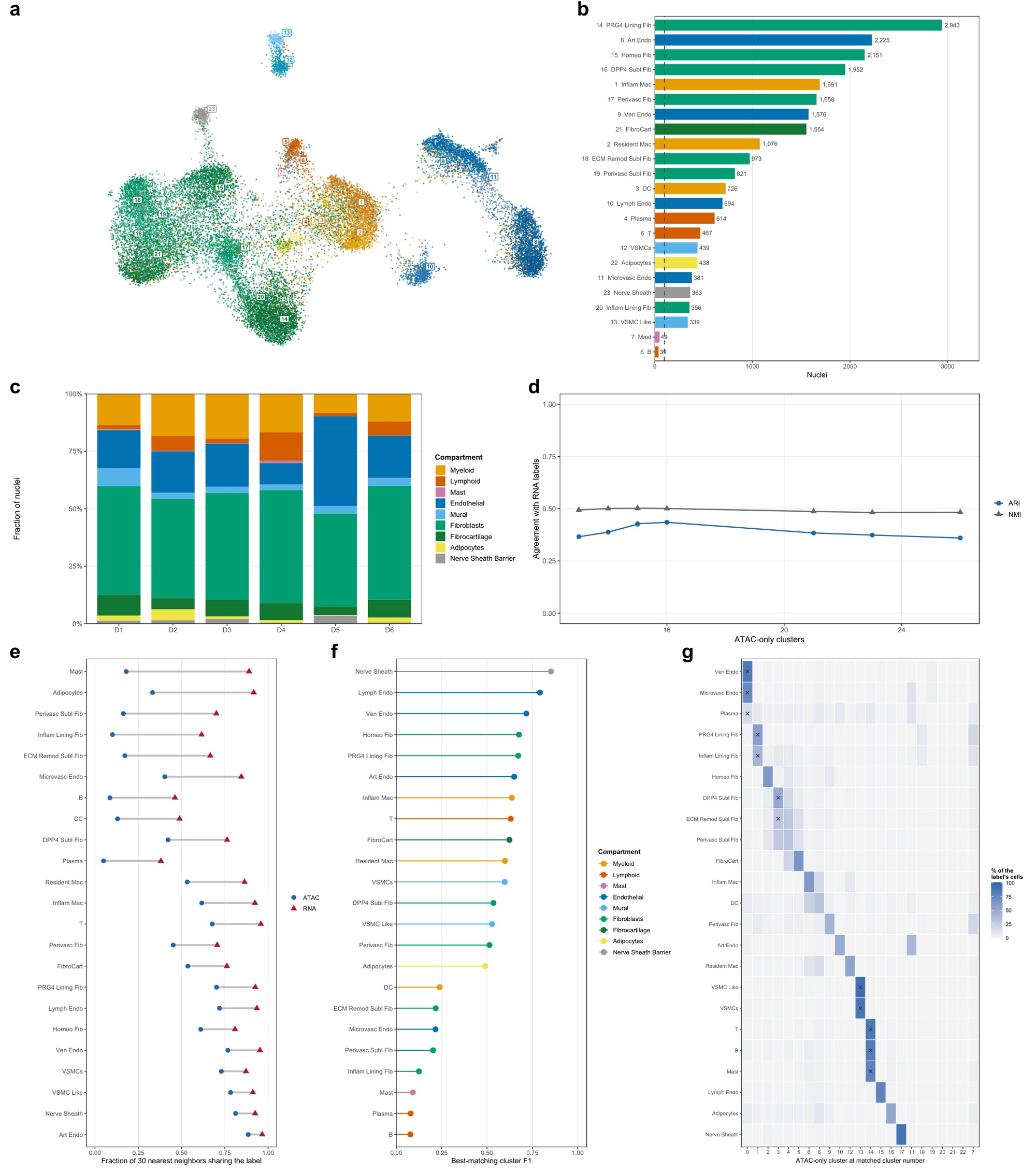

**Supplementary Fig. 8 | The synovial chromatin atlas, and what chromatin alone resolves.** **a** The chromatin embedding, 23,525 nuclei, 23 populations, numbered as in b. **b** Nuclei per population; dashed line is the 100-nuclei analyzable floor. **c** Compartment composition per donor. **d** Chromatin-only clustering across a resolution sweep, scored against the transcriptomic taxonomy; adjusted Rand index peaks at 0.435. **e** Neighborhood purity, the fraction of a nucleus's 30 nearest neighbors sharing its label: 0.56 in chromatin against 0.82 in RNA on the same nuclei. **f** Per-population recovery at matched cluster number. **g** At 23 clusters against 23 labels, 11 populations receive a cluster of their own; the other twelve populations share five clusters, two or three populations each, including both fibroblast pairs within a layer. Residual donor structure (Supplementary Fig. 9d) makes e an upper bound.

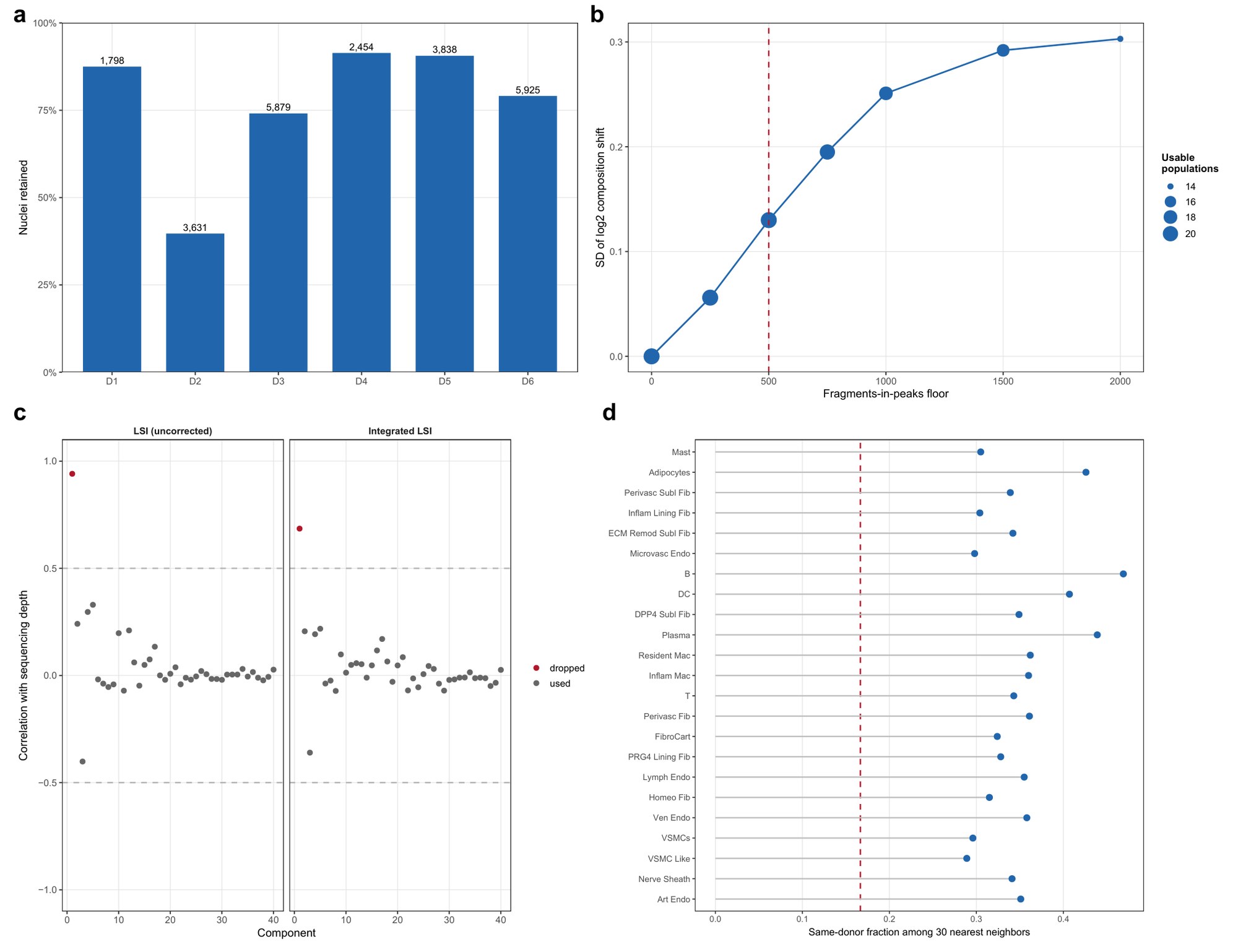

**Supplementary Fig. 9 | Depth threshold, batch correction and residual donor structure (synovium chromatin).** **a** Retention per donor after the fragment-depth filter, applied on the unified peak set. **b** Composition shift against candidate thresholds; the inherited floor of 1,000 dropped three populations below the analyzable bar, 500 keeps 21, and the dashed line marks the threshold used. **c** Correlation of each latent semantic indexing component with sequencing depth; only component 1 tracks it and is excluded downstream. **d** Same-donor nearest-neighbor fraction against expectation.

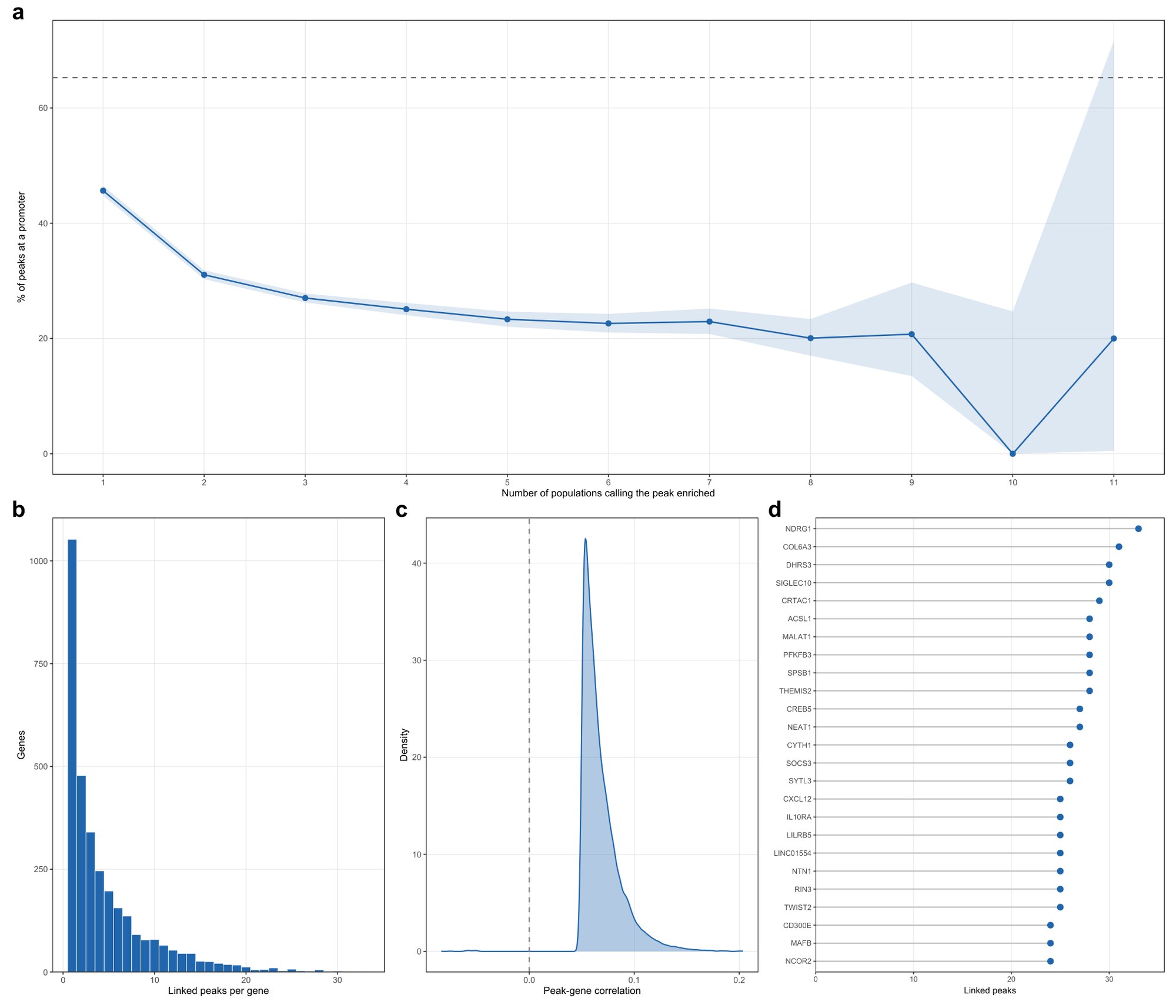

**Supplementary Fig. 10 | Sharing, genomic context and linked genes of the synovial marker peaks.** **a** The promoter trend of Fig. 3c at per-degree resolution with binomial confidence intervals: 45.7% at exclusive peaks to 22.3% at peaks shared by six or more, surviving adjustment for peak coverage (logistic odds ratio 0.812 per additional population, 0.732 adjusted). The dashed line is peaks enriched in no population, 65.3% promoter. **b** Peak-to-gene links from LinkPeaks within 500 kb in the same nuclei: 14,844 links across 3,226 genes. **c** Link strength distribution. **d** The most densely linked genes.

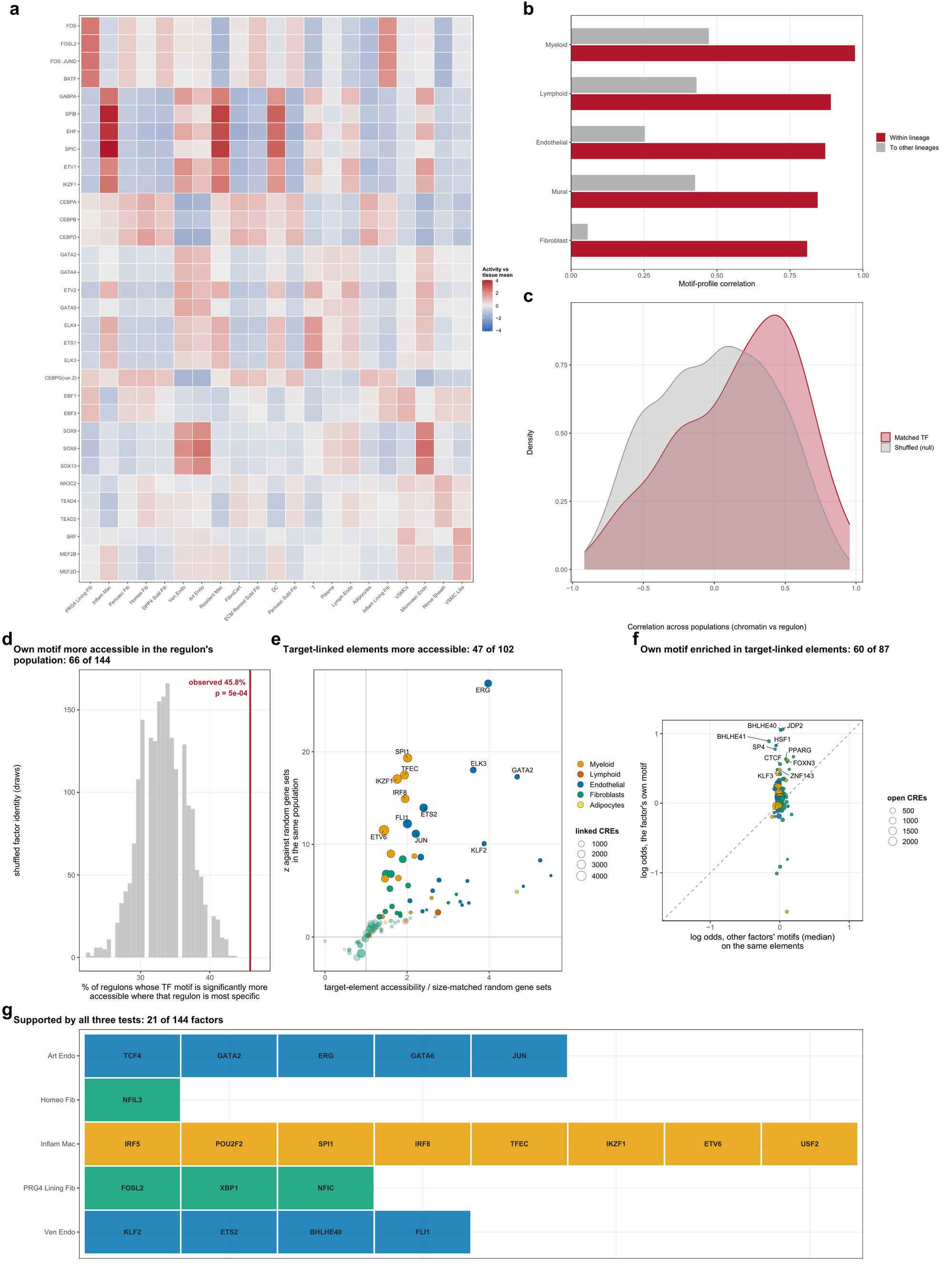

**Supplementary Fig. 11 | Synovial transcription-factor programs are shared within lineages and agree with the regulome in aggregate.** **a** chromVAR activity of the 32 leading motifs (of 633 JASPAR CORE motifs) across populations, an extended version of Fig. 3d; the leading motifs are a property of lineage, with CEBPD and CEBPA heading most fibroblast subtypes, SPIC and SPIB all three myeloid populations, and AP-1 the two lining subtypes. **b** Motif-profile correlation within and between lineages: r = 0.809–0.973 within; gray bars, mean correlation of each lineage to the other lineages (the mean over all population pairs, 0.369, is not drawn). **c** Matched against shuffled cross-modal correlation for the 144 factors carrying both a motif and a regulon, the null pairing each chromatin profile with a random regulon; three factors clear correction individually, SPI1, TFEC and ETV6 at padj = 0.048. **d–g** Whether each regulon is supported by open chromatin in the population where it is most specific. **d** The factor's own motif is more accessible there, 66 of 144 against 48 expected under shuffling. **e** The elements reaching its target genes are more accessible there than for size-matched random gene sets, 47 of 102 testable factors. **f** Those elements carry the factor's own motif above other factors', 60 of 87. **g** The 21 factors supported by all three tests, by the population in which each regulon is most specific.

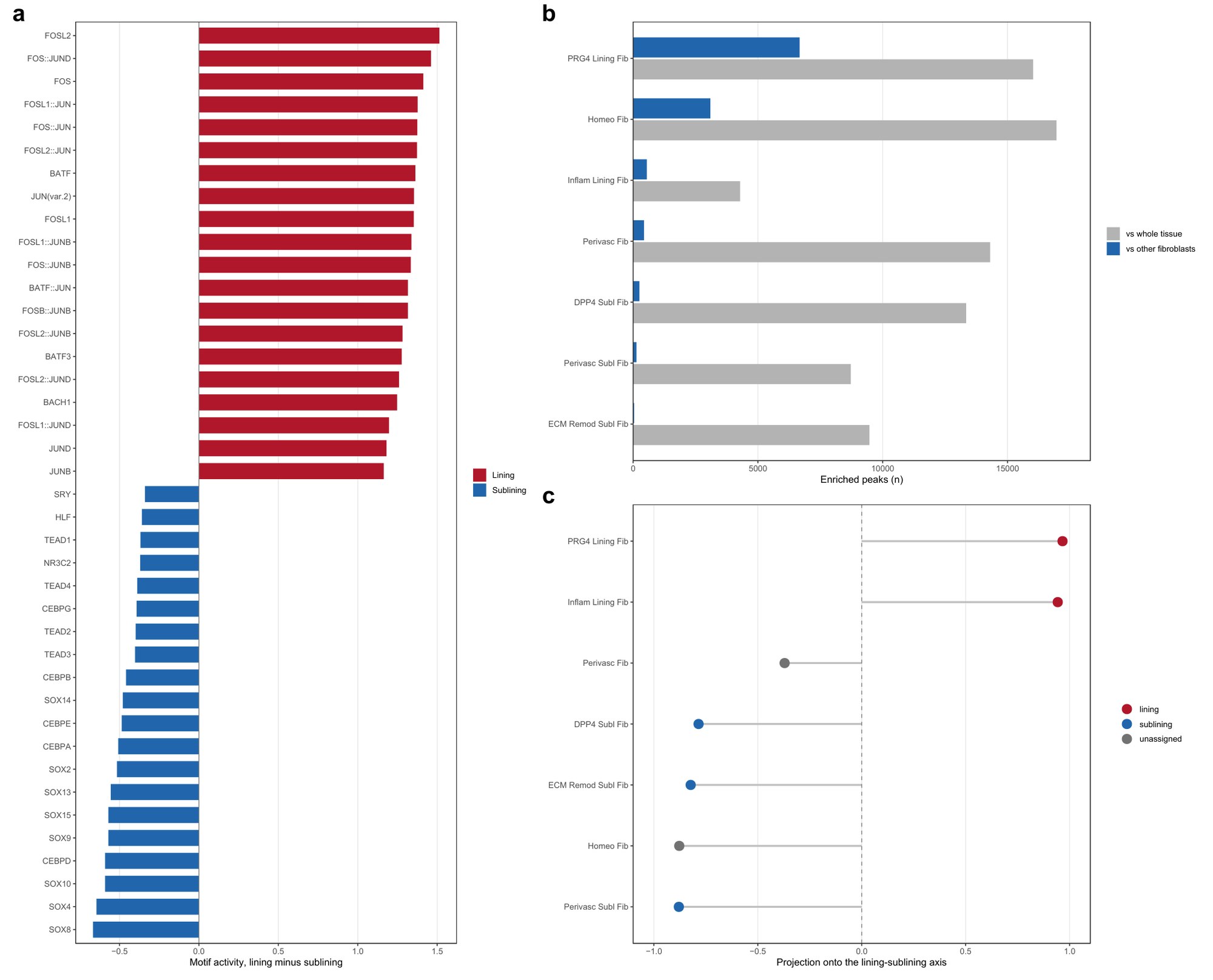

**Supplementary Fig. 12 | One regulatory boundary in the synovial fibroblast compartment, and a continuum below it.** Both models refitted on fibroblast columns only, so each population is contrasted against other fibroblasts. **a** Motifs separating the two layers: 421 of 633, twenty strongest each way; this is the reference version of Fig. 3f. **b** Enriched peaks retained when the denominator changes from the whole tissue to the compartment; PRG4 lining fibroblasts retain 42% and the other six subtypes 0.4–18% (homeostatic 18%, inflammatory lining 13%, perivascular 3% and the three sublining subtypes 0.4–2%). **c** A lining–sublining axis built from the five populations whose names declare a layer, with the other two projected onto it; both fall on the sublining side.

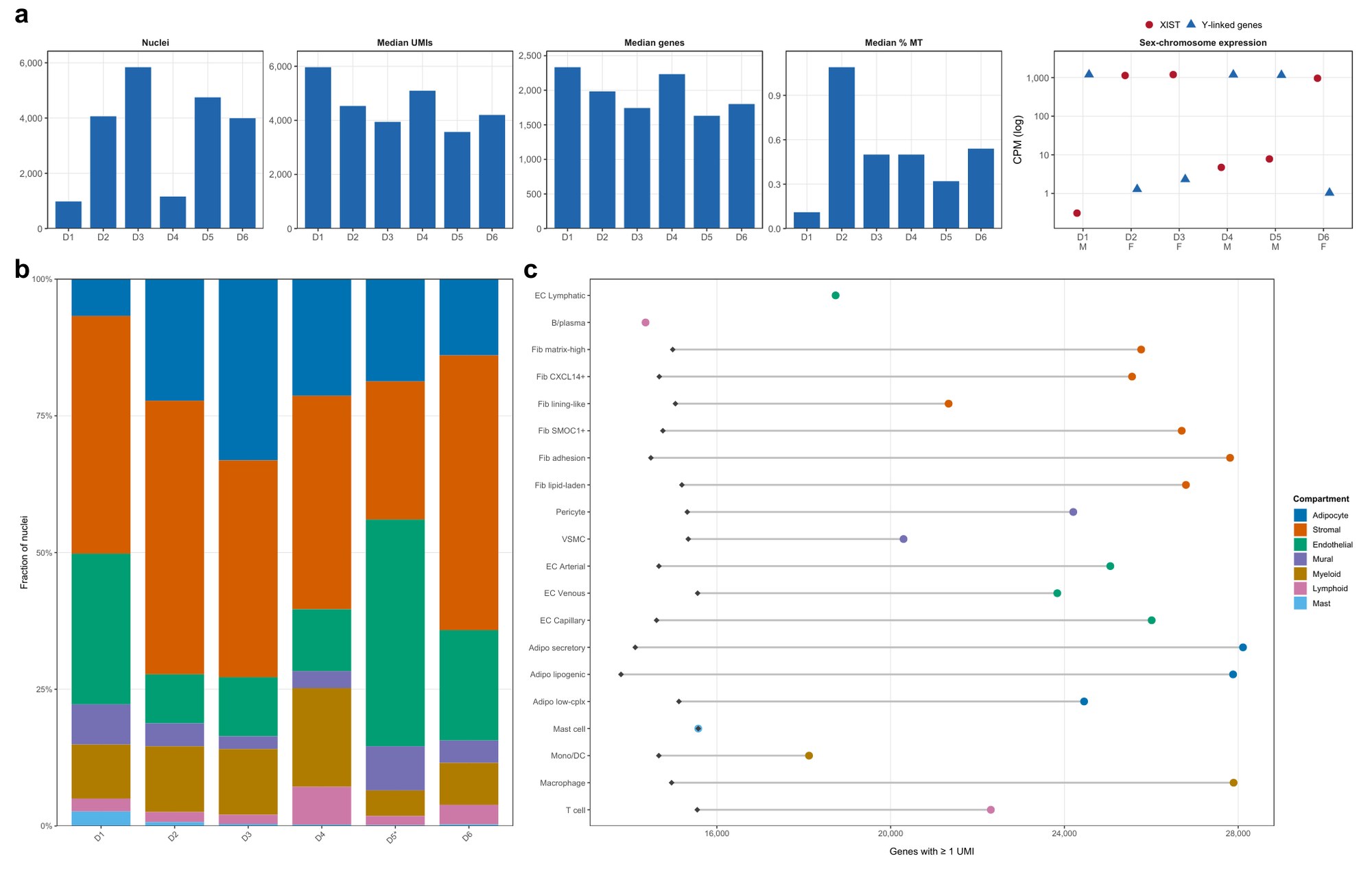

**Supplementary Fig. 13 | Cohort, sequencing quality, composition and gene detection (IPFP).** Panels as Supplementary Fig. 2, for 20,784 nuclei and 20 populations. **a** Per-donor sequencing quality, and sex-chromosome expression (XIST and the summed Y-linked genes, counts per million in the IPFP nuclei) against recorded sex; all six donors agree. **b** Compartment composition per donor; of the three largest compartments the stromal is the most consistent (25–50%) and the endothelial the least (9–41%), and the starred donor is the most endothelium-skewed. **c** Genes detected, as sequenced and depth-matched to the shallowest of the 18 abundance-filtered populations; lymphatic endothelium and B/plasma cells carry no matched point. Once depth is equalized the 18 populations detect within 13% of each other (13,794–15,572 genes). Population labels in Supplementary Figs. 13–24 are abbreviated (for example, Fib SMOC1+, SMOC1+/GHR+ fibroblast; Fib CXCL14+, CXCL14+/IGFBP7+ fibroblast; Adipo low-cplx, low-complexity adipocyte).

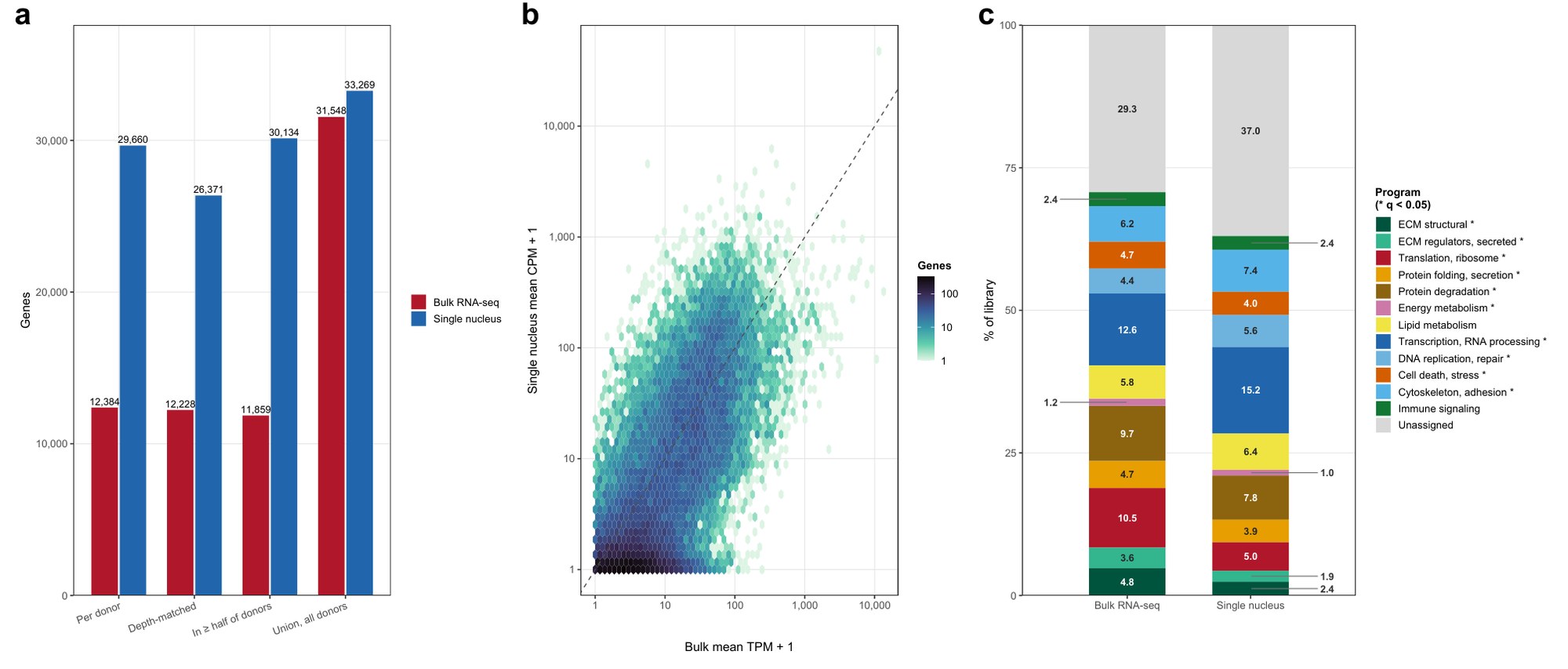

**Supplementary Fig. 14 | The healthy IPFP reference against bulk RNA-seq of the same tissue.** Panels as Supplementary Fig. 3. **a** Genes detected per platform. **b** Concordance, Spearman ρ = 0.66 over 29,759 shared genes. **c** The twelve programs of Fig. 4c on each platform; 10 of 12, marked with asterisks, differ at q < 0.05. All six donors have a bulk library from the same knee, five in the quality-passed set, and same-knee pairs agree no better than mismatched ones (mean ρ = 0.580 against 0.572).

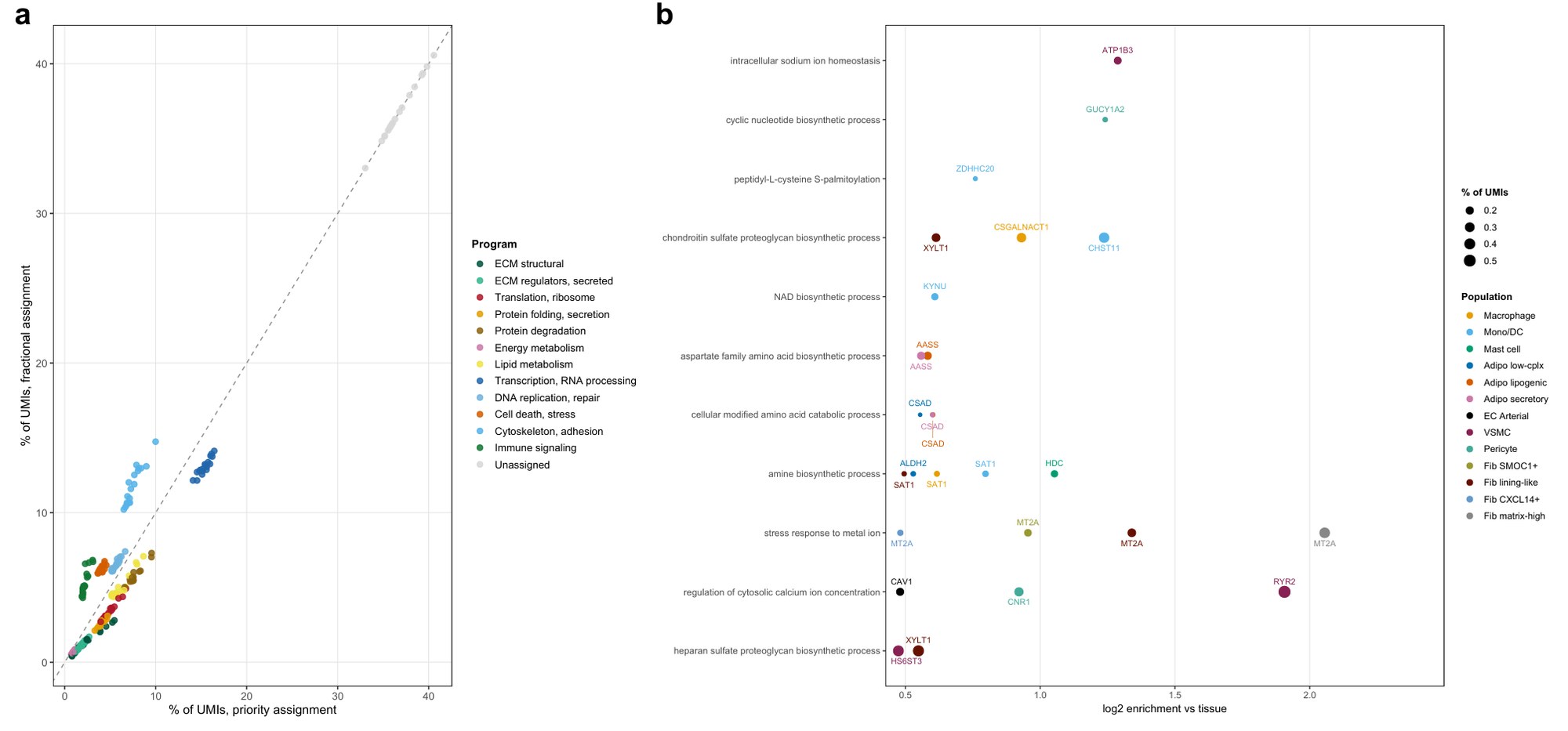

**Supplementary Fig. 15 | Robustness and coverage of the twelve-program transcriptional partition (IPFP).** Panels as Supplementary Fig. 4. **a** Fractional against priority assignment, r = 0.977. **b** The unbiased gene ontology scan; the strongest hit is a metal-ion stress response in the matrix-high fibroblast (log2 enrichment 2.06, top gene MT2A), followed by regulation of cytosolic calcium in vascular smooth muscle (1.91, RYR2).

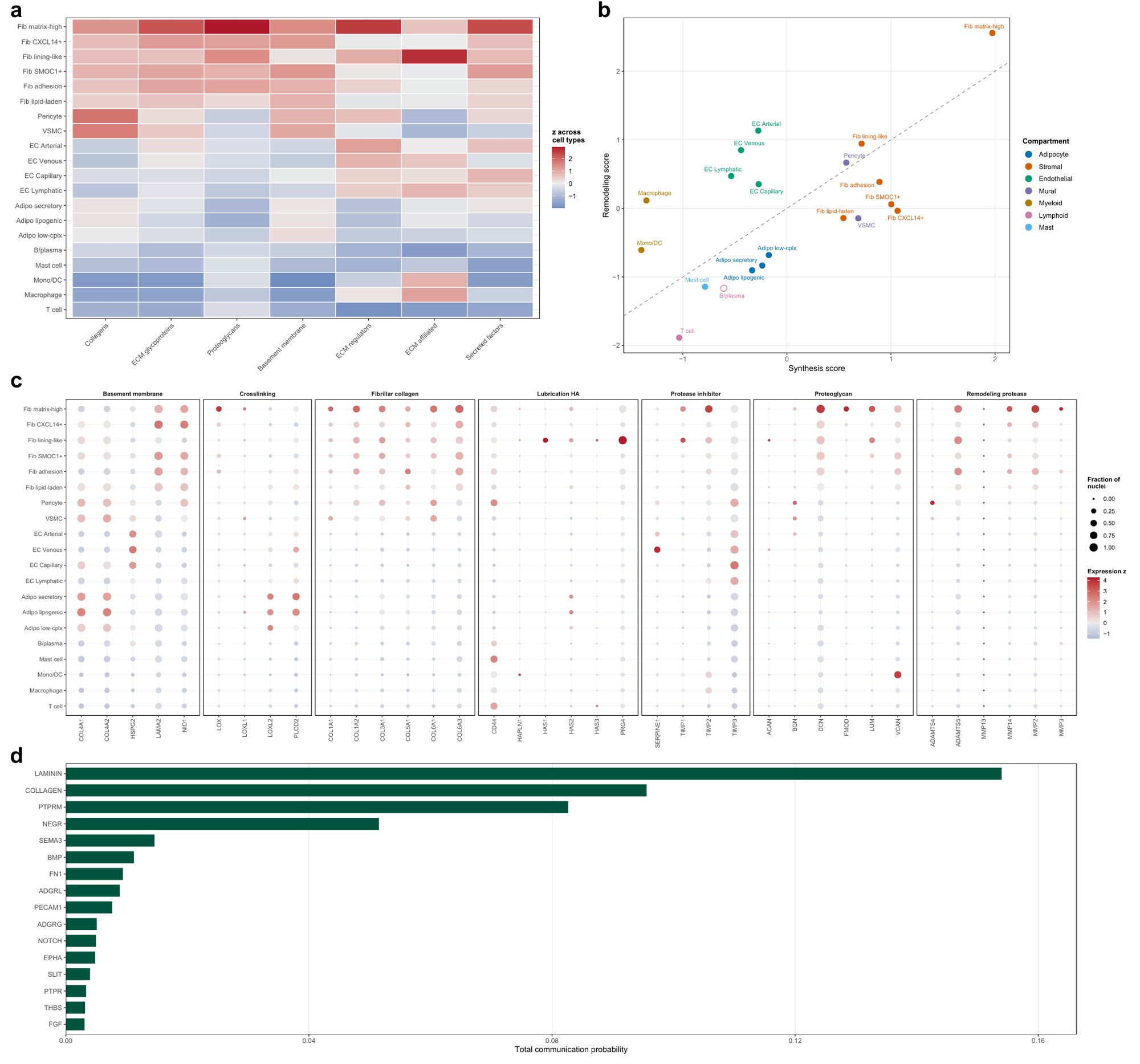

**Supplementary Fig. 16 | Matrisome composition, the synthesis–remodeling balance and predicted matrix signaling (IPFP).** Panels as Supplementary Fig. 5. The marker panel in c is the one curated for synovium, reused here because those groups are generic matrix biology; it has not been re-reviewed for the IPFP.

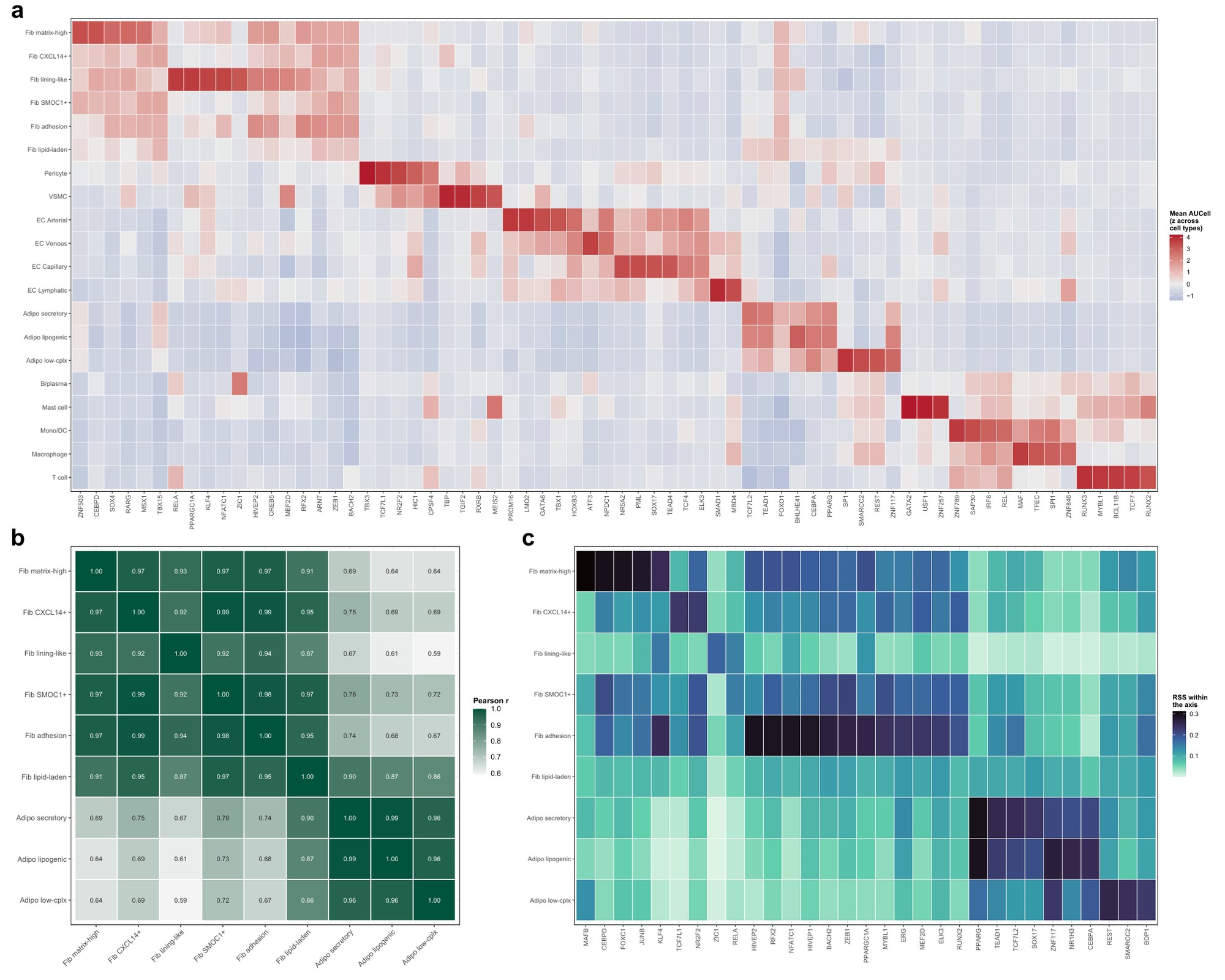

**Supplementary Fig. 17 | Regulon activity of the IPFP populations.** pySCENIC on healthy IPFP nuclei only, 225 regulons, without our labels. **a** The five most specific regulons per population; PPARG and CEBPA are recovered in adipocytes, SPI1 and IRF8 in macrophages, BCL11B and TCF7 in T cells, GATA2 in mast cells. **b** Similarity of regulon activity across the nine states of the adipogenic axis. The two most similar populations in the whole tissue are the two adipocyte subtypes and the SMOC1+/GHR+ and CXCL14+/IGFBP7+ fibroblasts, both at r = 0.993, and all fifteen fibroblast pairs correlate at 0.87 or above. **c** Specificity recomputed within the axis. No subtype separates from its neighbors by regulon activity: mean silhouette widths range from −0.17 to 0.13, so raising the resolution of the mesenchyme did not give its subtypes separate regulatory programs. Of the 225 regulons, 39 are most specific to the adhesion subtype and 22 to the matrix-high one, against 3, 1 and 0 for the three subtypes drawn from the other stromal label.

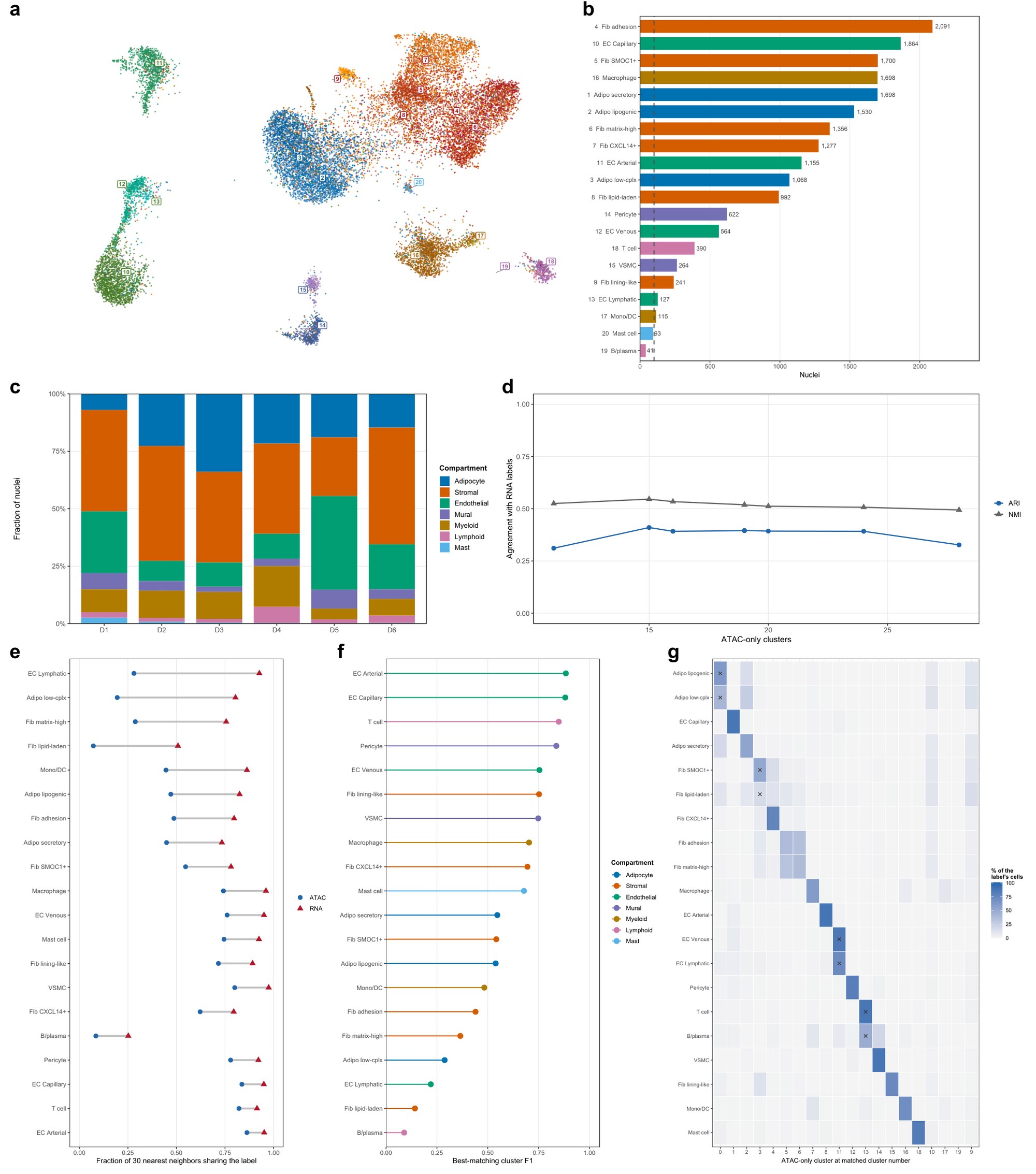

**Supplementary Fig. 18 | The IPFP chromatin atlas, and what chromatin alone resolves.** Panels as Supplementary Fig. 8, for 18,886 nuclei and 20 populations. b Dashed line, 100 nuclei; 18 of the 20 populations lie above it and 14 meet the testing threshold (at least 25 nuclei in each of at least four donors). **d** Adjusted Rand index peaks at 0.41 at 15 clusters. **e** Neighborhood purity 0.56 in chromatin against 0.83 in RNA, both nuclei-weighted. **g** At 20 clusters against 20 labels, 12 populations receive a cluster of their own; x marks the four clusters that are the largest destination of two populations (lipogenic with low-complexity adipocytes, SMOC1+/GHR+ with lipid-laden fibroblasts, venous with lymphatic endothelium, and T with B/plasma cells). The adhesion and matrix-high fibroblasts each lead one cluster but are split almost evenly between the same two. The direct test of whether the stromal subtypes separate within the axis is Supplementary Fig. 22, and of whether the original two labels did is Supplementary Fig. 23.

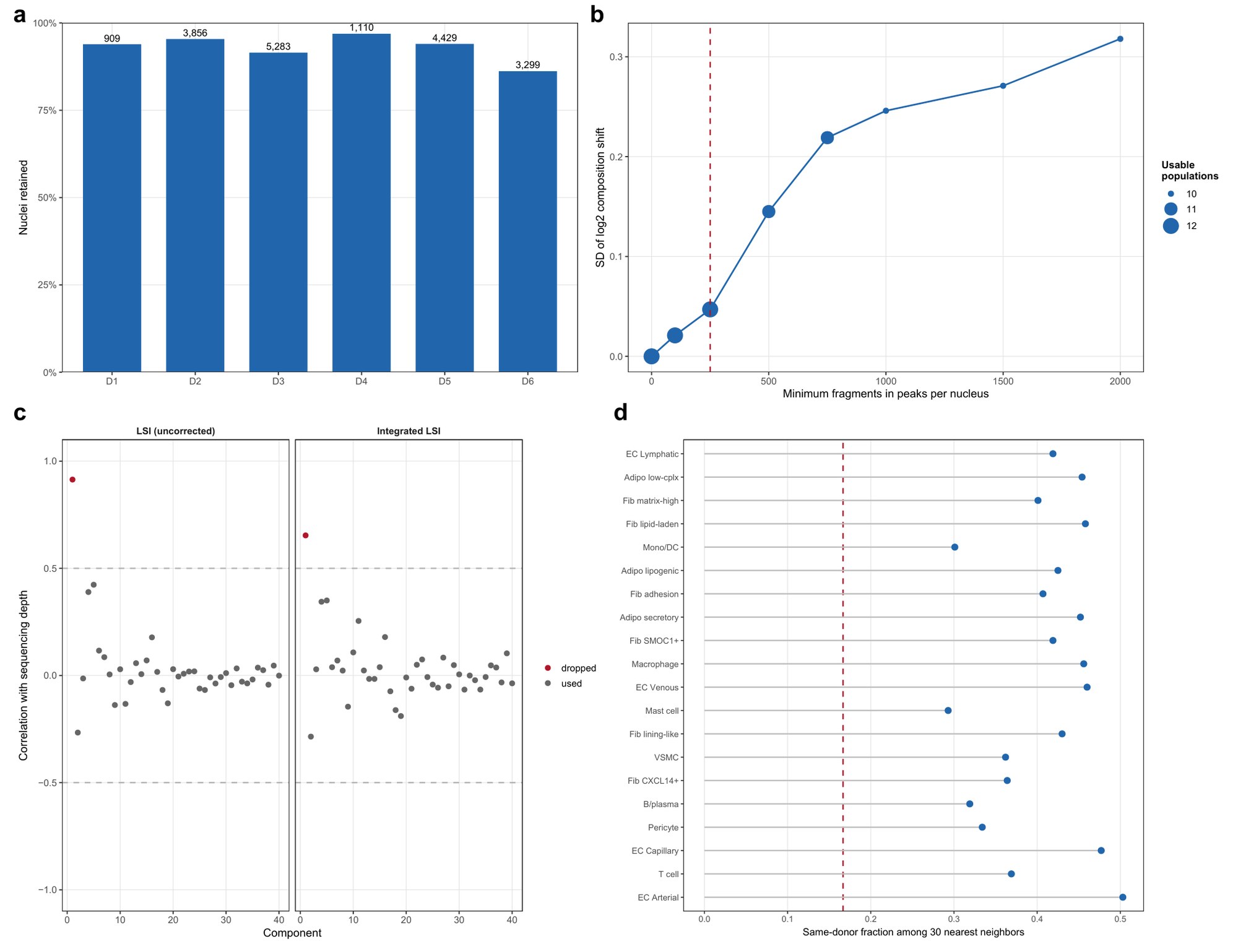

**Supplementary Fig. 19 | Depth threshold, batch correction and residual donor structure (IPFP chromatin).** Panels as Supplementary Fig. 9. **b** The floor was chosen by sweeping candidates against per-population retention: 250 retains 12 of the 14 original (pre-subclustering) populations as usable (at least 100 nuclei in at least four donors) at a composition-shift SD of 0.047, where 500 would retain 11 at 0.145. Retention ranges from 86.2% to 96.9% across donors. **c** Only latent component 1 tracks depth and is excluded.

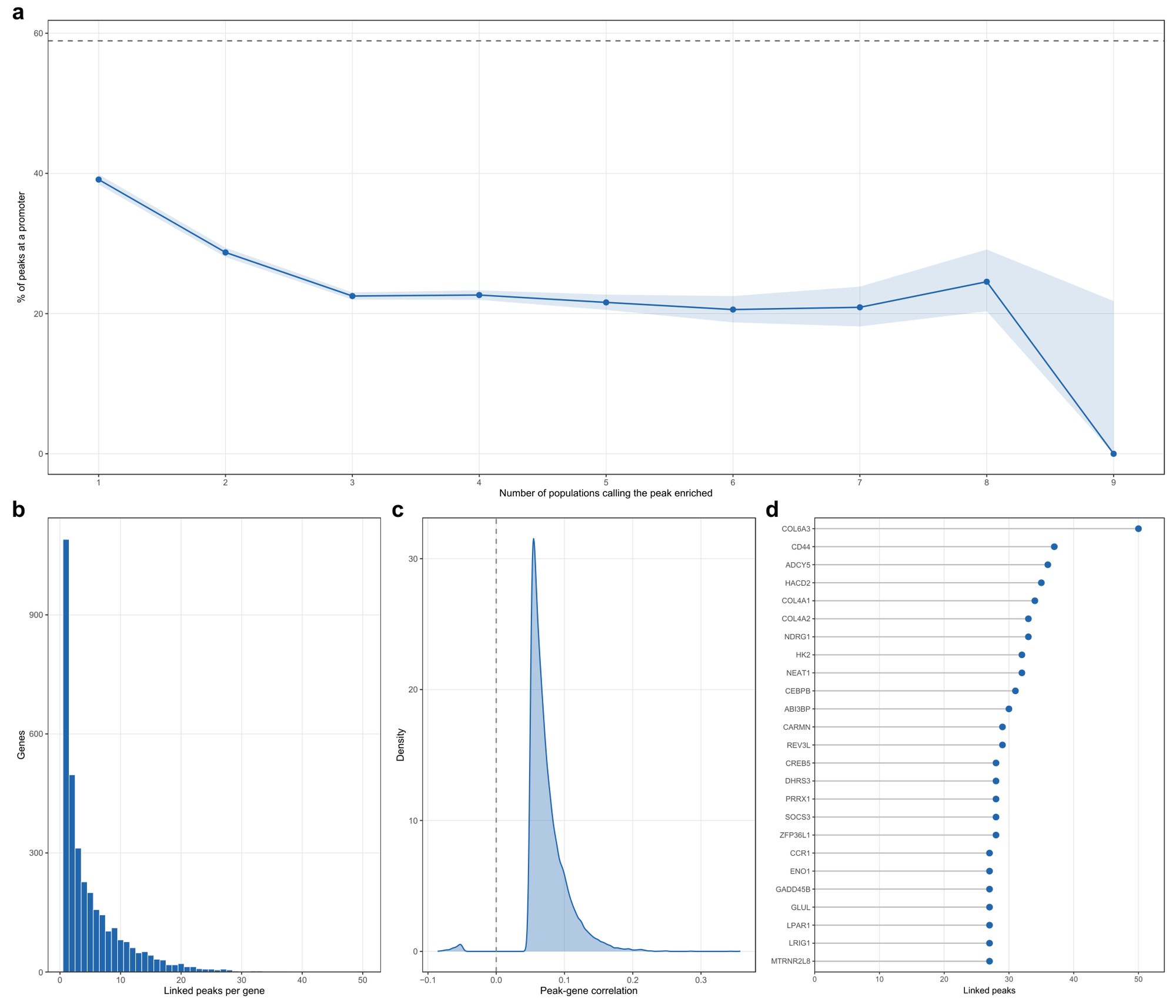

**Supplementary Fig. 20 | IPFP marker peaks, their genomic context, and the genes they reach.** Panels as Supplementary Fig. 10. **a** The promoter trend of Fig. 5c per degree with binomial confidence intervals: 39.1% at exclusive peaks falling to 21–23% at peaks enriched in three or more populations, the same decline as in synovium (the few peaks enriched in eight or nine populations give wide intervals). Dashed line: peaks enriched in no population, 58.9% promoter. **b** 17,141 peak-to-gene links across 3,397 genes. **c** Link strength distribution. **d** The most densely linked genes.

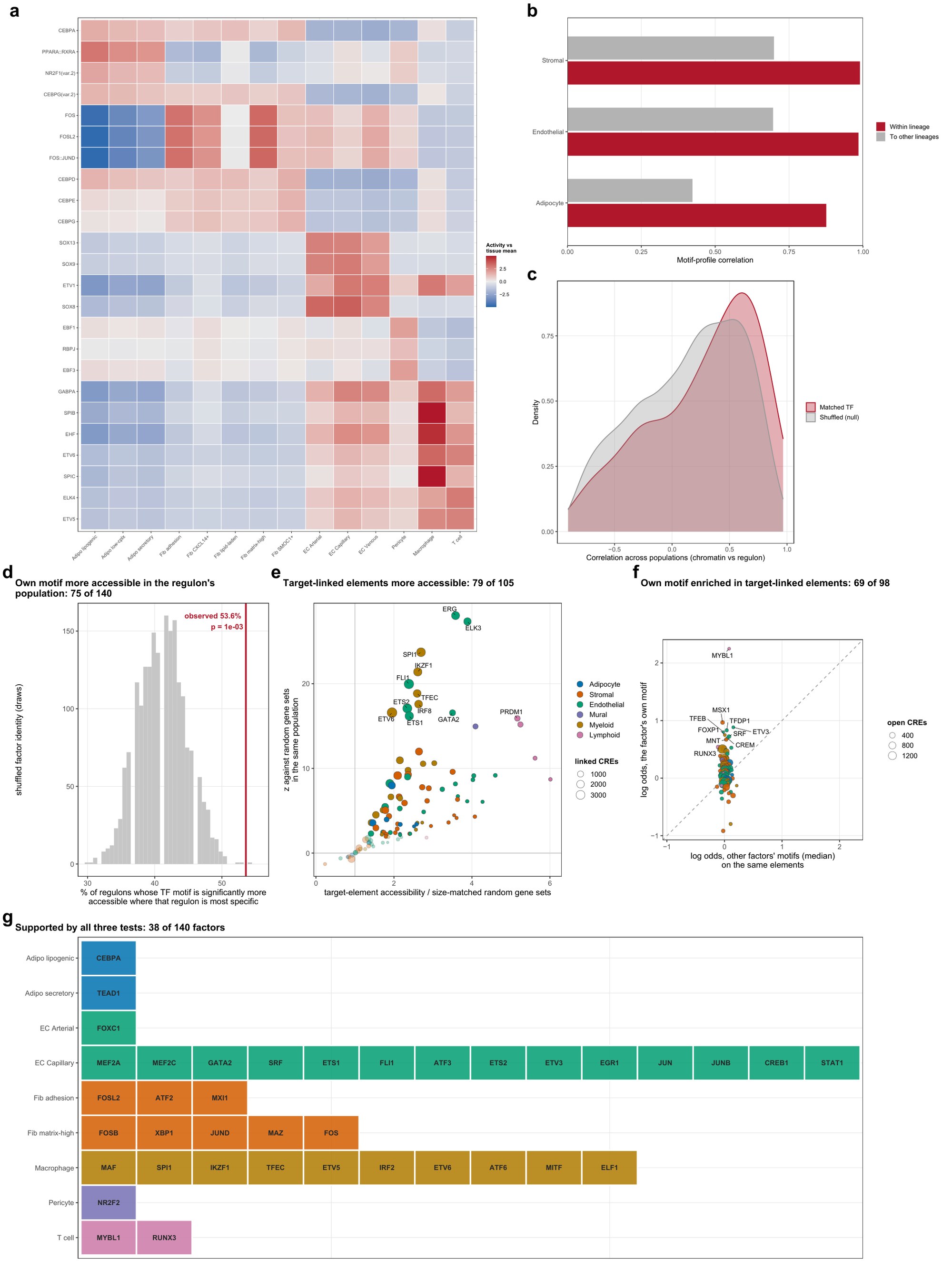

**Supplementary Fig. 21 | IPFP transcription-factor programs and cross-validation against the regulome.** Panels as Supplementary Fig. 11. **a** chromVAR activity of the 24 leading motifs (of 633) per population; all three adipocyte subtypes are headed by PPARA::RXRA followed by NR2F1 and CEBPA, while the fibroblast subtypes split into two signatures: the adhesion, CXCL14+/IGFBP7+ and matrix-high subtypes are led by AP-1, the SMOC1+/GHR+ and lipid-laden subtypes by the CEBP family. **b** Motif-profile correlation 0.99 across the fibroblast subtypes and 0.985 across the endothelial subtypes, against a mean of 0.70 over all population pairs (not drawn; gray bars, mean correlation of each lineage to the other lineages); the three adipocyte subtypes are the least coherent lineage at 0.876. **c** Cross-modal agreement for the 140 factors carrying both; median matched r = 0.387 against 0.212 for randomly paired factors. **d** The factor's own motif is more accessible there, 75 of 140 factors against 41.1% expected under a shuffled-identity null. **e** The elements reaching its target genes are more accessible there than for size-matched random gene sets, 79 of 105 testable factors. **f** Those elements carry the factor's own motif above other factors', 69 of 98. **g** The 38 factors supported by all three tests, by the population in which each regulon is most specific.

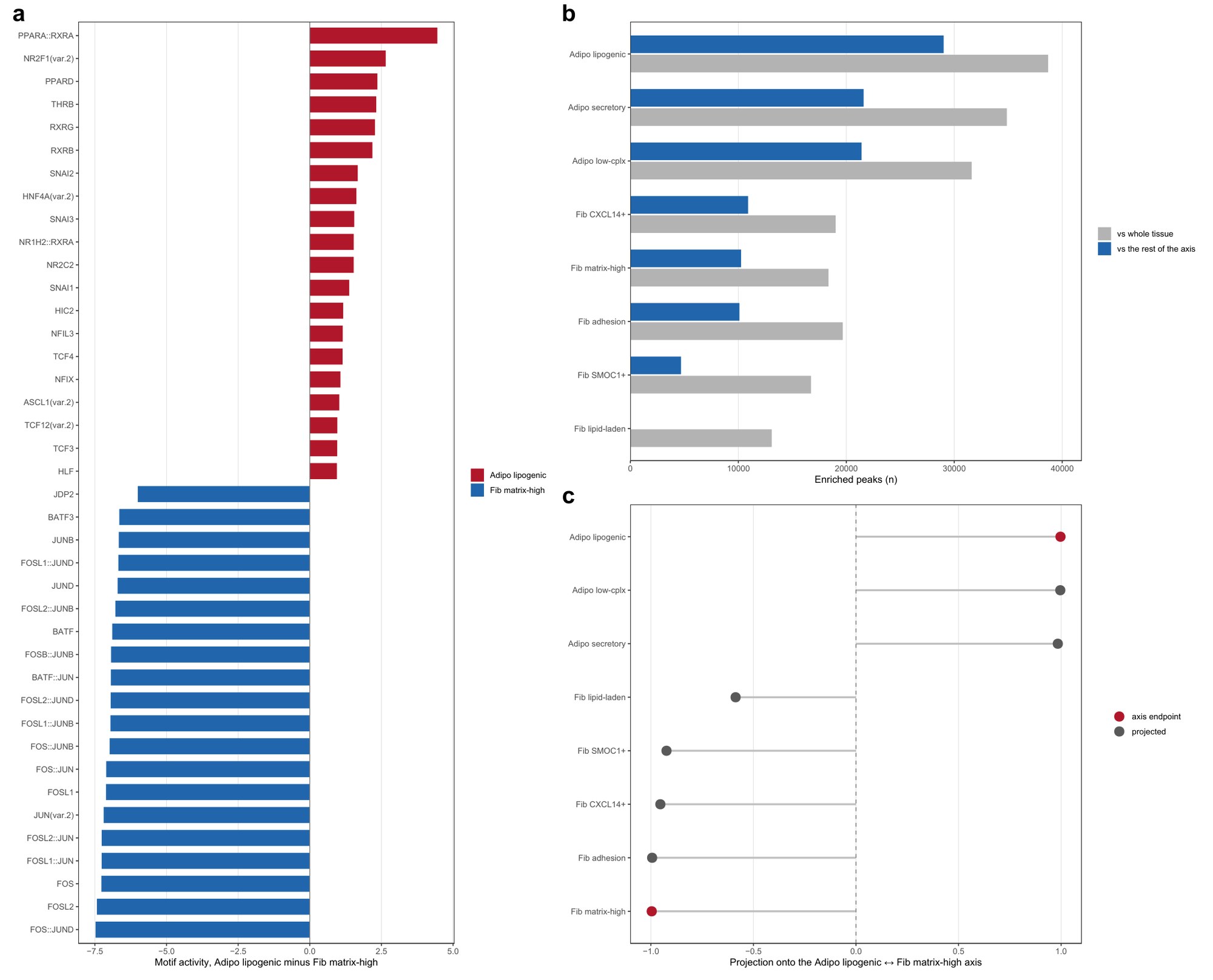

**Supplementary Fig. 22 | Inside the adipogenic axis: the adipocyte is a distinct state and the fibroblast subtypes occupy graded positions below it.** Both models refitted on axis columns only, over the three adipocyte and five testable fibroblast subtypes (the lining-like subtype has at least 25 nuclei in only three donors). **a** The contrast between the axis endpoints, lipogenic adipocyte against matrix-high fibroblast, twenty strongest motifs each way. Across all pairs (counts in Fig. 5f), every adipocyte-versus-fibroblast contrast separates 306 to 488 of 633 motifs; among the fibroblasts the number of differing motifs is graded, from 284 motifs between the lipid-laden and CXCL14+/IGFBP7+ subtypes down to none between the adhesion and matrix-high subtypes, and the two biologically named adipocyte subtypes differ by 17 (Fig. 5f). **b** Enriched peaks retained when the denominator changes from the whole tissue to the axis: 0.62–0.75 for the adipocyte subtypes and 0.51–0.57 for the adhesion, CXCL14+/IGFBP7+ and matrix-high subtypes, against 0.28 for SMOC1+/GHR+ and almost none for the lipid-laden subtype. **c** An axis built from the two endpoints, with the remaining subtypes projected onto it: matrix-high sits at the pole (−0.996), then adhesion (−0.994), CXCL14+/IGFBP7+ (−0.954) and SMOC1+/GHR+ (−0.924), with the lipid-laden subtype alone well away from the pole at −0.587 and the adipocytes at +0.98 to +1.00.

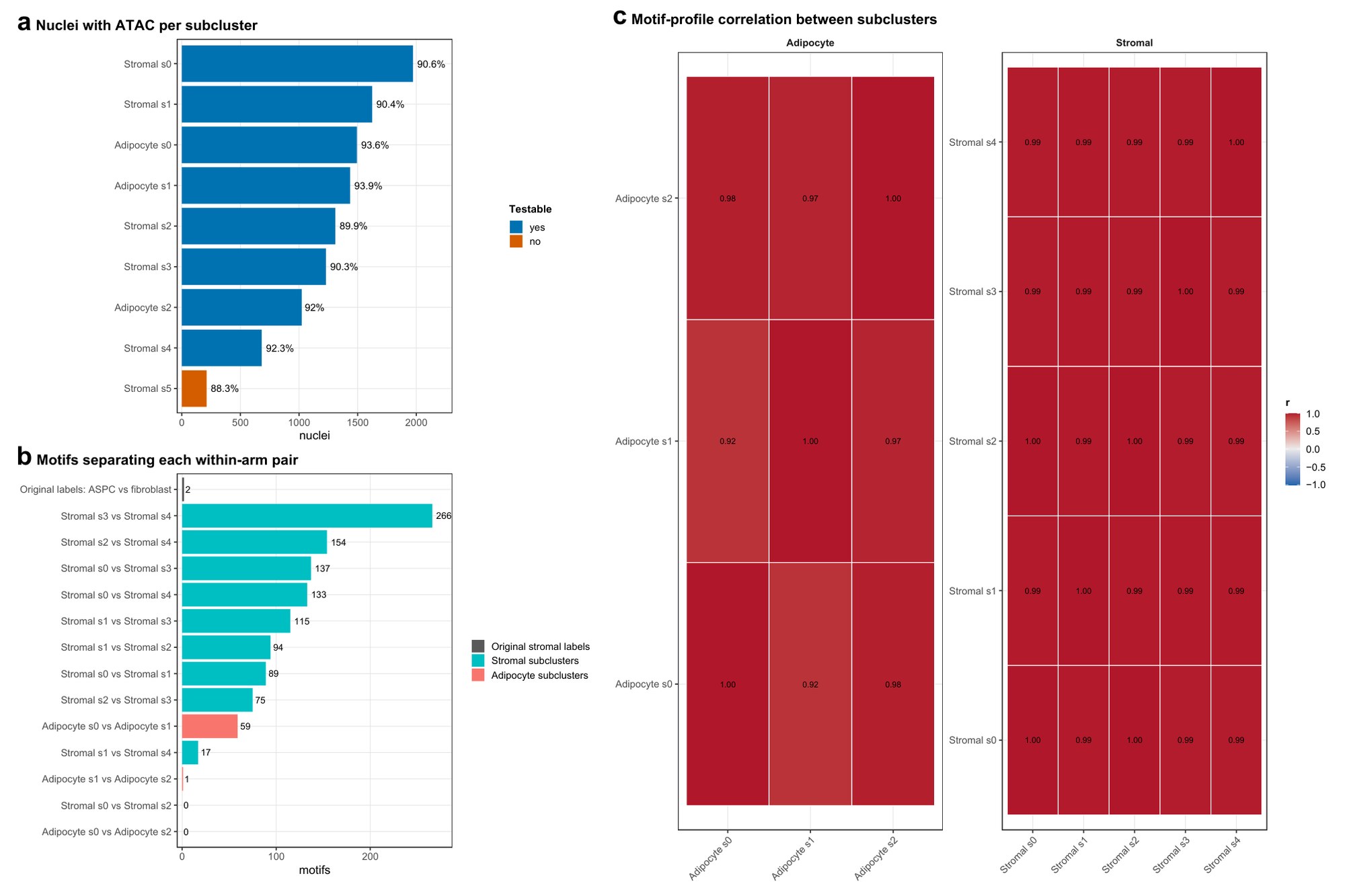

**Supplementary Fig. 23 | Chromatin test of the IPFP subdivision and of the original stromal labels.** **a** Nuclei with chromatin data per subcluster, with the percentage of the subcluster's nuclei that carry ATAC; subclusters below the testing threshold are marked. **b** Motifs separating each within-arm pair of subclusters (padj < 0.05, of 633 JASPAR motifs, same donor-blocked model as Fig. 5); the top bar is the parent-level control, the two stromal labels the reference originally carried re-fitted in the same model on the same nuclei. **c** Motif-profile correlation between subclusters within each arm. The main comparison is the parent-level control (top bar in b): re-fitted on the two stromal labels the reference originally carried, within the same nuclei, they separate at 2 of 633 motifs, while the subtypes drawn from those same nuclei separate at up to 266. The adipocyte arm shows that the test discriminates between subclusters: the two biologically named subtypes separate at 59 motifs, while the low-complexity population separates from them at 0 and 1. Subclusters: Adipocyte s0, secretory (CFD+/C1QTNF7+); s1, lipogenic (GPAM+/DGAT2+); s2, low-complexity. Stromal s0, adhesion (ITGA11+/SULF1+); s1, SMOC1+/GHR+; s2, matrix-high (COMP+/CILP+); s3, CXCL14+/IGFBP7+; s4, lipid-laden (CIDEA+/CD36+); s5, lining-like (CLIC5+/HAS1+).

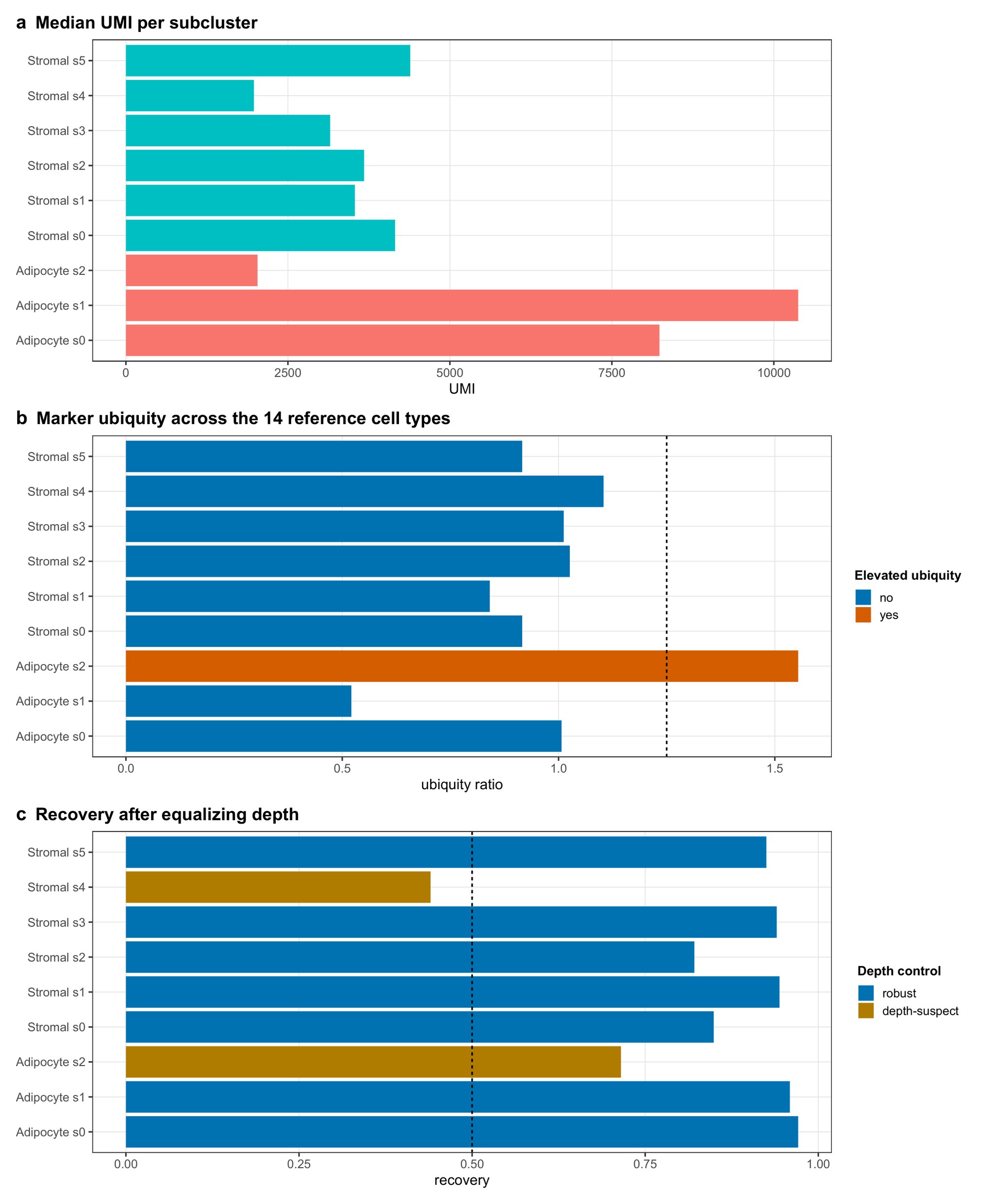

**Supplementary Fig. 24 | Depth control: which subclusters are defined by sequencing depth.** **a** Median UMI per subcluster. **b** Marker ubiquity: the median across-cell-type detection rate of each subcluster's markers relative to its arm, scored against the whole fourteen-population reference so the test is independent of this subclustering; dashed line, the elevation threshold. **c** Recovery after every nucleus in an arm is downsampled to a common UMI target and the arm re-clustered from scratch, as the adjusted Rand index between the full-depth and downsampled partitions; dashed line, 0.5. Bars are colored by the depth control: robust if recovery is at least 0.5 and marker ubiquity is not elevated (ratio ≤ 1.25, b), depth-suspect if either test fails. The low-complexity adipocyte subcluster (Adipocyte s2) fails as a cell state: median 2,031 UMI against 8,234 and 10,376, markers 1.55 times as ubiquitous as the arm's, and no chromatin identity (Supplementary Fig. 23). The lipid-laden stromal subcluster (Stromal s4) is depth-suspect, at median 1,976 UMI and 0.44 recovery. Subclusters as in Supplementary Fig. 23.

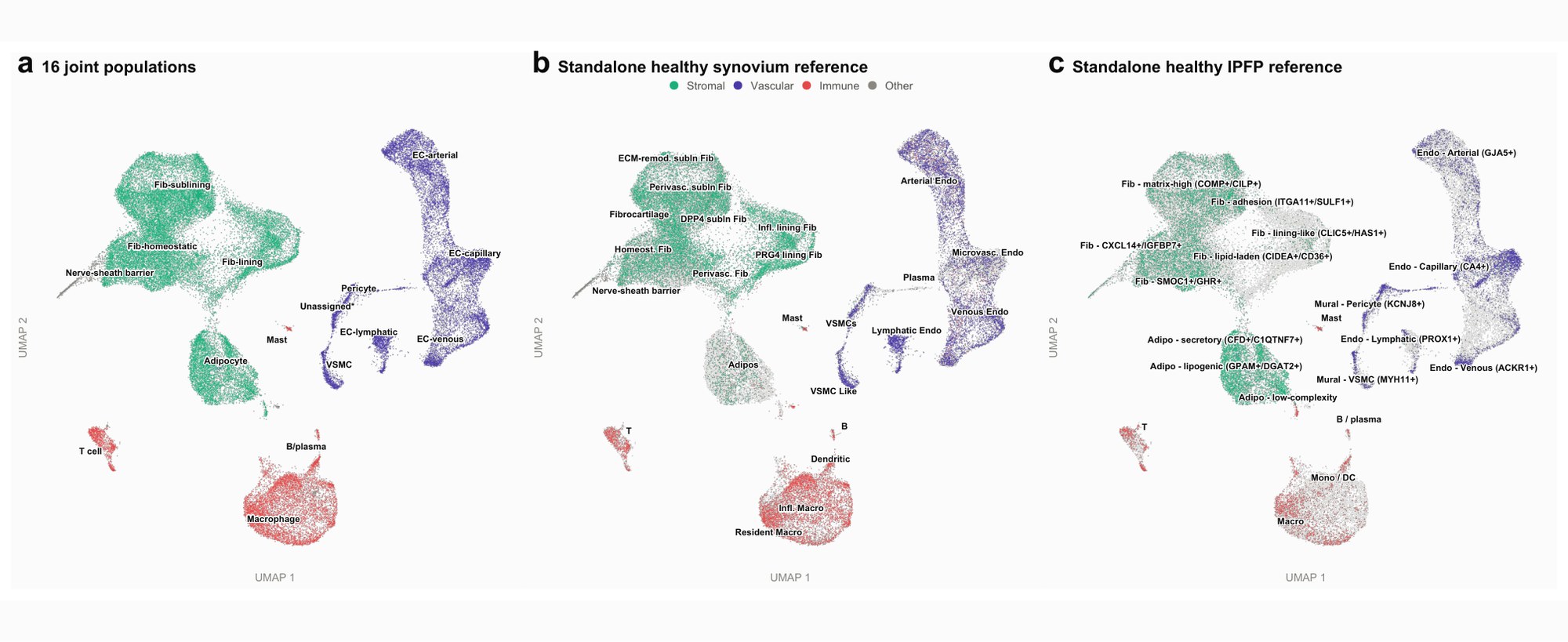

**Supplementary Fig. 25 | The 16 joint populations and the two standalone references, on one embedding.** **a** UMAP of all 54,716 nuclei by lineage family, labeled with the 16 populations annotated on the joint object (Methods), "Unassigned*" being the low-signal population retained for transparency — the same data as Fig. 6b at larger scale. **b** The same embedding with labels from the standalone healthy synovium reference (23 labels) transferred by exact barcode join; only the 33,932 synovial nuclei carry them and IPFP nuclei are gray. **c** As b for the standalone IPFP reference (20 labels) on the 20,784 IPFP nuclei. Transfer used exact barcode matching, with no label prediction, so all nuclei carry a reference label and the two references are tissue-disjoint by construction.

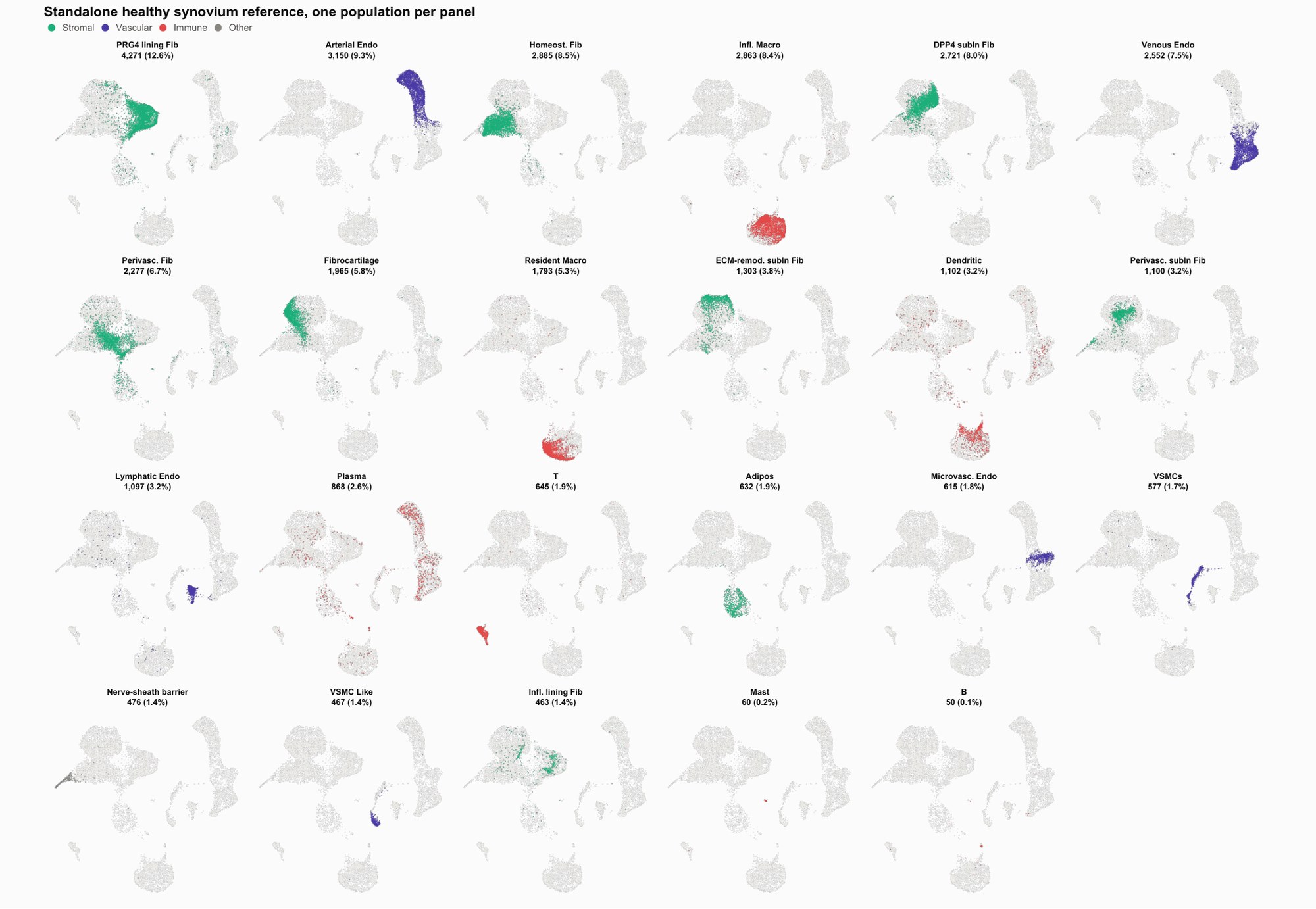

**Supplementary Fig. 26 | Each synovial reference population, located on the joint embedding.** One panel per label from the standalone synovium reference on the 33,932 synovial nuclei, ordered by size, with the number of nuclei and percentage of synovial nuclei in each title; gray is the synovial manifold for context (a 15,000-nucleus subsample, for rendering only) and color marks lineage family. Twenty-two of 23 populations place at least half their nuclei in one joint population and 19 place at least 70%. The exception is plasma cells (868 nuclei), which form no cluster: 36.2% fall in the Unassigned population and 33.8% on the arterial endothelial arm. The synovium reference's VSMCs place 82.0% of their nuclei in the joint pericyte population and its VSMC-like population 97.0% in joint vascular smooth muscle, so the mural compartment is split differently here.

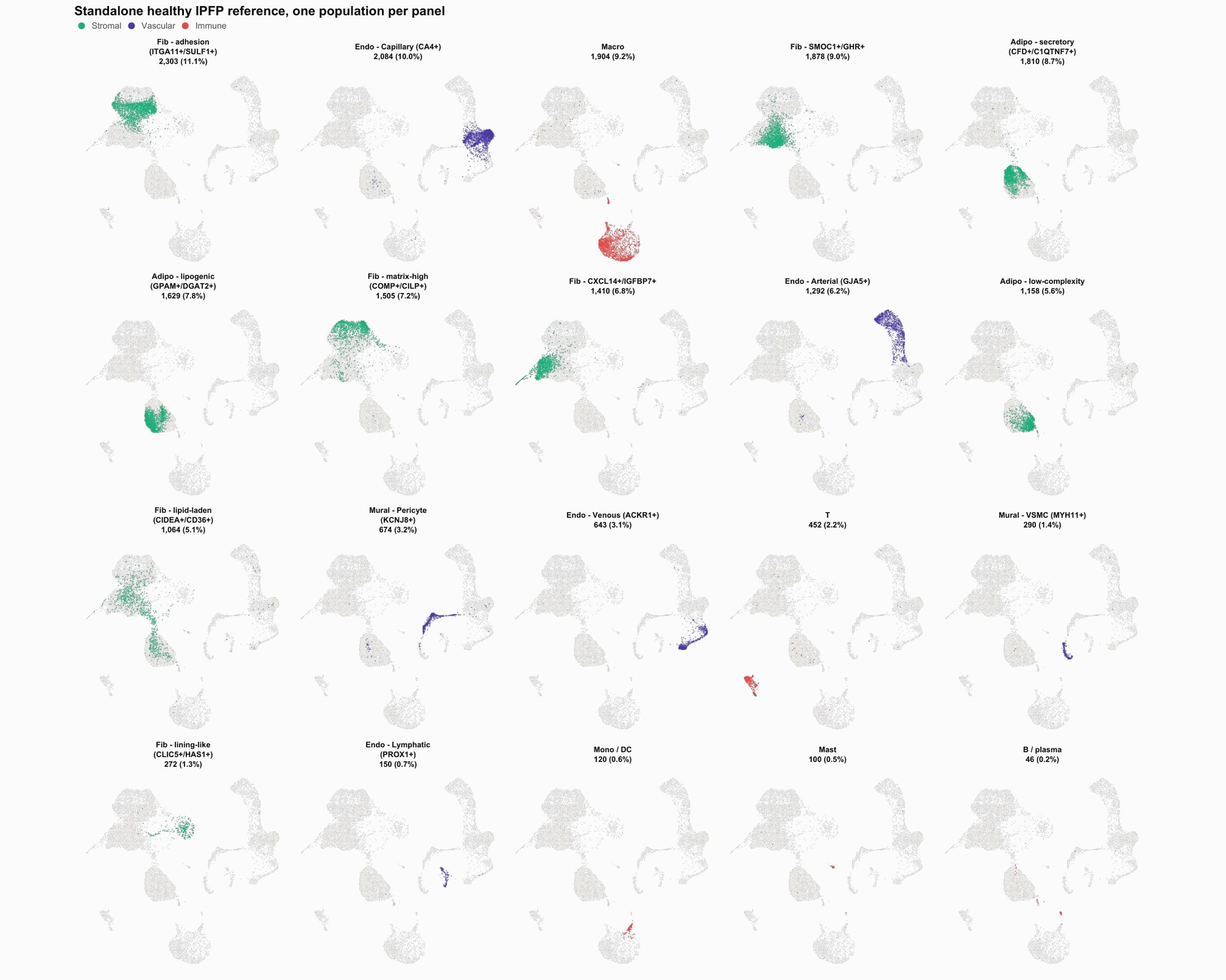

**Supplementary Fig. 27 | Each IPFP reference population, located on the joint embedding.** As Supplementary Fig. 26, for the 20 labels of the standalone IPFP reference on its 20,784 nuclei, one population per panel over four rows. The 118 IPFP nuclei the joint object places in its nerve-sheath barrier population sit mostly (99) among the CXCL14+/IGFBP7+ fibroblasts of this reference, which has no nerve-sheath category of its own (for the same reason only three lineage families appear here). The synovium reference does have one, and maps 95.4% of its 476 nerve-sheath nuclei onto the same joint population.

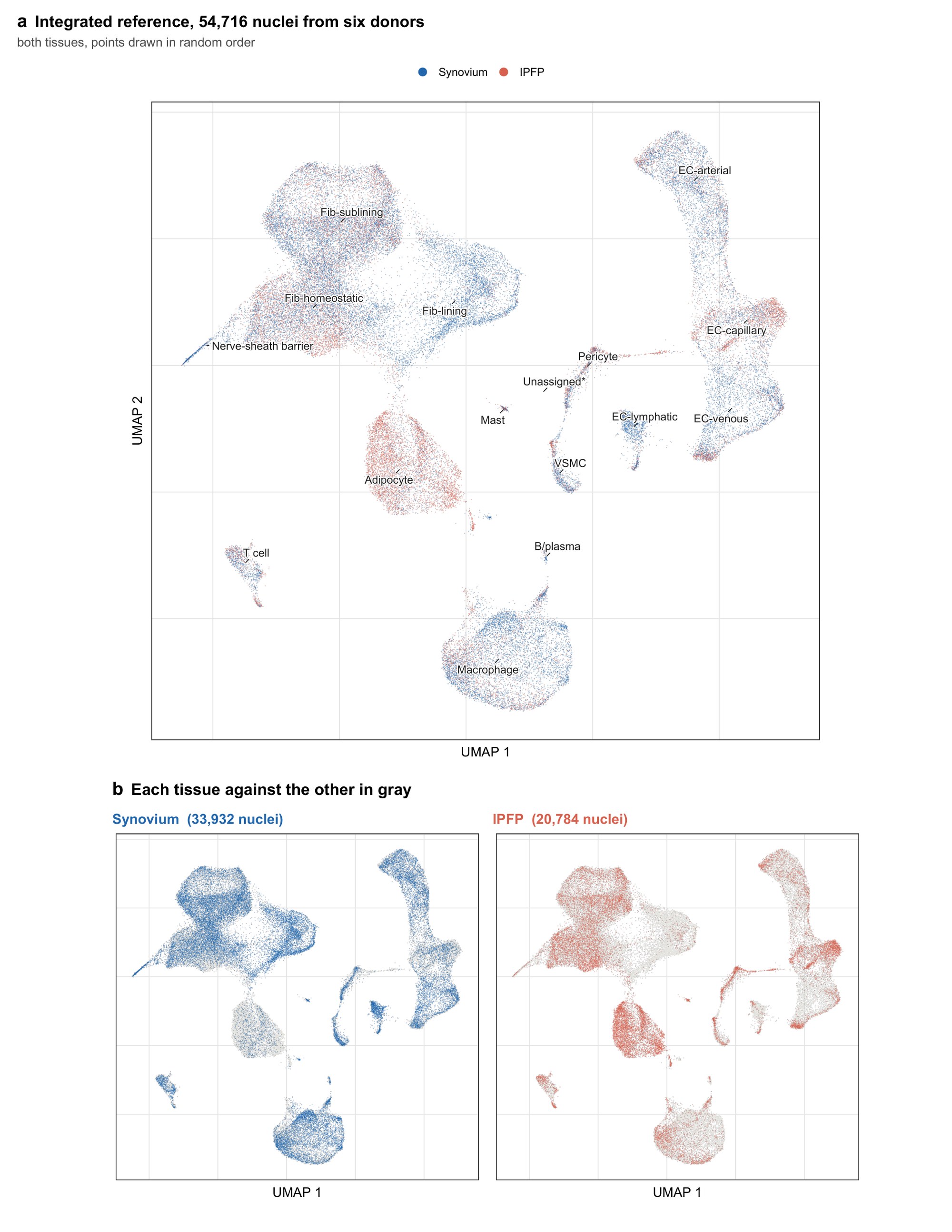

**Supplementary Fig. 28 | The integrated reference colored by tissue of origin.** The joint reference of 54,716 nuclei from six donors, each contributing both tissues, in the embedding the 16 populations were annotated in. **a** All nuclei, colored by dissected tissue of origin, with populations labeled at their median position. Points are drawn in random order, because on a single overlaid panel the tissue plotted last covers the other wherever they overlap. **b** The same embedding with each tissue shown separately against the other in gray, so that the position of each tissue is not affected by draw order. Synovium contributes 33,932 nuclei and the IPFP 20,784. Every population contains nuclei of both tissues; the extremes are the synovial lining fibroblast at 13.1% IPFP and the adipocyte at 84.5%. Nuclei per population per tissue are in the accompanying composition table.

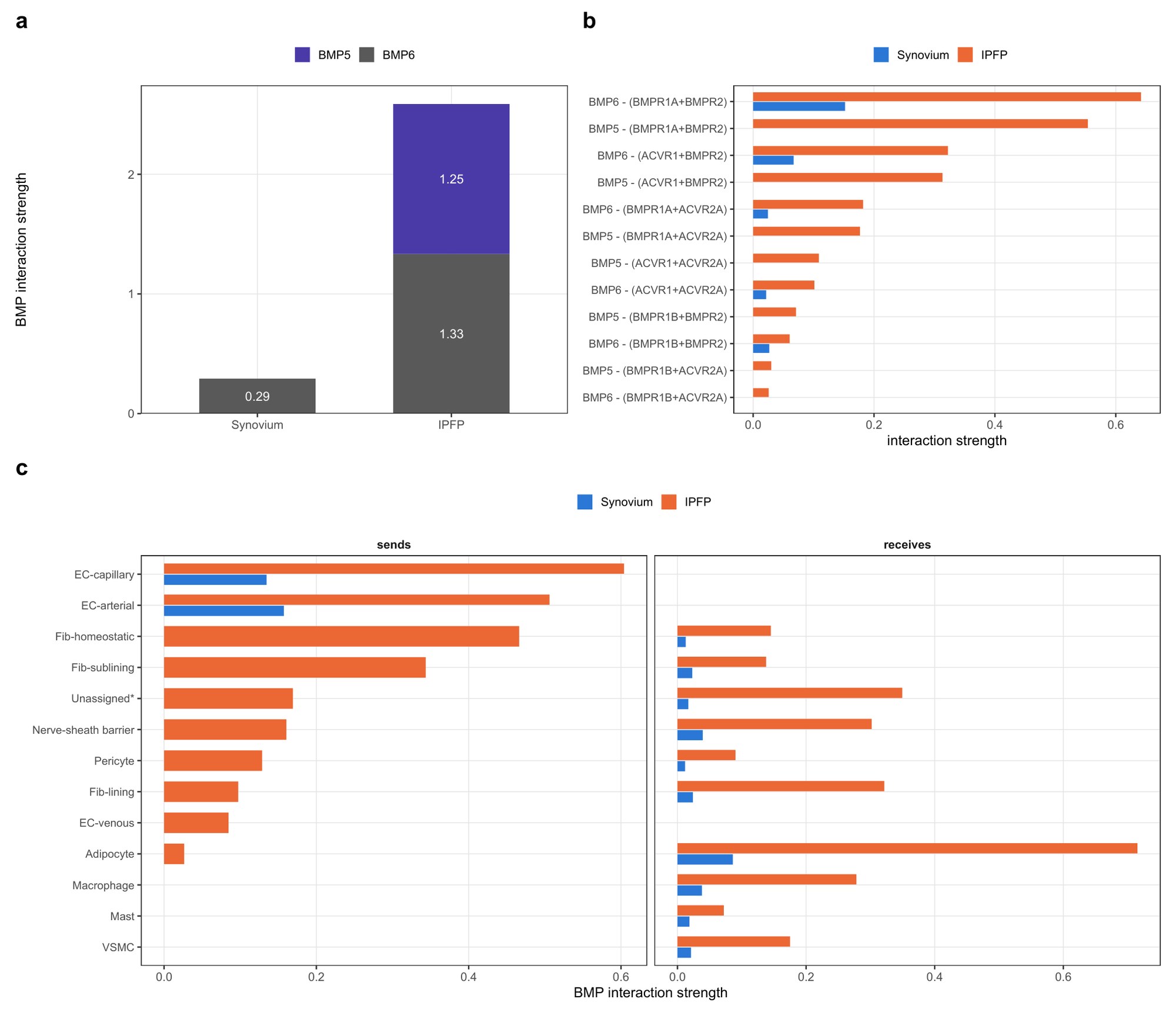

**Supplementary Fig. 29 | The IPFP's BMP signal is carried by two ligands, one of them not detected in synovium.** **a** Total BMP interaction strength in each tissue, split by ligand; numbers, summed strength per ligand. **b** Every BMP ligand–receptor pair with inferred signaling, by tissue. **c** BMP interaction strength sent (left) and received (right) by each population of the joint reference, by tissue; populations are sorted by IPFP sending strength, and those that only receive are at the bottom. "Unassigned*" is the low-signal population retained for transparency (Fig. 6). Interaction strength is the summed CellChat communication probability, inferred separately within each tissue without a population-size term; values compare within the BMP pathway only. Totals reproduce the BMP log2 fold change of Fig. 6f (+3.15). n = 33,932 synovium and 20,784 IPFP nuclei, six donors.

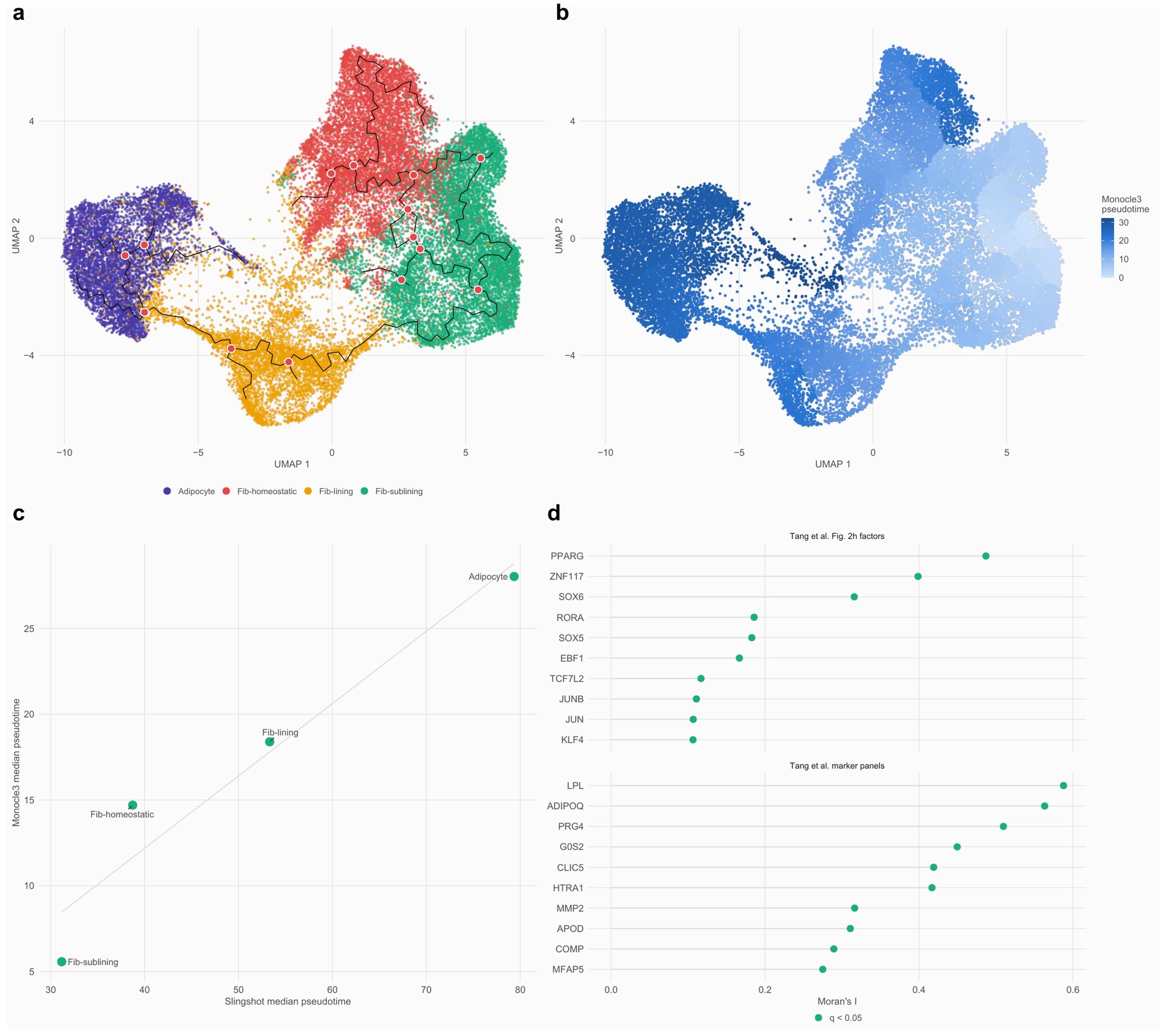

**Supplementary Fig. 30 | A second trajectory method, free to branch anywhere, on the same embedding (Monocle3).** **a** The principal graph over the nuclei with its branch nodes marked, on the same PCA and UMAP as the Slingshot fit. **b** Monocle3 pseudotime, rooted at the same population. **c** Median pseudotime per population under the two methods, against each other. **d** Variation across the graph (Moran's I), a trajectory-free test, for the published trajectory factors and marker-panel genes; the ten highest of each set are shown; read the effect size, since at 30,455 nuclei all 70 genes clear significance.

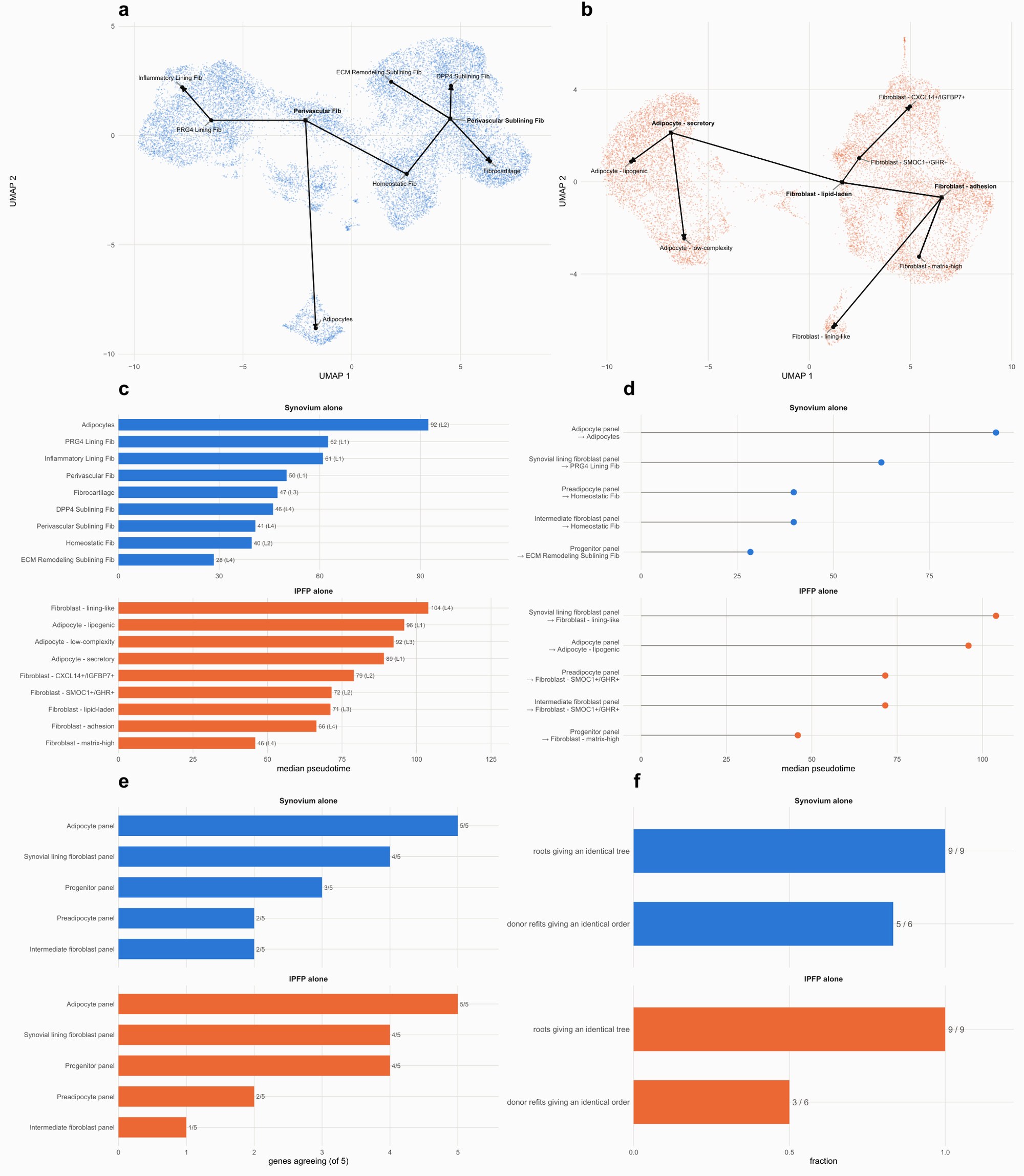

**Supplementary Fig. 31 | The trajectory refitted independently inside each tissue's own reference.** Each tissue in its own embedding from its own within-tissue integration, with its own annotation and its own root, and no correction between the tissues in either geometry. **a** Synovium alone, 17,617 nuclei over nine mesenchymal populations, with the fitted curves. **b** IPFP alone, 13,029 nuclei over nine mesenchymal populations. At the reference's earlier three-population resolution a spanning tree could only be a path, so this arm could test only the order of populations; at nine it tests the topology on the same footing as the synovium, and returns four lineages as the synovium does. The branch sits at neither tissue's DPP4+ population, which is a terminal end of both. **c** Median pseudotime per population, each read on the lineage carrying most of its nuclei. **d** Where the population nominated by each of the five published mesenchymal marker panels (Tang et al.^1^) falls along that tissue's own trajectory; in both tissues the preadipocyte and intermediate-fibroblast panels nominate the same population (homeostatic fibroblast in synovium, SMOC1+/GHR+ fibroblast in the IPFP). **e** Marker genes per panel agreeing on one population, at each tissue's native resolution. **f** Robustness: the synovial tree was identical from all nine roots, in five of six leave-one-donor-out refits, and without fibrocartilage. The IPFP tree was likewise identical from all nine roots, but reproduced in only three of six donor refits, and one of its three branch nodes is the lipid-laden fibroblast, 30.5% of whose nuclei (325 of 1,064) are projected doublets; the IPFP trajectory is therefore better supported than the placement of its branches.

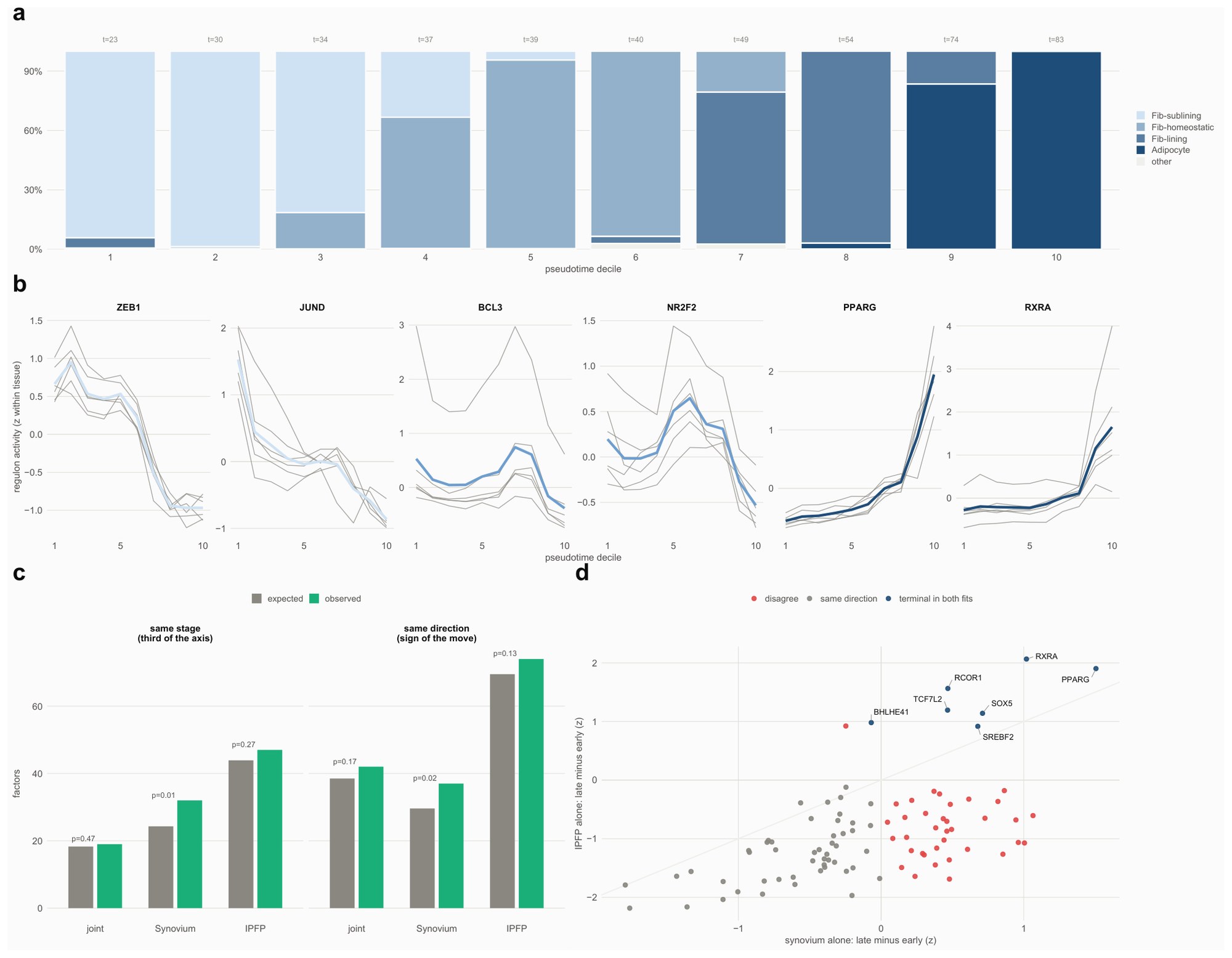

**Supplementary Fig. 32 | Donor support and reproducibility behind the stage regulators.** Regulon activity (AUCell) and motif accessibility (chromVAR over 633 motifs) attached to the trajectory nuclei by exact barcode join, standardized within tissue, and averaged per donor per pseudotime decile with the donor as replicate. The stage assignments are shown in Fig. 7e,f; this figure shows the donor-level data behind them. **a** What each decile is made of: population composition, with median pseudotime above each bar. **b** The largest mover of each stage per donor: one gray line per donor, donor mean in color. Six donors is the whole sample, so each donor is shown and no confidence band is drawn. **c** Factor-by-factor agreement between the two modalities against the expectation from 1,000 shuffles of which motif profile belongs to which factor, in each of the three trajectories; p is the permutation p-value. The count of factors whose regulon and whose own motif peak in the same third of the axis exceeds expectation only in the synovium's own trajectory (32 of 59 against 24.3, p = 0.01) and not in the joint trajectory (19 of 63 against 18.3, p = 0.47) or the IPFP's (47 of 107 against 43.9, p = 0.27); the count whose regulon and motif move in the same direction from the first to the last deciles (right half) likewise exceeds expectation only in the synovium's own trajectory (p = 0.02, against 0.17 joint and 0.13 IPFP), so cross-modal agreement along the axis holds at the level of factor families. **d** Each regulon's move from the first two deciles to the last two along the synovium's own trajectory against the same quantity in the IPFP's; labeled points are terminal in both fits.

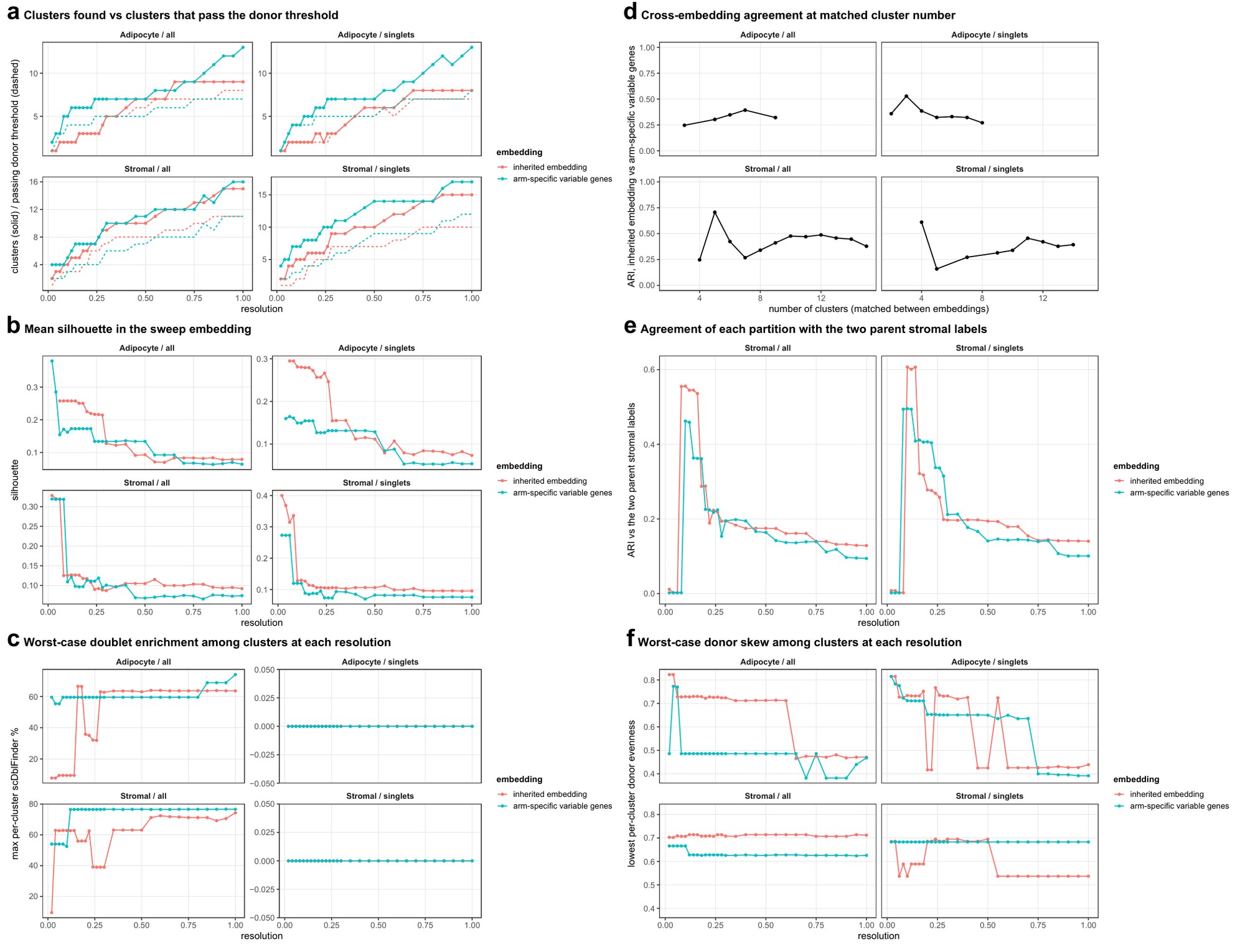

**Supplementary Fig. 33 | Choosing the resolution at which the IPFP mesenchyme is subdivided.** The adipocyte and pooled stromal nuclei were clustered across resolutions 0.02–1.0 in two feature sets: the embedding inherited from the whole-tissue integration and variable genes selected within the arm alone with Harmony correction on donor (arm-specific variable genes). Each panel shows both arms, for all nuclei and with predicted doublets removed (singlets). **a** Clusters found at each resolution (solid) and clusters passing the donor threshold of at least 25 nuclei in at least four of six donors (dashed). **b** Mean silhouette width, which does not identify a number: it is highest at the coarsest adipocyte split (two clusters) and flat near 0.10 (0.07–0.12) across stromal resolutions from six clusters to ten. **c** Worst-case doublet enrichment among the clusters at each resolution, as the largest per-cluster scDblFinder percentage; with predicted doublets removed it is zero throughout. **d** Agreement between the two feature sets at matched cluster number (adjusted Rand index), the criterion used: with predicted doublets removed, as they were for clustering, the adipocyte arm peaks at three subclusters (0.53, against 0.36 at two), and the stromal arm does not reach 0.50 at any resolution that places at least two donor-supported subclusters, so its subdivision is reported as descriptive. **e** Agreement of each partition with the two parent stromal labels. **f** Worst-case donor skew, as the lowest per-cluster donor evenness.

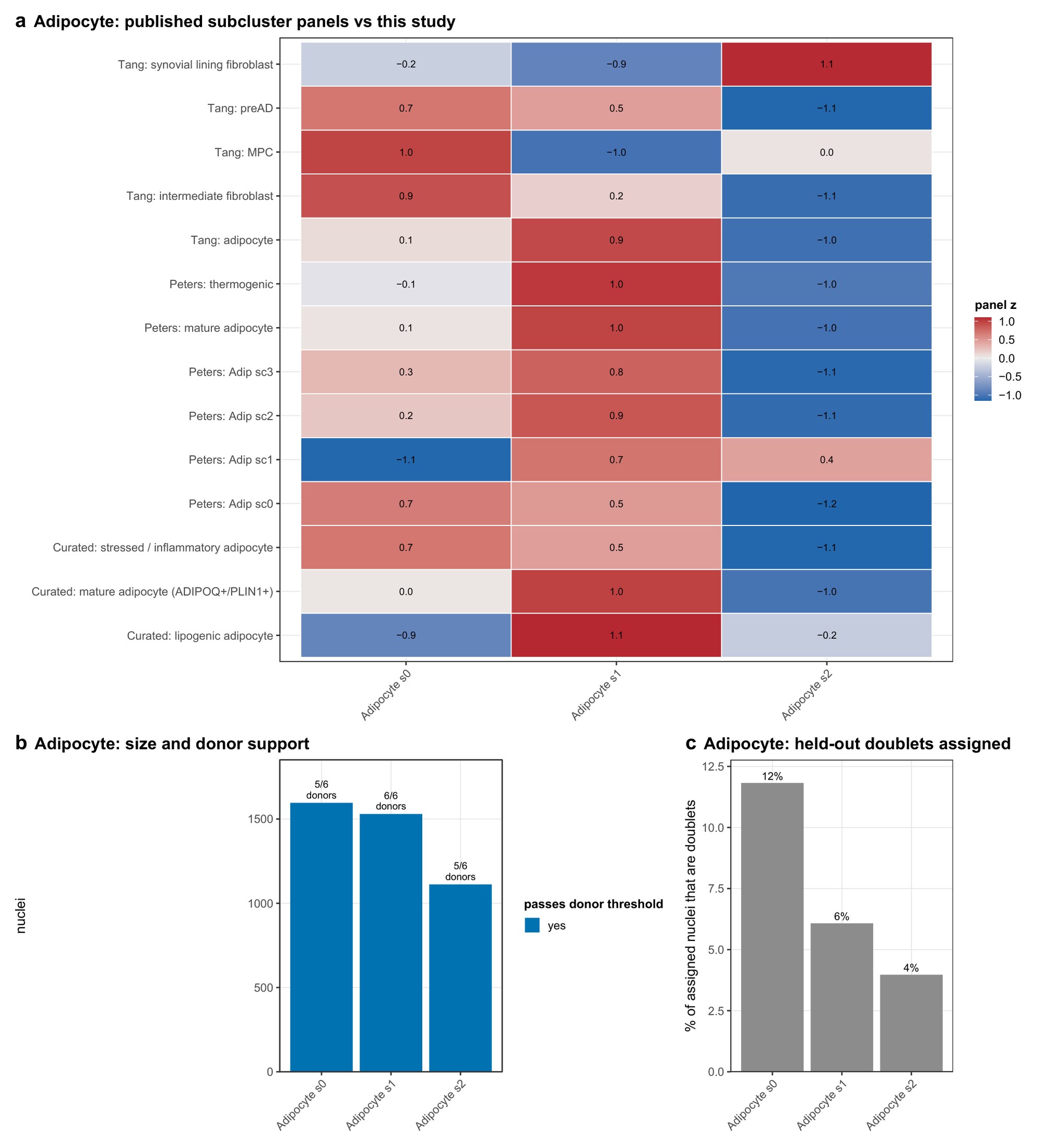

**Supplementary Fig. 34 | The three adipocyte subclusters.** **a** Published and curated adipocyte panels scored per subcluster, as the z-score of mean panel expression across the three subclusters; prefixes give the source (Tang, Tang et al.; Peters, Peters et al.; Curated, marker panels curated for this study). The external mature-adipocyte panels score highest on the lipogenic subtype, except Peters Adip sc0, which scores highest on the secretory subtype, so this split was defined in this study and does not reproduce a published one. **b** Subcluster size (singlet nuclei), with the number of donors contributing at least 25 nuclei; color marks whether a subcluster passes the donor threshold, and all three do. **c** Share of the nuclei assigned to each subcluster that are held-out predicted doublets, projected by nearest-neighbor vote. Two subclusters carry biological names; the third (Adipocyte s2) is named for its low library complexity because it has no program of its own (Supplementary Fig. 24). Adipocyte s0, secretory (CFD+/C1QTNF7+); s1, lipogenic (GPAM+/DGAT2+); s2, low-complexity.

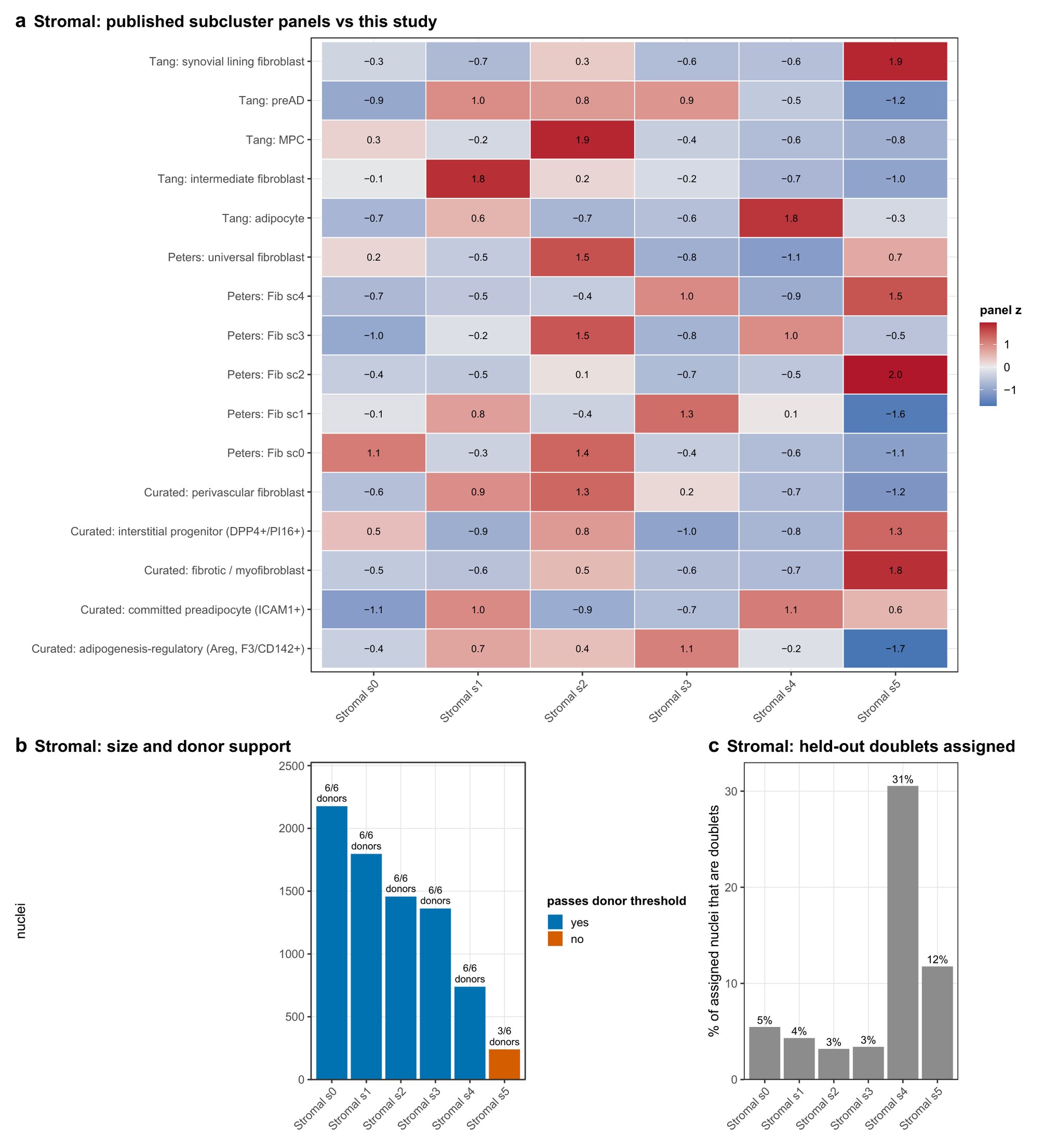

**Supplementary Fig. 35 | The six fibroblast subclusters.** Panels as Supplementary Fig. 34. The two stromal labels of the reference were pooled before splitting, so these six are not a refinement of one label each; three derive from each side. The lining-like subcluster (Stromal s5) fails the donor threshold (3 of 6 donors), and the lipid-laden subcluster (Stromal s4) receives 31% of its assigned nuclei as held-out doublets. The adhesion subcluster (Stromal s0) is the largest and scores highest on none of the published panels, so it is a candidate population without a published counterpart. Panel names containing the words progenitor or preadipocyte refer to the published signature being scored and do not describe our population. Stromal s0, adhesion (ITGA11+/SULF1+); s1, SMOC1+/GHR+; s2, matrix-high (COMP+/CILP+); s3, CXCL14+/IGFBP7+; s4, lipid-laden (CIDEA+/CD36+); s5, lining-like (CLIC5+/HAS1+).

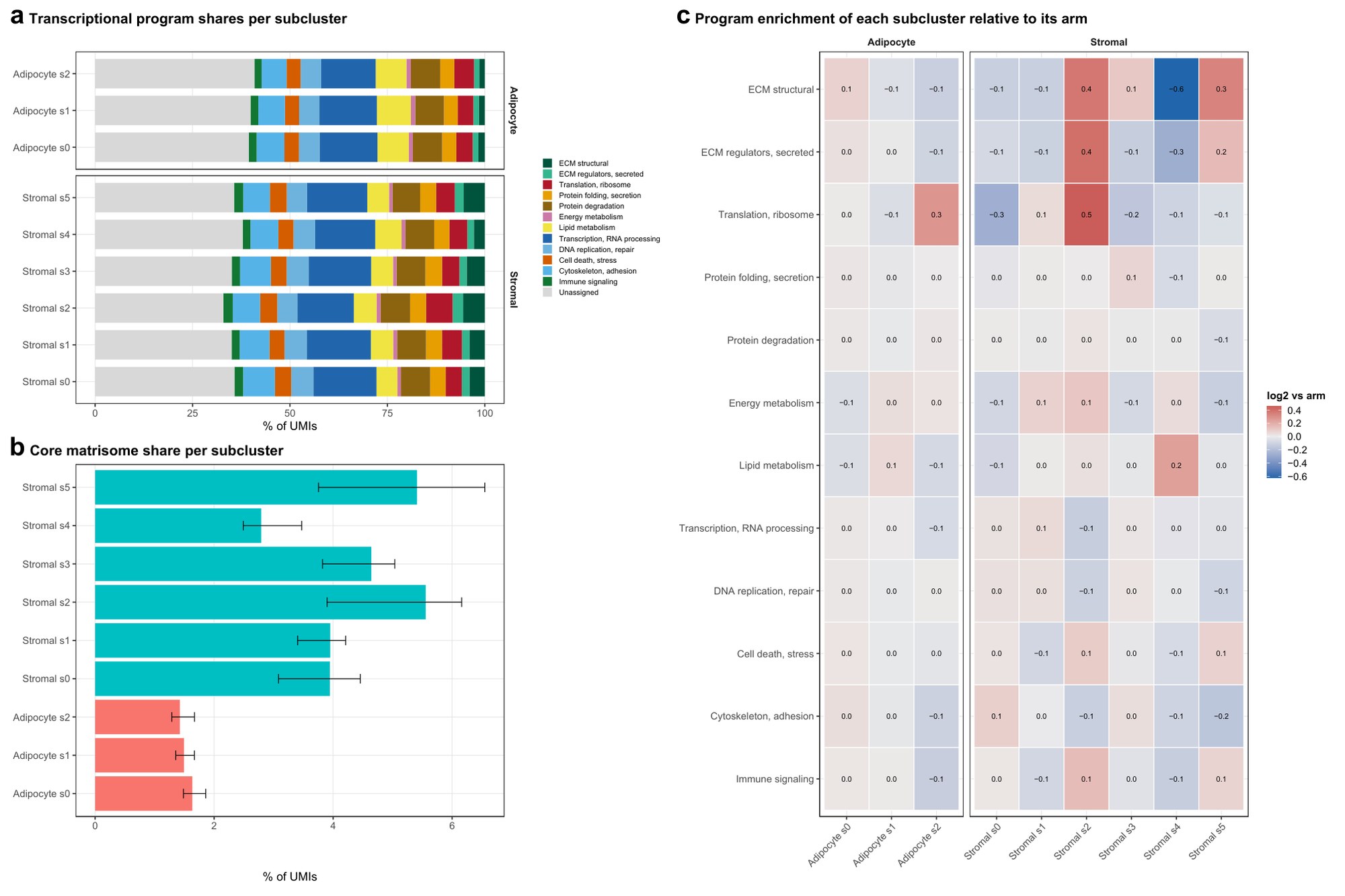

**Supplementary Fig. 36 | Transcriptional program shares of the IPFP subclusters.** **a** The twelve-program partition per subcluster, as Fig. 4c. **b** Core matrisome share of counts; bars, pooled share; error bars, mean ± SD across donor pseudobulks. **c** Enrichment of each program in each subcluster relative to the arm. These are the measurements behind the descriptors in the names: matrix-high carries the highest core matrisome of any IPFP population, lipogenic the highest lipid-metabolism share in the tissue, lipid-laden the highest in the stroma together with the lowest matrisome, and adhesion the highest cytoskeleton and adhesion share. Two subtypes received no descriptor (their strongest within-arm program enrichment was log2 0.08 and 0.11) and carry marker-only names.

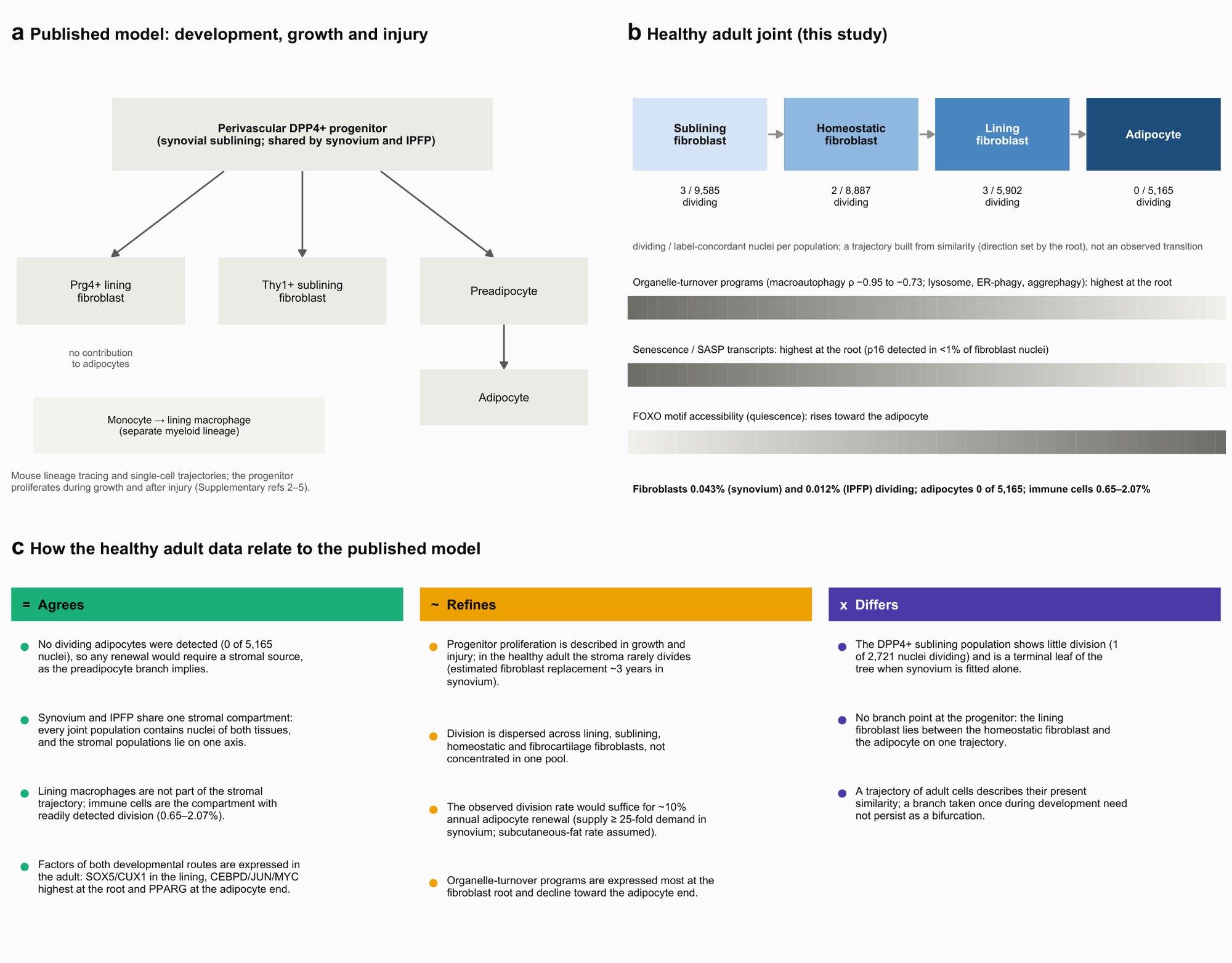

**Supplementary Fig. 37 | How the healthy adult joint relates to the published lineage model.** **a** The model assembled from mouse lineage tracing and single-cell trajectories: a perivascular DPP4+ progenitor in the synovial sublining gives rise to lining and sublining fibroblasts and, through a preadipocyte, to IPFP adipocytes;^4^ lining fibroblasts also derive from Gdf5-lineage progenitors (not drawn);^2,3^ lining macrophages derive from a separate myeloid lineage.^5^ **b** The healthy adult joint in this study: the four populations along the stromal trajectory with their dividing nuclei over label-concordant nuclei, and the programs that change along it. The trajectory is built from similarity; its direction is set by the root. **c** Where the healthy data agree with, refine, or differ from the model. Every number is read from the analysis tables at rendering.

**Supplementary References**

1. Tang, S. et al. Single-cell atlas of human infrapatellar fat pad and synovium implicates APOE signaling in osteoarthritis pathology. *Sci. Transl. Med.* **16**, eadf4590 (2024).

2. Roelofs, A. J. et al. Joint morphogenetic cells in the adult mammalian synovium. *Nat. Commun.* **8**, 15040 (2017).

3. Collins, F. L. et al. Taxonomy of fibroblasts and progenitors in the synovial joint at single-cell resolution. *Ann. Rheum. Dis.* **82**, 428–437 (2023).

4. Li, J. et al. Synovium and infrapatellar fat pad share common mesenchymal progenitors and undergo coordinated changes in osteoarthritis. *J. Bone Miner. Res.* **39**, 161–176 (2024).

5. Culemann, S. et al. Locally renewing resident synovial macrophages provide a protective barrier for the joint. *Nature* **572**, 670–675 (2019).

**Supplementary Data**

Supplementary Data 1. Function of the 200 most abundant protein-coding transcripts in each tissue and of the tissue-distinct part of each list (Hallmark and GO Biological Process over-representation).

Supplementary Data 2. Bulk versus single-nucleus gene detection over the 36,366 genes annotated in both references, at 1, 5 and 10 TPM or CPM.

Supplementary Data 3. The 29 opposing gene pairs: program, driver and blocker, mechanistic class, supporting reference (PMID), per-tissue direction and robustness checks.

Supplementary Data 4. Mapping of the per-tissue populations onto the 16 populations of the joint synovium and IPFP reference.

Supplementary Data 5. The 20 healthy donors: age, sex, SPARC subject identifier and the assays (bulk RNA-seq, single-nucleus multiome) for which each donor contributed synovium and IPFP; D1–D6 are the multiome donor labels used in the figures.

Supplementary Data 6. The 26 cell-turnover programs: MSigDB definitions, gene coverage and trends along each pseudotime trajectory.
